# Plant longevity, drought and island isolation favoured rampant evolutionary transitions towards insular woodiness

**DOI:** 10.1101/2022.01.22.477210

**Authors:** Alexander Zizka, Renske E. Onstein, Roberto Rozzi, Patrick Weigelt, Holger Kreft, Manuel J. Steinbauer, Helge Bruelheide, Frederic Lens

## Abstract

Insular woodiness (IW)—the evolutionary transition from herbaceousness towards woodiness on islands—is one of the most iconic features of island floras. Since pioneering work by Darwin and Wallace, five IW drivers have been proposed: (i) favourable aseasonal climate and (ii) lack of large native herbivores promote plant longevity that (iii) results in prolonged flowering favouring outcrossing. Alternatively, (iv) competition for sunlight requires taller and stronger stems, and (v) drought favours woodiness to safeguard root-to-shoot water transport. However, information on the occurrence of IW is fragmented, hampering tests of these potential drivers. Here, we identify 1,097 insular woody species (IWS) on 375 islands, and infer at least 175 evolutionary transitions on 31 archipelagos, concentrated in six angiosperm families. Structural equation models reveal that the IWS richness on oceanic islands correlates with aseasonal favourable climate, followed by increased drought and island isolation (approximating competition). When continental islands are included, reduced herbivory pressure by large native mammals, increased drought and island isolation are most relevant. The repeated evolution of IW opens promising avenues to disentangle the variation in gene regulatory networks triggering wood formation, and emphasize individual archipelagos as laboratories of evolution, where similar environmental conditions replicate convergent evolution of similar traits.

## Introduction

The repeated evolution of peculiar morphological, physiological, and behavioural traits in insular lineages is described in island syndromes^1–4^. Iconic examples in the animal kingdom include flightlessness in birds and insects^5,6^, naivety toward predators^7^, and changes in body size^8,9^. In angiosperms, the evolution of herbaceousness towards insular woodiness (IW) is one of the most prominent aspects of island floras^4,10–12^. Famous examples include the Hawaiian silverswords (*Dubautia, Argyroxiphium*) and woody violets (*Viola*) as well as the Macaronesian tree lettuces (*Sonchus*) and viper bugglosses (*Echium)*. Interestingly, angiosperms evolved from a woody ancestor, making them ancestrally woody and invoking that non-woody (herbaceous) angiosperms lost their woodiness during evolutionary history^13^. Therefore, IW represents a phylogenetically derived state that is treated as an evolutionary reversion^14^.

In contrast to well-documented island syndromes in animals, IW and its evolutionary drivers remain poorly understood^10,11^. IW may be induced by a longer life span via three potential drivers (Table 1): (i) *more favourable aseasonal climate* buffered by the surrounding oceans leading to continuous growth that is not interrupted by frost^11^, (ii) *reduced herbivory* due to a lack of large native island herbivores and hence no grazing of small short-lived herbaceous species^11^, and (iii) advantages of longer flowering time in an insect-poor island environment *promote cross pollination*^15,16^. Alternative IW hypotheses point to (iv) *biological competition* among colonizing herbs favouring taller stems to capture more sunlight^17,18^, and (v) *increased drought stress* demanding a better protection of root-to-shoot water transport against hydraulic dysfunction^19–21^ (Table 1). Testing these five hypotheses at the global scale has so far been hampered by the fragmented knowledge about IW, missing information on evolutionary relationships, and a lack of standardized descriptions of island environments. For instance, existing IW studies have focused on a few iconic island clades^22,23^ or a single archipelago^12,24^, leaving most insular woody clades across the world unidentified. Therefore, documenting and understanding IW remains at the forefront of island biology^25^, despite pioneering work of Darwin, Wallace, Hooker and others^16,17,26^.

**Table 1.** Existing hypotheses for the evolution of IW, along with resulting predictions and the outcome of the analyses in this study. A fifth hypothesis on the role of pollinators and longevity was not tested in this study due to lack of a global island pollinator dataset. The results of the analyses concerning each prediction are shown in bold in the last column, and relate to a model including only oceanic islands. *indicate instances in which the results from a model including oceanic and continental islands disagree (Fig. 4).

| Hypothesis | Argumentation | Predictions tested in this study |
| --- | --- | --- |
| *Competition* | When isolated oceanic islands emerge, they are first colonized by herbaceous lineages that generally have better long-distance dispersal abilities compared to tree lineages. In this initial vegetation without trees, competition for light among dense herbaceous populations selects for taller plants, which need to mechanically reinforce their stems, leading to wood formation<sup>17</sup>. This idea is extended by<sup>18</sup> who argued that natural selection will favour woodiness if pioneer immigrants of open habitats invade denser habitats. | 1. The number of IWS is higher on more isolated oceanic islands (**supported**).<br>2. Evolutionary transitions towards IW are more common on more isolated oceanic archipelagos (**rejected**). |
| *Drought* | Species with increased woody stems are more drought tolerant than their herbaceous counterparts, and often grow in (seasonally) dry habitats<sup>12,20</sup>. The more lignified vessel cell walls and its fine-scale adaptations in IW stems are able to avoid the formation and spread of spontaneous drought-induced air bubbles that can block the long-distance water transport from roots to leaves<sup>20</sup>. This air blockage is one of the prime mechanisms leading to drought-induced plant mortality<sup>19</sup>, and points to drought as a potential driver of IW<sup>21</sup>. | 1. The number of IWS is positively correlated with drought expressed by precipitation seasonality (**supported**), less precipitation in the warmest quarter of the year (**supported**) and by a lower aridity index (**rejected\***).<br>2. IWS occur predominantly in open rather than forest habitats on individual archipelagos (**rejected**, since habitat importance varies with plant family, Fig. 1). |
| *Favourable aseasonal climate* | A favourable aseasonal climate (without frost), caused by the buffering effect of the surrounding ocean, leads to a continuous growth period throughout the year. This continuous growth period leads to increased plant age which in turn favours woodiness in stems<sup>11</sup>. | 1. The number of IWS is negatively correlated with number of frost days (**supported\***), higher past climate change velocity in temperature (**rejected**) and higher past climate change velocity in precipitation (**supported\***), and higher temperature seasonality (**rejected\***).<br>2. Evolutionary transitions towards IW are more common on islands in lower latitudes characterized by a more stable favourable paleoclimate (**rejected**). |
| *Reduced Herbivory* | The absence of native large mammal herbivores on islands allows short-lived herbaceous species to live longer, inducing evolution of woody shrubs<sup>11</sup>. | 1. The number of IWS is positively correlated with the time-integrated species richness of large mammal herbivores richness **(rejected\*)**. |

To reveal the global drivers of IW, we compiled a novel dataset of insular woody species (IWS) in angiosperms. We identified IWS, and inferred the timing and number of evolutionary transitions to IW per archipelago and plant family based on information from hundreds of molecular phylogenies, floras and taxonomic revisions. We then combined this dataset with past and present environmental data on islands worldwide to test four of the five IW hypotheses. We specifically target three central themes with respect to IW evolution:

1. *Species identity and evolution of IW*. How many IWS are there? How many times did IW evolve and in which lineages? How clustered is IW on the angiosperm Tree of Life?
2. *Geographic distribution of IW*. Which islands and archipelagos harbour IWS and evolutionary transitions to IW? Is IW prevailing on oceanic compared to continental islands?
3. *Potential drivers of IW*. Which of the four IW hypotheses tested (competition, drought, favourable aseasonal climate, and reduced herbivory) is supported by the extant distribution of IWS worldwide? Do the IW drivers differ between all islands compared to only oceanic islands?

## Results

### Species identity and evolution of IW

We identified 1,097 IWS belonging to 149 genera and 32 families, representing at least 175 evolutionary transitions. IW was widespread across the Tree of Life of angiosperms, but concentrated in only few plant families (Fig. 1, Supplementary Fig. 1). Most IWS (898/82%) and evolutionary transitions towards IW (131/75%) occurred in the super-asterids clade, with the majority of the species in only two families: Gesneriaceae (327 species/30%) and Asteraceae (256/23%). The top five families with evolutionary transitions (91/52%) were: Asteraceae (47/27%, super-asterids), Amaranthaceae (15/9%, super-asterids), Brassicaceae (12/7%, super-rosids), Rubiaceae (10/6%, super-asterids), and Campanulaceae (7/4%, super-asterids) (See Supplementary Information Section 2 for the number of IWS and minimum number of IW shifts per genus). In Gesneriaceae, a low number of transitions lead to at least 245 IWS due to the spectacular radiation of the genus *Cyrtandra*. Stem ages for 61 insular woody clades with time-calibrated phylogenies available varied from 19.7 Ma to 0.1 Ma before present, with 51 of the lineages dating back less than 10 Ma; many of these recent IW clades were endemic to the Canary Islands (Extended Data Fig. 1).

**Figure 1.**
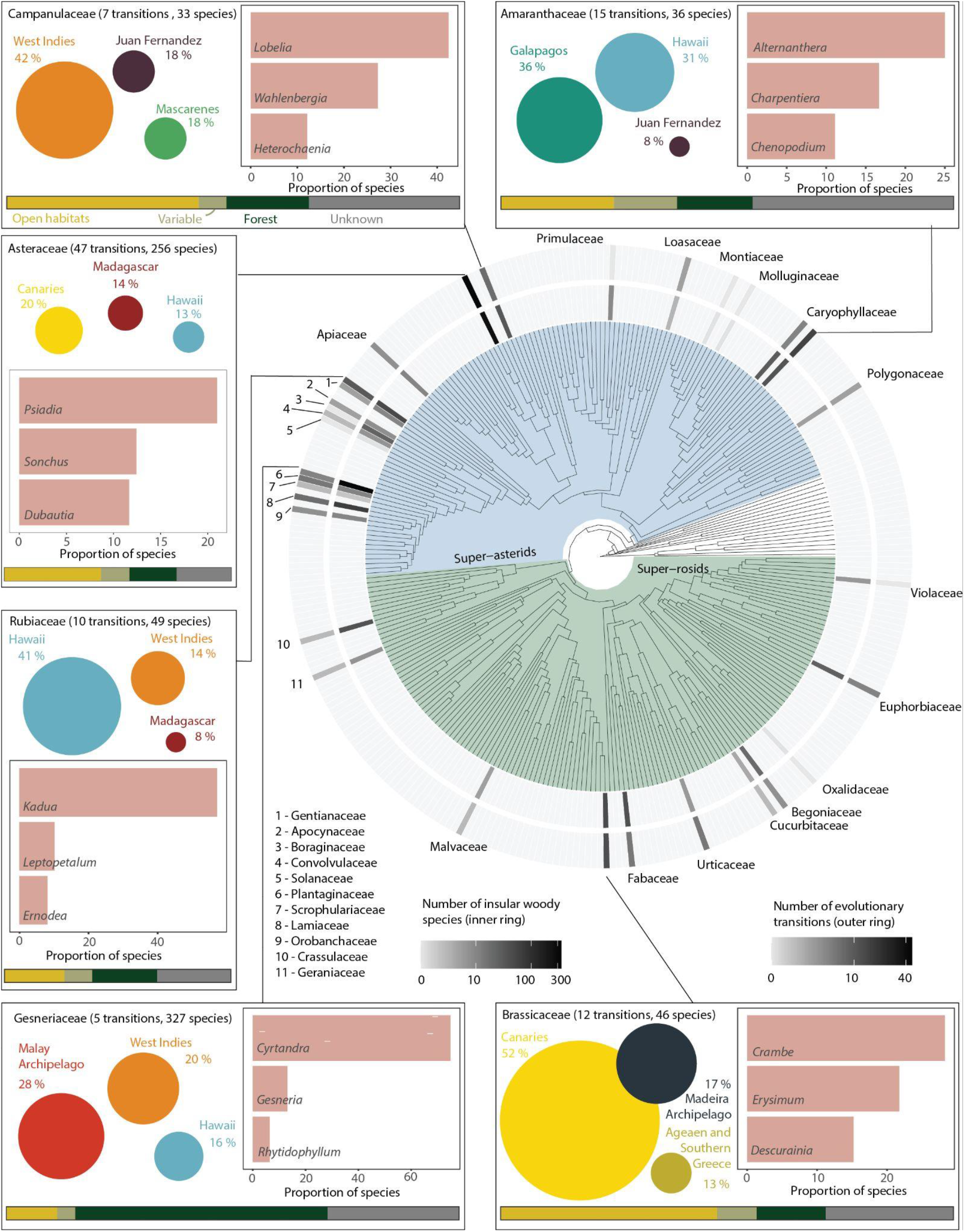
Insular woodiness across the angiosperm tree of life. The phylogenetic tree shows the number of insular woody species (IWS, inner ring) and the minimum number of evolutionary transitions in each family (outer ring). The inlets show additional information for the families with the highest number of species and transitions: the bubbles show the three archipelagos comprising the highest proportion of IWS in the family (bubbles are to scale across families). The bar chart shows the proportion of IWS in the three genera with most IWS in each family. The stacked bar chart at the bottom shows the proportion of IWS occurring in open habitats, forest, and forest and open habitats (variable); grey shows the proportion of species for which no habitat information was available.

We found mixed results concerning the phylogenetic clustering of IW. At the family level, the mean pairwise distance (MPD) suggested phylogenetic clustering of the occurrence of IWS compared to a null model (MPD_observed_ = 222.2, MPD_null_ = 237.2, Standardized effect size = −3.34, p = 0.01). Additionally, at the genus level, Pagel’s lambda (ƛ) indicated weak but significant phylogenetic clustering in the proportion of IWS per genus (mean ƛ across replicates = 0.013, with p< 0.05 for difference to ƛ= 0 for all replicates). In contrast, at the family level the mean nearest taxon distance (MNTD) rejected phylogenetic clustering of the occurrence of IWS compared to a null model (MNTD_observed_ = 136.9, MNTD_null_ = 152.1, Standardized effect size = −1.499, p = 0.08). Additionally, at the genus level, Blomberg’s K indicated no phylogenetic signal in the proportion of IWS per genus (mean K across replicates = 0.03, p> 0.05 for all replicates). These mixed results reflect that IWS and evolutionary transitions towards IW were concentrated in specific families, predominantly in the super-asterids, yet at the same time IW evolved repeatedly in distantly related groups across the angiosperm phylogeny (Fig. 1).

### Geographic distribution of IW

IWS occurred on islands worldwide, but generally, the number and relative representation of IWS was low on islands outside the (sub)tropics (Fig. 2). Of the 175 evolutionary shifts towards IW, we could confidently assign 162 to 31 individual archipelagos (Fig. 3). Globally, the Canary Islands (204 IWS representing 18% of all IWS and at least 33 evolutionary transitions to IW equal to 19% of all transitions) and the Hawaiian archipelago (199/18% species, 17/10%) emerged as centres of IW, followed by the Malay archipelago (129/11% species, 12/7% transitions), and the West Indies (98/9% species, 11/6% transitions). Additional oceanic archipelagos with a high number of IWS and evolutionary transitions were Madeira (36/3% species, 6/3% transitions), Juan Fernandez Islands (35/3% species, 8/5% transitions), and Mascarenes (34/3% species, 3/2% transitions; Supplementary Information Section 3). The top three individual islands with the highest number of IWS were Tenerife (97 species, Canary Islands), Kaua’i (79, Hawaiian archipelago) and Madagascar (78). Of the 10 single islands with the most IWS, eight belonged to the Canary Islands or the Hawaiian archipelago. The islands with the highest proportion of IWS on the total angiosperm flora were Santa Fé (25%, Galápagos), Robinson Crusoe Island (25%, Juan Fernández), and Kaho’olawe Island (21%, Hawaiian archipelago). The continental island with the highest proportion of IWS was the South Island of New Zealand (3.2%, rank 61 of all islands). The proportion of IWS varied significantly within archipelagos, in particular for the two archipelagos with most IWS: from 21% (Kaho’olawe) to 9% (Lana’i) for the Hawaiian archipelago, and from 16% (Tenerife) to 7% (Lanzarote) for the Canary Islands (Fig. 2, Extended Data Fig. 2).

**Figure 2.**
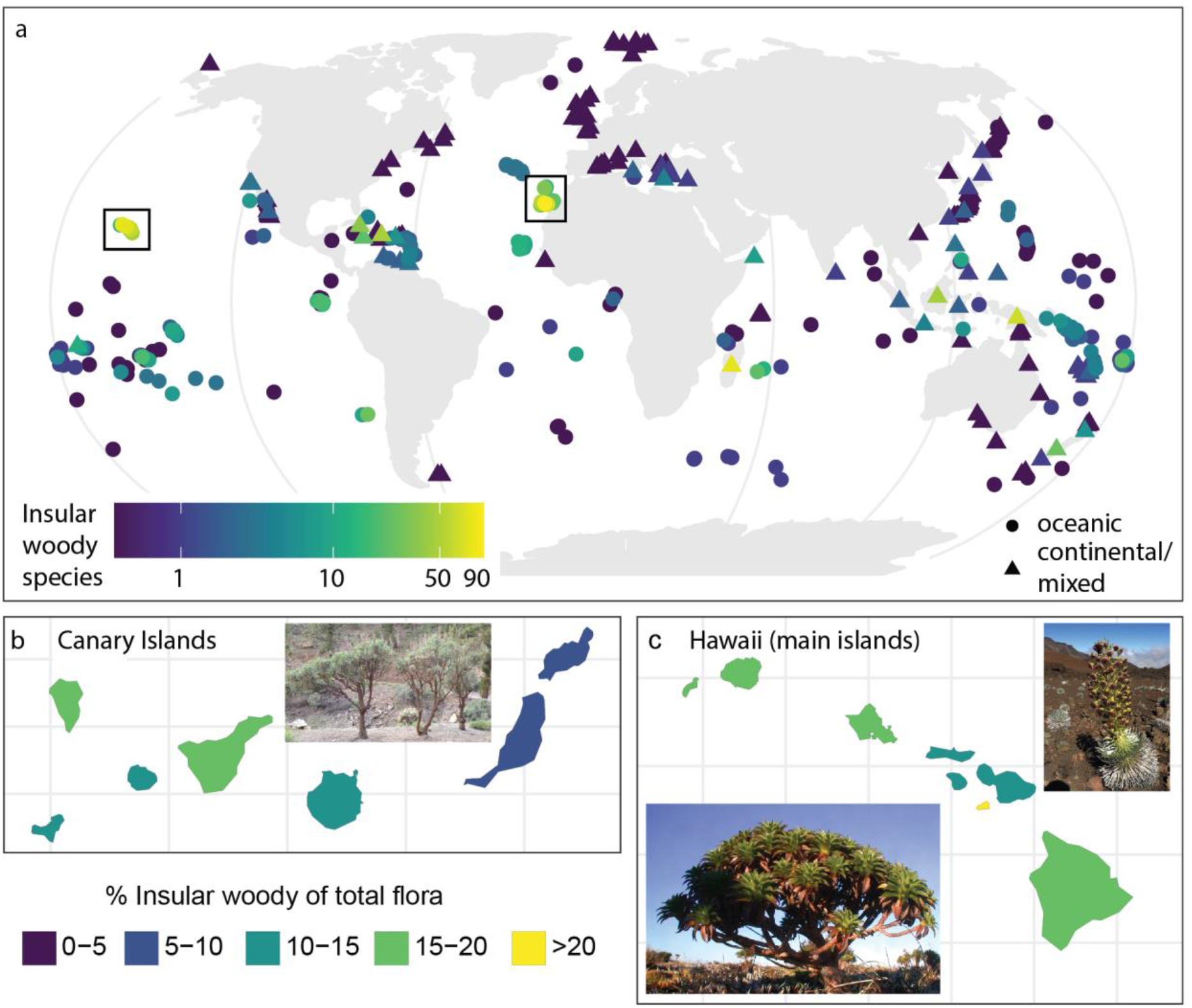
Global geographic distribution of IW at the level of islands. **a)** The numbern of IWS across all islands. **b,c)** Proportion of IWS of the total flora on islands of the two archipelagos with most IWS, the Canary Islands and Hawaii. The inlet pictures show three iconic examples of IWS: *Echium virescens* on the Canaries (picture F. Lens), *Argyroxiphium sandwicense*, and *Dubautia waialealae* on Hawaii (silverswords, pictures by Seana Walsh and Ken Wood, courtesy of National Tropical Botanical Garden, Hawaii).

**Figure 3.**
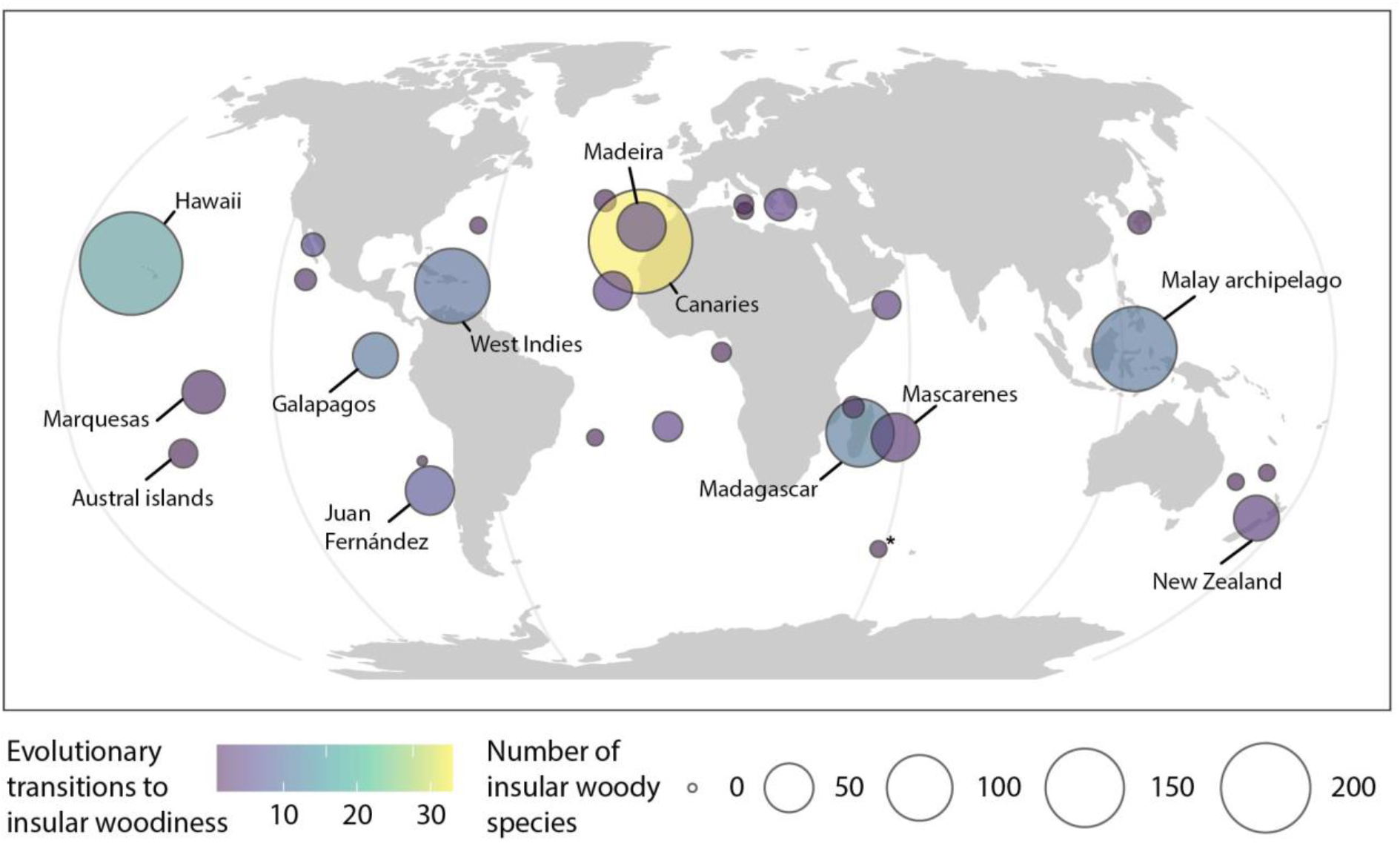
Minimum number of evolutionary shifts to IW and number of IWS on archipelagos worldwide. Only archipelagos with at least one evolutionary shift are shown for clarity. An additional 13 shifts could not be linked unambiguously to any of the archipelagos. The * summarizes multiple Southern Indian Ocean islands (Kerguelen, Crozet, Prince Edward Islands and Heard & MacDonald).

### Potential drivers of IW

On oceanic islands, we found direct correlations of the number of IWS with explanatory variables related to the *competition, drought*, and *favourable aseasonal climate* hypotheses after statistically accounting for the differences in island area, mean elevation and total number of angiosperm species, (Table 1, Fig. 4, Extended Data Figs. 3&4). As expected, we found direct, negative effects of precipitation change velocity since the last glacial maximum, precipitation of the warmest quarter, and the number of frost days; and positive effects of precipitation seasonality and distance to the nearest continent (Table 1, Fig. 4a). When including continental islands, the importance of the explanatory variables changed considerably (Fig. 4b). Most notably, the direct effects of the explanatory variables related to *reduced herbivory* were most important, followed by variables related to *drought*, *competition* (approximated by isolation, the distance to nearest continent), and *favourable aseasonal climate*. As expected under the individual hypotheses, we found positive effects of distance to the nearest continent and precipitation seasonality and negative effects of time-integrated mammal herbivore species richness, the aridity index (lower aridity index indicating drier climate), precipitation of the warmest quarter and temperature seasonality on the number of IWS species per island (Fig. 4b). Additionally, for both models, we found indirect effects of explanatory variables related to all hypotheses via the total angiosperm richness and the time-integrated species richness of large mammal herbivores (Fig. 4). The results were qualitatively similar when excluding the Hawaiian archipelago and the Canary Islands as outliers. Overall, the model fit was high for all SEMs and best for oceanic islands alone (incl. Canary Islands and the Hawaiian archipelago; R^2^_oceanic_ = 0.61 *v*. R^2^_all_ = 0.55; Extended Data Fig. 4).

**Figure 4.**
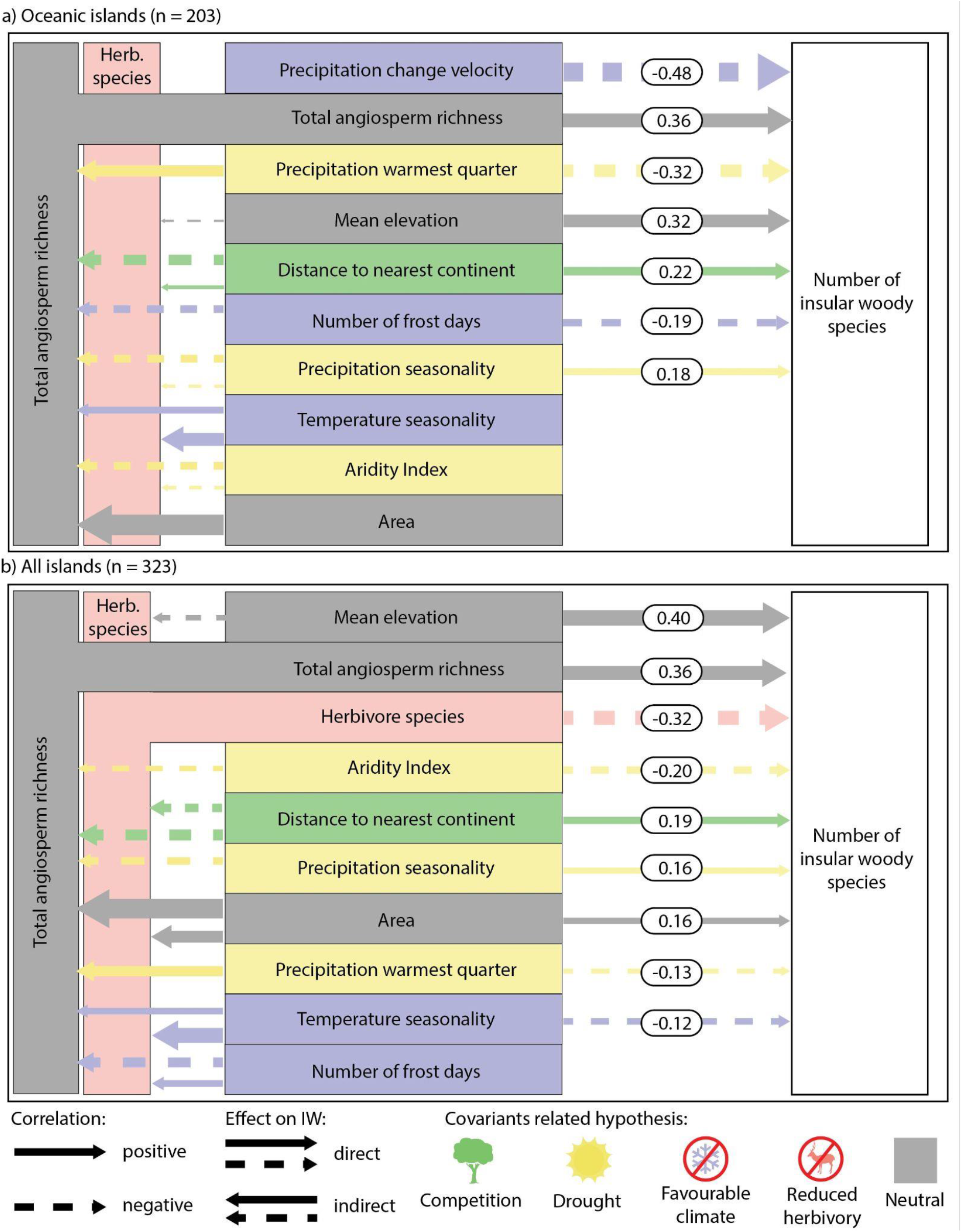
Environmental explanatory variables of IW. **a)** oceanic islands, **b)** oceanic and continental islands. The explanatory variables are ordered according to the direct effect size from the top to the bottom. The numbers show standardized effect sizes of direct effects; thickness of arrows is proportional to effect size; colours link explanatory variables with hypotheses for the evolution of IW. Herbivore species = time-integrated richness of large mammal herbivores. See Table 1 for details on the hypotheses, Table S1 for details on the individual explanatory variables and Extended Data Fig. 4 for all effect sizes. Only predictors with at least one significant direct or indirect effect are shown. The deer silhouette by Birgit Lang in the Public domain from www.phylopic.org.

Concerning the number of evolutionary transitions to IW at the archipelago level, we found a strong and significant positive effect of archipelago area on the number of transitions (but not on the presence or absence of any shifts, i.e. in the count but not the zero hurdle components of the model). In contrast, we did not find a significant effect of distance to nearest continent, vicinity to the equator, or minimum archipelago age (Extended Data Figs. 5&6), neither for the number nor the occurrence of evolutionary transitions. These results contradicted the predictions from the *competition hypothesis* (more transitions on more isolated islands) and the *favourable climate hypothesis* (more transitions on islands closer to the equator with more favourable climate on evolutionary time scales).

## Discussion

Here, we provide a comprehensive global synthesis of the taxonomy, geography and evolution of IW, and use global scale data and a correlative approach to test insular woodiness (IW) hypotheses originally postulated based on data from individual islands and archipelagos. We identified at least 175 evolutionary transitions from herbaceousness towards IW across angiosperms, giving rise to over 1,000 insular woody species (IWS). This more than triples the known number of IWS and transitions^e.g., 23,24,27^. A large majority of these IWS are nested in the super-asterids clade (super-asterids clade, particularly in the families Gesneriaceae and Asteraceae; Fig. 1), and mainly belong to relatively young lineages that originated less than 10 Ma ago (Extended Data Fig. 1). Our results show IW as a widespread phenomenon, and reveal oceanic (i.e. typically volcanic) islands in the Mediterranean and (sub)tropical climatic belt as centres of IWS richness (Fig. 2) and as main locations for evolutionary transitions towards IW (Fig. 3). Among oceanic islands, the Canary and Hawaiian Islands (excluding the large lobelioid radiation with an unclear origin of woodiness) prevail in terms of number of IWS (204 vs 199) as well as transitions (17 vs 33)^12,23,28^ (Supplementary Information Section 3). Among continental islands, the flora of Madagascar stands out (78 IWS, 13 transitions). New Caledonia, in contrast, harbours surprisingly few IWS and evolutionary transitions (3/0), despite its tropical climate and the total submergence of the island during the Late Cretaceous and Early Paleogene^29^ (resulting in the chance of herbaceous populations to colonize the island after re-emergence and evolve IW as response to competition). The occurrence of 395 IWS on continental islands as well as the 808 derived woody island species that evolved their woodiness on nearby continents invoke the question how common evolutionary transitions towards woodiness have occurred on continents. For instance, we consider the Hawaiian lobelioid clade (Campanulaceae)—the most conspicuous radiation of (woody) species on the archipelago with 126 species—provisionally as derived woody (woodiness evolved on mainland instead of insular woody) until molecular phylogenies provide more insight into the habit of the ancestral lineage that colonized the archipelago^30^. A prime example of a woody island radiation that developed its woodiness on adjacent continents is *Veronica* (Plantaginaceae) with 118 derived woody island species, mainly native to New Zealand (former genera *Hebe* and *Parahebe*).

We used IW species richness and its correlation with island environmental characteristics to test four hypotheses that explore why plants evolved their woodiness on islands. The direct (and most indirect) effects observed in our structural equation models (SEMs) indicate that variables that prolong plant longevity—due to either favourable aseasonal climatic conditions and perhaps also reduced herbivory^11^—as well as variables linked with drought^20,21^ and competition^17,18^ (approximated by island isolation) likely acted as global drivers of IW (Fig. 4, Table 1). In terms of the number of evolutionary transitions per archipelago, only archipelago area showed a strong and significant positive effect on the number—but not on presence/absence—of transitions. This effect of archipelago area may be related to the higher number of plant lineages on larger archipelagos^31^, and the resulting higher chance for transitions even if the transition rates per se are independent from archipelago size. Our results contradicted the predictions from both the *competition hypothesis* and the *favourable climate hypothesis* (Extended Data Figs. 5&6). This suggest that these IW drivers may be of lower importance for evolutionary transitions as compared to IWS distribution. One possible explanation is that the available niche space for IWS is filled by the radiation of few IW lineages rather than repeated evolutionary transitions in distantly related lineages. Our results, in combination with future improved approximations of climate and competition on evolutionary timescales, will help to better disentangle the drivers of evolutionary transitions towards IW.

When comparing the importance of the explanatory variables in the SEMs including oceanic islands vs. all islands (Fig. 4), some additional differences with respect to potential IW drivers apply specifically to oceanic islands. For instance, the reduced importance of island area and time-integrated richness of large mammal herbivores on oceanic islands is likely caused by their small area and rare occurrence of native mammal herbivores. In contrast, precipitation change velocity since the last glacial maximum and the number of frost days increase in importance, suggesting that long-term climate stability in the more isolated oceanic islands is a more relevant driver of IW compared to continental islands that are often closer to continents (Fig. 4, Supplementary Fig. 2). Our results indicate that the mechanisms behind wood formation on islands are complex, including involvement of multiple variables relating to plant longevity (i.e., stable climate and reduced herbivore pressure) as well as drought and island isolation. Although the global IW drivers are similar when comparing only oceanic vs all islands at a global scale, the type of island does affect the extent to which these variables act on wood formation (Fig. 4), possibly due to higher speciation rates on oceanic islands.

At a finer scale, the mechanisms driving IW likely become even more complex, as different environmental conditions within and across archipelagos may have driven IW. For instance, in addition to the different age^32,33^ and isolation of the Canary Islands vs the Hawaiian archipelago (100 vs 3,200 km to the nearest continent, respectively), the majority of IWS on the Canary Islands are native to dry open habitats that receive less than 500 mm precipitation per year^12^. In contrast, a substantial proportion of IWS on the Hawaiian archipelago is thriving in extremely wet rainforests (up to 10,000 mm mean annual precipitation; Fig. 1)^34^. Moreover, drivers can be taxon-specific, since the habitat preferences of IWS may be unique for each family (Fig. 1). Additional complexity arises from the intersection of different environmental drivers and IW hypotheses, since isolation for instance may affect competition as well as herbivore pressure. One may assume that more detailed information about current-day distribution patterns of IWS leads to a precise estimation of these finer-scale drivers, but this may not be necessarily true. The geologically dynamic nature of the islands, the presence of contrasting habitats on a single island, and the uncertainties involved in any dating study (Extended Data Fig. 2) are important bottlenecks for accurately assessing in which habitat and paleoclimate insular woody lineages have evolved^24,35^.

In addition to the environmental conditions potentially favouring the evolution to IW on a specific archipelago, intrinsic aspects of the herbaceous colonizing population are important to determine whether this lineage has the potential to develop IW^36^. For instance, monocots never produce wood^37,38^, implying that IW is per definition not possible. Furthermore, herbaceous colonisers belonging to non-monocot angiosperm lineages that are pre-adapted to radiate into the available island niches and that have already undergone multiple IW shifts elsewhere in the world (e.g., *Euphorbia, Begonia, Bidens, Lobelia, Sonchus*; See Supplementary Information Section 2) will be more likely to evolve into multiple IWS compared to herbaceous colonisers without pre-adapted traits. Interestingly, a high number of IW transitions per genus does not necessarily induce radiation events on different archipelagos, as exemplified by *Plantago* (7 IWS, 5 transitions), *Urtica* (5 IWS, 5 transitions), and *Salvia* (3 IWS, 3 transitions). Likewise, 77 out of the 149 genera (51%) identified in our IW study only include a single IWS (38%) or two IWS (13%) descending from the same colonisation event (Supplementary Information Section 2). This contrasts with the idea of IW as a key innovation that links higher diversity of plant growth forms and the opportunity to occupy greater phenotypic trait space with accelerated rates of species diversification in comparison to herbaceous continental relatives^23^. Our observation that some insular woody lineages do not diversify whereas others undergo spectacular radiations (e.g., *Cyrtandra* with 245 species^27^) show that other traits than IW must play an important role in island diversification. Identifying traits that promote diversification remains one of the unresolved key questions in evolutionary biology^25,39^, but there is growing evidence that hybridization—which can be regarded as the most extreme form of outcrossing and may overcome loss of genetic variation via founder effects in the colonizing population—is a promising candidate^36,40^.

In summary, our global synthesis of the phylogenetic and geographical distribution of IW fills a gap in documenting and understanding global patterns of rampant reversions from herbaceousness towards woodiness on islands across the world, particularly in the Mediterranean and (sub)tropical climatic belt. We show that multiple drivers, associated with variables increasing drought, island isolation, and plant longevity as result of more favourable aseasonal climatic conditions and/or reduced herbivory, affect the distribution and evolution of IW. The unexpected high number of evolutionary transitions towards IW in distantly related lineages confirm earlier studies in model plants suggesting that the gene regulatory mechanism(s) affecting the wood pathway(s) must be simple^41^ and potentially conserved through evolutionary time. Our results open a route forward to investigate these mechanisms^37,42^ that have shaped the more than 1,000 iconic woody island species.

## Methods

### Species identity and evolution of IW

Wood is defined as secondary xylem produced by a vascular cambium^43^. In angiosperms that are able to produce wood, i.e. non-monocot angiosperms, the exact boundary between woody and herbaceous species is hard to define because of the continuous variation in wood development in the aboveground stem among species^44,45^. We considered only woody island species that (1) produce a distinct wood cylinder in the stem extending towards the upper parts (i.e. shrubs, trees, lianas), but excluding species with only a woody stem base (woody herbs or suffrutescent species which are not woody enough according to our definition)^46^, and that (2) clearly evolved from a herbaceous ancestor on an island as inferred from the available phylogenetic literature. Our method allowed to distinguish between woody island species that evolved woodiness on the islands (IWS), other woody island species that evolved their woodiness on nearby continents and then expanded their range to islands (derived woody species), and members of lineages with only woody ancestors (ancestrally woody species).

To identify IWS worldwide, we screened molecular (when available time-calibrated) phylogenies from over 100 angiosperm families including woody and herbaceous species to assess whether the woody island species are insular woody, derived woody or ancestrally woody. We retrieved the habit and evolutionary relationships of the species in these families from 416 publications from the floristic and taxonomic literature (Supplementary Information Section 4). We ignored species with incomplete information on species evolutionary relationships, especially in the family Asteraceae where at least several dozen IWS could be added once future phylogenies allow distinguishing between IW or derived woodiness in some island clades (e.g., the iconic *Scalesia* trees on the Galapagos Islands). Based on our conservative definition of woodiness and due to the sometimes poorly resolved or insufficiently sampled phylogenies of island clades, our IW numbers represent minimum estimates. In addition to the IWS, we found 808 species which were derived woody and occurred on islands, but did not classify as insular woody, either because the evolutionary origin of woodiness could not be unambiguously placed on islands, or because they clearly evolved woodiness on a continent with a subsequent dispersal to islands. Most derived—not insular—woody species occurred on the Hawaiian archipelago (136 species), New Zealand (99), and the Malay Archipelago (87) (Extended Data Fig. 7). We relied on individual published phylogenies rather than a *de novo* supertree or supermatrix approach, because most IWS do not have enough sequence data to confidently place them in a large-scale phylogeny^24^. When in doubt about habit, we assessed growth form using herbarium specimens at Naturalis Biodiversity Center (L, U, WAG), and observed hand-made cross sections of stems at different heights. We then used the taxonomic information of the Leipzig Catalogue of Vascular Plants^47^ to obtain the accepted species names (see Supplementary Information Section 5 for additional details on the methods). Our dataset comprised (1) the identity of IWS and (2) Information on growth form, habitat preferences, and geographic distribution of IWS. The habitat preferences of IWS can inform on the drivers of IW, specifically with regard to the *drought hypothesis* predicting more IWS in open habitats (which are on average drier compared to forests within the same archipelago). We therefore recorded habitat information from free-text habitat descriptions found in the literature and classified species into forest vs. open habitat species. We considered all species occurring in both forest and any open habitat as “Variable”. Finally, our dataset includes (3) the number and age of evolutionary transitions towards IW per lineage and archipelago.

To test if IW was phylogenetically clustered on the angiosperm Tree of Life, we calculated the phylogenetic signal in the proportion of IWS per taxon using Blomberg’s K^48^ and Pagels *λ*^49^ as implemented in phytools^50^. We calculated the phylogenetic signal for the proportion of IWS species per tip on two taxonomic levels: (1) the family level, using a family level phylogeny^51^, and (2) the genus level, using a genus-level phylogeny obtained by randomly pruning all but one species per genus from a large-scale phylogeny of seed plants^52^. We repeated the genus-level analysis 100 times to account for paraphyletic genera. Furthermore, we calculated the phylogenetic signal in the presence of IWS (binary: yes/no) across plant families (as a binary trait) using the mean pairwise distance (MPD) and mean nearest taxon distance (MNTD)^53^.

### Geographic distribution of IW

To identify global hot- and cold-spots of IW, we obtained information on islands location (latitude and longitude of the island centroid), area, mean elevation, geologic origin, and archipelago affiliation via the Global Inventory of Floras and Traits database GIFT^54^. Additionally, we obtained the total number of native angiosperm species per island from GIFT when available. We classified islands based on their geological origin into oceanic (including volcanic islands, atolls, raised ocean floor), continental (shelf islands and continental fragments), and any combination of oceanic and continental origin as “mixed”. For 83 islands where no information on the geological origin was available from GIFT, we filled this information from publicly available sources. We then summarized the number of unambiguous independent evolutionary transitions towards IW as well as the number of IWS per archipelago based on the consulted phylogenetic papers and following the archipelago scheme of GIFT. To synthesize the geographic distribution of IWS, we summarized the number of IWS per island and archipelago (Supplementary Information Section 3). Because the Hawaiian archipelago and the Canary islands are well-studied IW hotspots, we additionally calculated the proportion of IWS across all native angiosperms on individual islands of these archipelagos to illustrate the variation of IW across islands of individual archipelagos.

For all analyses, we only included islands with a size larger than 10 km^2^ and with 20 or more angiosperm species or at least 1 known IWS to exclude very small islands with a low number of species. For structural equation models (in contrast for visualization in the figures), we only included single landmasses for which an estimate of the total angiosperm species richness was available. As a result, we obtained a dataset with 425 islands for visualization and of those 322 from 94 archipelagos as data for the structural equation models (Supplementary Figure 2, Supplementary Information Section 6).

### Potential drivers of IW

To test the four existing hypotheses on the drivers of IW (Table 1), we used structural equation models (SEMs) to relate the number of IWS per island to different island environmental characteristics. We selected as explanatory variables environmental conditions for which we postulate a direct influence on the number of IWS per island under each of the four research hypotheses (Table 1, Supplementary Table 1), and used the SEMs to identify statistical significance and to rank explanatory variables by effect size. We interpret a significant correlation of an explanatory variable with the number IWS, in the expected direction (positive or negative), as corroboration for the linked hypothesis. In addition to the direct effect of the environmental conditions on the number of IWS, we included their indirect effect via the total angiosperm richness and the time-integrated number of large mammal herbivores in the model when appropriate (Supplementary Figure 3). We obtained per-island values for all environmental variables via GIFT, and the values are originally derived from gridded products of various climate and elevation models or calculated based on the islands’ shapes^54^.

To test the *competition hypothesis*, we included the distance to the next continent obtained from GIFT, as explanatory variable in the SEM. Based on the assumption that trees disperse worse across long distance than herbs, we expected more IWS on more isolated islands, since competition among herbs for light (and hence selective pressure to evolve mechanically stronger woody stems) will only be relevant in ecosystems without or with only few ancestrally woody trees.

To test the *drought hypothesis*, we included the aridity index (Mean annual precipitation / mean annual evapotranspiration; lower values indicate drier conditions)^55,56^, the precipitation of the warmest quarter^57^ and precipitation seasonality^57^ in the SEM. Based on the assumption that IW increases the resistance of plants against drought-induced gas bubble formation inside the water conducing cells in the wood cylinder (Table 1), we expected more IWS under drier conditions. Hence, we expected more IWS on islands with a lower aridity index, more IWS on islands with more precipitation seasonality, and less IWS on islands with more precipitation in the warmest quarter (as approximation for the season with most plant growth).

To test the *favourable aseasonal climate hypothesis*, we included the average number of frost days^58^, temperature seasonality^57^, as well as temperature and precipitation change velocity since the last glacial maximum^59,60^ (the latter two calculated as the ratio between the temporal change from 21,000 years before present to today and the contemporary spatial change per island) in the SEM. We included precipitation change velocity since the last glacial maximum as linked to the favourable aseasonal climate hypothesis, rather than the drought hypothesis, since the index captures climate stability over thousands of years, rather than specific drought conditions. Based on the assumption that more stable favourable climate (particularly the absence of frost) favours an increased plant lifespan and therefore wood formation, we expected more IWS under more stable climate conditions. Hence, we expect more IWS on islands with less frost days, less temperature seasonality, and less temperature and precipitation change velocity since the last glacial maximum velocity.

To test the *reduced herbivory hypothesis*, we included the time-integrated species richness of large mammal herbivores in the SEM. We combined this index as measure of herbivore pressure from two sources: First, we obtained all terrestrial mammal species with a body mass above 1 kg and more than, or equal to 20% of plant material in their diet, and a potential natural distribution on islands via the Phylogenetic Atlas of Mammal Macroecology v1.2 (Phylacine)^61^. *Phylacine* includes mammal species that lived from the last interglacial (~130,000 years ago) until the present. We selected the diet threshold to also include species with a small percentage of vegetative plant material in their diet, since in an island setting where herbivore pressure is generally low, even species with a small fraction of plant material in their diet may have a strong impact on plant life history. Furthermore, we excluded species that exclusively eat seeds and fruits, such as for instance many rodents, jackals, badgers and foxes, which have little impact on a plant’s vegetative growth. Second, to also account for the impact of herbivory at deeper time scales we complemented the species list from Phylacine with a novel list of extinct terrestrial mammalian herbivores and omnivores from fossils recorded in scientific literature (Supplementary Information Section 1). We included selected fossil taxa with a high amount of plant material in their diet above 1 kg body mass that occurred over the past 20 Ma. We manually removed duplicate entries that arose in some cases from species present in the fossil and Phylacine data, and used this list to compile the time-integrated species richness of large native mammal herbivores per island (Supplementary Information Section 7). We included fossils to quantify the herbivore pressure throughout the evolutionary history of island plant lineages. We focused on mammal herbivore pressure, since bird herbivory has distinctly different effects on plant adaptations and because most birds on (oceanic) islands have a diet primarily based on marine animals. Based on the assumption that reduced herbivore pressure increases plant life span and therefore wood formation, we expect more IWS on islands with a lower time-integrated species richness of large mammal herbivores.

To control for the effects expected under a neutral model of IW evolution, we included the direct effect of island area, mean elevation and total angiosperm richness in the SEM. We expect more IWS on larger islands (due to lower extinction rates), on islands with a higher mean elevation (due to increased habitat heterogeneity caused by geological heterogeneity), and on islands with a higher overall angiosperm richness (due to a higher pool of evolutionary lineages and potential higher background diversification rates).

We designed our SEMs following Onstein et al.^62^: First, we normalized all variables between 0 and 1. We started with an *a priori* SEM that included all hypothesized pathways among all predictor variables (Supplementary Fig. 3) and evaluated the model’s modification indices, model fits and residual correlations among those^63^. To ensure an adequate fit of SEMs, we made sure that p-values of chi-square tests > 0.05, comparative fit index (CFI) > 0.90 and confidence intervals of the root mean square error of approximation (RMSEA) < 0.05. We progressively deleted paths with the least statistical significance from the SEM until our final SEM only consisted of significant pathways (at p < 0.05), for which we extracted the standardized coefficients. We tested the model residuals for normality and equal variances, and we checked for extreme outliers that could affect the results. Since IW has until recently mostly been discussed using volcanic islands, we fitted one SEM focusing on oceanic islands only (global oceanic island dataset with 203 islands), and a second model including all islands (i.e. also islands of continental origin; global dataset with 322 islands). Furthermore, as a sensitivity analysis to investigate drivers of IW beyond the most iconic archipelagos which potentially are outliers with regard to the number of IW species, we repeated both analyses excluding the Hawaiian archipelago and the Canary Islands (188 and 307 islands, for oceanic and all islands respectively). We fitted the SEMs using the R package ‘lavaan’ v.0.6.8^64^. Since multiple islands were missing climatic information, we used a random forest multiple imputation method to fill these gaps^65,66^. Specifically, we estimated information on temperature seasonality, precipitation seasonality and precipitation of the warmest quarter for four islands (1% of the islands included in the analysis), mean aridity index for nine islands (2.1%), mean elevation for 12 islands (2.9%), climate change velocity of temperature for 13 islands (3.1%), and climate change velocity of precipitation for 27 islands (6.5%).

Spatial autocorrelation can affect results from non-spatial analyses^67^. To assess the extent of spatial autocorrelation in our data, we used correlograms and visualized changes in Moran’s I values in the raw data response variables and in the residuals when fitting an ordinary least squares (OLS) regression model with the same set of predictor variables on IWS richness as in the SEM. Moran’s I values indicated that there was little (non-significant) spatial autocorrelation in the data. The OLS model residuals were computed using the R package ‘ncf’ v.1.2.9 ^68^ and Moran’s I values were calculated using the R package ‘spdep’ v.1.1.5 ^69^.

To determine under which conditions evolutionary transitions to woodiness on islands may occur, we, in a second step, related the number of independent evolutionary transitions per archipelago with archipelago characteristics relevant under our research hypothesis using generalized linear regression models. We did this analysis on the archipelago rather than island level, since we could rarely trace back transitions to individual islands, and because archipelagos persist longer than individual volcanic islands, and hence are more relevant on evolutionary timescales. To cope with zero inflation due to a large number of archipelagos without any shifts, we used a hurdle model (i.e., a regression framework fitting separate models to the binary outcome—was there any transition present: yes or no?—and to the count reflecting the number of transitions) as implemented in the R package ‘pscl’ v.1.5.5 ^70^. We used a truncated Poisson generalized linear model with log-link modelling counts above zero (i.e., the number of IW transitions if any occurred) and a hurdle component modelling zero vs larger counts (i.e., on which archipelagos did IW transitions occur at all) using a binomial model and a logit link. See Supplementary Information Section 5 for details on model structure^70^. To test the *competition hypothesis*, we included the distance to the nearest continent as explanatory variable in the hurdle regression model. Based on the assumptions that colonizing herbaceous lineages in the absence of ancestrally woody trees compete for light (and hence are under selective pressure to evolve structurally supporting wood) and that ancestrally woody trees due to dispersal limitation are rarer on more isolated islands. *To test the favourable aseasonal climate hypothesis*, we included the mean absolute latitude of the archipelago centroid as explanatory variable in the hurdle regression model. Based on the assumption that on evolutionary time scales climate was more stable closer to the equator, we expect more evolutionary transitions to IW on islands at lower latitudes. To control for the effects expected under a neutral model of evolution, we included total archipelago land area and minimum archipelago age as explanatory variables in the hurdle regression model. Based on the assumption that a greater area harbours more evolutionary lineages and that a greater pool of evolutionary lineages leads to higher chance to observe transitions to IW, we expected more evolutionary transitions towards IW on larger archipelagos. Furthermore, based on the assumption that a longer evolutionary time increases the number of transitions, we expect more evolutionary transitions to IW on older archipelagos.

## Supporting information

SUpplementary Information Section 1

SUpplementary Information Section 4

SUpplementary Information Section 5

## Supporting Information

There is Supporting Information available in Sections 1-7.

## Data sharing and accessibility

The number and proportion of IWS per island and flowering plant family are available in the online Supplementary material of this manuscript. All analysis scripts necessary to replicate the analyses presented in this study will be available at a zenodo repository after publication.

## Acknowledgements

We dedicate this manuscript to the work of Sherwin Carlquist who dedicated his career to the study of island biology and comparative wood anatomy.

## References

1. Baeckens, S. & Van Damme, R. The island syndrome. Curr. Biol. 30, R338–R339 (2020).

2. Lomolino, M. V. The unifying, fundamental principles of biogeography: understanding Island Life. Front. Biogeogr. 8, (2016).

3. Losos, J. B. & Ricklefs, R. E. Adaptation and diversification on islands. Nature 457, 830–836 (2009).

4. Whittaker, R. J., Fernández-Palacios, J. M., Matthews, T. J., Borregaard, M. K. & Triantis, K. A. Island biogeography: Taking the long view of nature’s laboratories. Science 357, (2017).

5. Sayol, F., Steinbauer, M. J., Blackburn, T. M., Antonelli, A. & Faurby, S. Anthropogenic extinctions conceal widespread evolution of flightlessness in birds. Sci. Adv. 6, eabb6095 (2020).

6. Wright, N. A., Steadman, D. W. & Witt, C. C. Predictable evolution toward flightlessness in volant island birds. Proc. Natl. Acad. Sci. 113, 4765–4770 (2016).

7. Carthey, A. J. R. & Banks, P. B. Naïveté in novel ecological interactions: lessons from theory and experimental evidence. Biol. Rev. Camb. Philos. Soc. 89, 932–949 (2014).

8. Benítez-López, A. et al. The island rule explains consistent patterns of body size evolution in terrestrial vertebrates. Nat. Ecol. Evol. 5, 768–786 (2021).

9. Lomolino, M. V. et al. Of mice and mammoths: generality and antiquity of the island rule. J. Biogeogr. 40, 1427–1439 (2013).

10. Burns, K. C. Evolution in Isolation: The Search for an Island Syndrome in Plants. (Cambridge University Press, 2019). doi:10.1017/9781108379953.

11. Carlquist, S. J. Island biology. 1–686 (Columbia University Press, 1974). doi:10.5962/bhl.title.63768.

12. Lens, F., Davin, N., Smets, E. & del Arco, M. Insular woodiness on the Canary Islands: A remarkable case of convergent evolution. Int. J. Plant Sci. 174, 992–1013 (2013).

13. Doyle, J. A. Molecular and Fossil Evidence on the Origin of Angiosperms. Annu. Rev. Earth Planet. Sci. 40, 301–326 (2012).

14. Porter, M. L. & Crandall, K. A. Lost along the way: the significance of evolution in reverse. Trends Ecol. Evol. 18, 541–547 (2003).

15. Böhle, U. R., Hilger, H. H. & Martin, W. F. Island colonization and evolution of the insular woody habit in Echium L. (Boraginaceae). Proc. Natl. Acad. Sci. U. S. A. 93, 11740–11745 (1996).

16. Wallace, A. R. Island life; or, the phenomena and causes of insular faunas and floras, including a revision and attempted solution of the problem of geological climates. 1–598 (Macmillan, 1880). doi:10.5962/bhl.title.7552.

17. Darwin, C. On the Origin of Species: By Means of Natural Selection. (John Murray, 1859).

18. Givnish, T. J. Adaptive plant evolution on islands: classical patterns, molecular data, new insights. in Evolution on islands 281–304 (Oxford University Press, 1998).

19. Choat, B. et al. Triggers of tree mortality under drought. Nature 558, 531–539 (2018).

20. Dória, L. C. et al. Insular woody daisies (Argyranthemum, Asteraceae) are more resistant to drought-induced hydraulic failure than their herbaceous relatives. Funct. Ecol. 32, 1467–1478 (2018).

21. Lens, F. et al. Embolism resistance as a key mechanism to understand adaptive plant strategies. Curr. Opin. Plant Biol. 16, 287–292 (2013).

22. Kim, S. C. et al. Timing and tempo of early and successive adaptive radiations in Macaronesia. PLoS ONE 3, 1–7 (2008).

23. Nürk, N. M., Atchison, G. W. & Hughes, C. E. Island woodiness underpins accelerated disparification in plant radiations. New Phytol. 224, 518–531 (2019).

24. Hooft van Huysduynen, A. et al. Temporal and palaeoclimatic context of the evolution of insular woodiness in the Canary Islands. Ecol. Evol. 11, 12220–12231 (2021).

25. Patiño, J. et al. A roadmap for island biology: 50 fundamental questions after 50 years of The Theory of Island Biogeography. J. Biogeogr. 44, 963–983 (2017).

26. Hooker, J. Hooker, J. D. (1867). On insular floras: A lecture. Journal of Botany, 5, 23–31. J. Bot. 5, 23–31 (1867).

27. Johnson, M. A., Clark, J. R., Wagner, W. L. & McDade, L. A. A molecular phylogeny of the Pacific clade of Cyrtandra (Gesneriaceae) reveals a Fijian origin, recent diversification, and the importance of founder events. Mol. Phylogenet. Evol. 116, 30–48 (2017).

28. Price, J. P. & Wagner, W. L. Speciation in Hawaiian angiosperm lineages: cause, consequence, and mode. Evol. Int. J. Org. Evol. 58, 2185–2200 (2004).

29. Garrouste, R. et al. New fossil discoveries illustrate the diversity of past terrestrial ecosystems in New Caledonia. Sci. Rep. 11, 18388 (2021).

30. Knox, C., Eric B.,. Li. The East Asian origin of the giant lobelias. Am. J. Bot. 104, 924–938 (2017).

31. Losos, J. B. & Schluter, D. Analysis of an evolutionary species-area relationship. Nature 408, 847–850 (2000).

32. Clague, D. A. et al. The maximum age of Hawaiian terrestrial lineages: geological constraints from Kōko Seamount. J. Biogeogr. 37, 1022–1033 (2010).

33. Fernández-Palacios, J. M. et al. A reconstruction of Palaeo-Macaronesia, with particular reference to the long-term biogeography of the Atlantic island laurel forests. J. Biogeogr. 38, 226–246 (2011).

34. Wagner, W., Herbst, D. & Sohmer, S. Manual of the flowering plants of Hawaii. Revised edition. (University of Hawai‘i Press/Bishop Museum Press, 1999).

35. García-Verdugo, C., Caujapé-Castells, J. & Sanmartín, I. Colonization time on island settings: lessons from the Hawaiian and Canary Island floras. Bot. J. Linn. Soc. 191, 155–163 (2019).

36. Baldwin, B. G. A new look at phenotypic disparity and diversification rates in island plant radiations. New Phytol. 224, 8–10 (2019).

37. Spicer, R. & Groover, A. Evolution of development of vascular cambia and secondary growth. New Phytol. 186, 577–592 (2010).

38. Roodt, D., Li, Z., Van de Peer, Y. & Mizrachi, E. Loss of Wood Formation Genes in Monocot Genomes. Genome Biol. Evol. 11, 1986–1996 (2019).

39. Sauquet, H. & Magallón, S. Key questions and challenges in angiosperm macroevolution. New Phytol. 219, 1170–1187 (2018).

40. Marques, D. A., Meier, J. I. & Seehausen, O. A Combinatorial View on Speciation and Adaptive Radiation. Trends Ecol. Evol. 34, 531–544 (2019).

41. Melzer, T., S; Lens, F; Gennen, J; Vanneste, S; Rohde, A; Beeckman. Flowering-time genes modulate meristem determinacy and growth form in Arabidopsis thaliana. Nat. Genet. (2008).

42. Davin, N. et al. Functional network analysis of genes differentially expressed during xylogenesis in soc1ful woody Arabidopsis plants. Plant J. Cell Mol. Biol. 86, 376–390 (2016).

43. Larson, P. R. The Vascular Cambium - Development and Structure. (Springer Berlin Heidelberg, 1994).

44. Lens, F., Eeckhout, S., Zwartjes, R., Smets, E. & Janssens, S. B. The multiple fuzzy origins of woodiness within Balsaminaceae using an integrated approach. Where do we draw the line? Ann. Bot. 109, 783–799 (2012).

45. Lens, F., Smets, E. & Melzer, S. Stem anatomy supports Arabidopsis thaliana as a model for insular woodiness. New Phytol. 193, 12–17 (2012).

46. Kidner, C. et al. First steps in studying the origins of secondary woodiness in Begonia (Begoniaceae): combining anatomy, phylogenetics, and stem transcriptomics. Biol. J. Linn. Soc. 117, 121–138 (2016).

47. Freiberg, M. et al. LCVP, The Leipzig catalogue of vascular plants, a new taxonomic reference list for all known vascular plants. Sci. Data 416 (2020) doi:10.1038/s41597-020-00702-z.

48. Blomberg, S. P., Garland, T. & Ives, A. R. Testing for Phylogenetic Signal in Comparative Data: Behavioral Traits Are More Labile. Evolution 57, 717–745 (2003).

49. Pagel, M. Inferring the historical patterns of biological evolution. Nature 401, 877–884 (1999).

50. Revell, L. J. phytools: an R package for phylogenetic comparative biology (and other things). Methods Ecol. Evol. 3, 217–223 (2012).

51. Harris, L. W. & Davies, T. J. A Complete Fossil-Calibrated Phylogeny of Seed Plant Families as a Tool for Comparative Analyses: Testing the ‘Time for Speciation’ Hypothesis. PLOS ONE 11, e0162907 (2016).

52. Smith, S. A. & Brown, J. W. Constructing a broadly inclusive seed plant phylogeny. Am. J. Bot. 105, 302–314 (2018).

53. Kembel, S. W. et al. Picante: R tools for integrating phylogenies and ecology. Bioinformatics 26, 1463–1464 (2010).

54. Weigelt, P., König, C. & Kreft, H. GIFT – A Global Inventory of Floras and Traits for macroecology and biogeography. J. Biogeogr. 47, 16–43 (2020).

55. Global Aridity and PET Database. CGIAR-CSI https://cgiarcsi.community/data/global-aridity-and-pet-database/ (2018).

56. Zomer, R. J., Trabucco, A., Bossio, D. A. & Verchot, L. V. Climate change mitigation: A spatial analysis of global land suitability for clean development mechanism afforestation and reforestation. Agric. Ecosyst. Environ. 126, 67–80 (2008).

57. Fick, S. E. & Hijmans, R. J. WorldClim 2: new 1-km spatial resolution climate surfaces for global land areas. Int. J. Climatol. 37, 4302–4315 (2017).

58. Karger, D. N. et al. Climatologies at high resolution for the earth’s land surface areas. Sci. Data 4, 1–20 (2017).

59. Hijmans, R. J., Cameron, S. E., Parra, J. L., Jones, P. G. & Jarvis, A. Very high resolution interpolated climate surfaces for global land areas. Int. J. Climatol. 25, 1965–1978 (2005).

60. Weigelt, P., Steinbauer, M. J., Cabral, J. S. & Kreft, H. Late Quaternary climate change shapes island biodiversity. Nature 532, 99–102 (2016).

61. Faurby, S. et al. PHYLACINE 1.2: The Phylogenetic Atlas of Mammal Macroecology. Ecology 99, 2626–2626 (2018).

62. Onstein, R. E. et al. Palm fruit colours are linked to the broad-scale distribution and diversification of primate colour vision systems. Proc. R. Soc. B Biol. Sci. 287, 20192731 (2020).

63. Grace, J. B. et al. Guidelines for a graph-theoretic implementation of structural equation modeling. Ecosphere 3, 73 (2012).

64. Rosseel, Y. lavaan: An R Package for Structural Equation Modeling. J. Stat. Softw. 48, 1–36 (2012).

65. Stekhoven, D. J. missForest: Nonparametric Missing Value Imputation using Random Forest. (2013).

66. Stekhoven, D. J. & Buehlmann, P. MissForest - non-parametric missing value imputation for mixed-type data. Bioinformatics 28, 112–118 (2012).

67. Kissling, W. D. & Carl, G. Spatial autocorrelation and the selection of simultaneous autoregressive models. Glob. Ecol. Biogeogr. 17, 59–71 (2008).

68. Bjornstad, O. N. ncf: Spatial Covariance Functions. (2020).

69. Bivand, R. S. & Wong, D. W. S. Comparing implementations of global and local indicators of spatial association. TEST 27, 716–748 (2018).

70. Zeileis, A., Kleiber, C. & Jackman, S. Regression Models for Count Data in R. J. Stat. Softw. 27, 1–25 (2008).

