## Extended Data Figures for "Plant longevity, drought and island isolation favoured rampant evolutionary transitions towards insular woodiness"

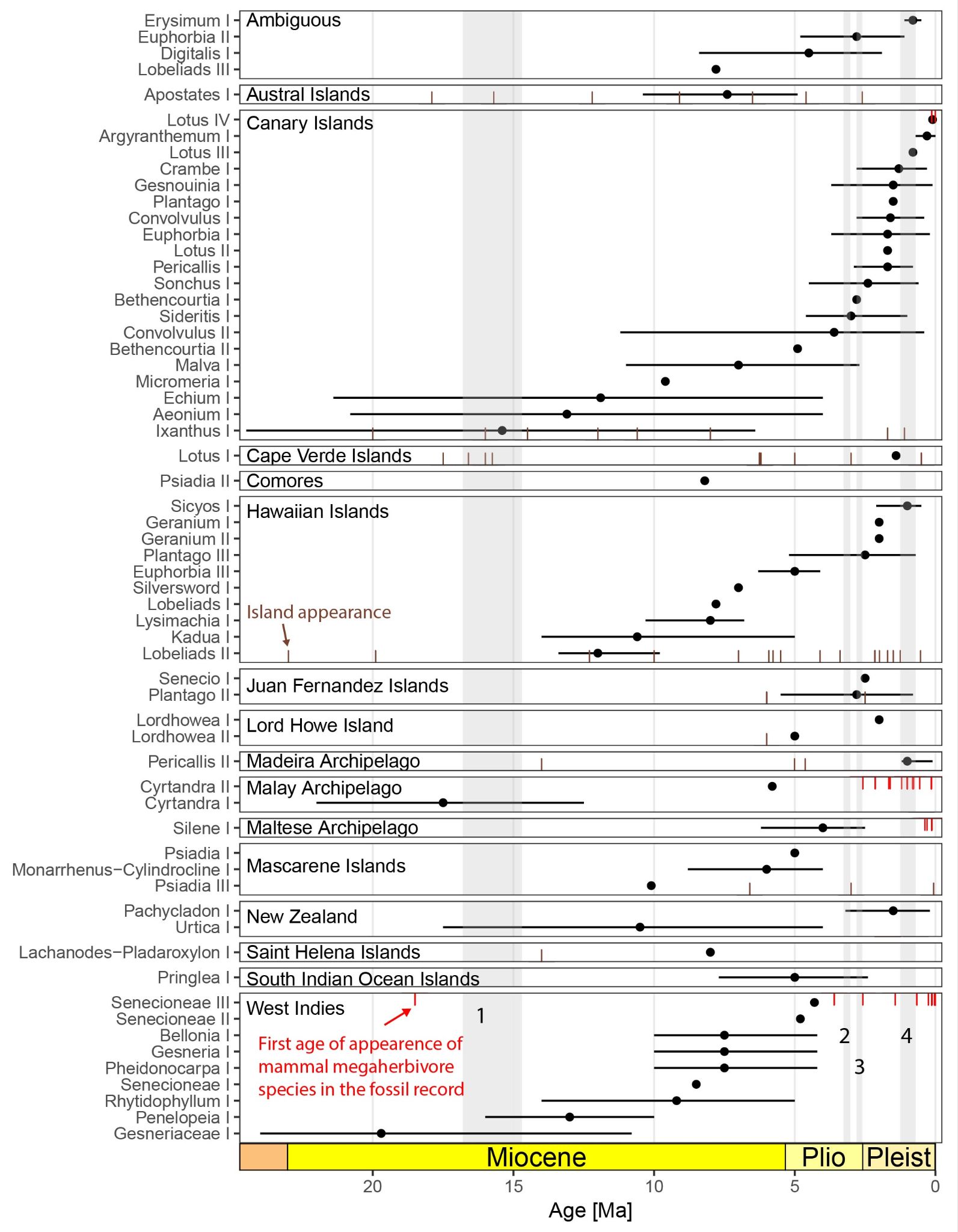

**Extended Data Figure 1.** The timing of evolutionary shifts to IW (stem age of insular woody clades) in relation to island age and number of herbivore species. The brown marks at the bottom of each panel show known island ages in the respective archipelago (only islands younger than 25 Ma shown); the red marks at the top show the first age of appearance of large mammal herbivores in the fossil record. Clades are labelled by their genus name for clarity, but may only include part of the genus. Important paleoclimatic events are marked in grey: 1: Miocene Climatic Optimum, 2: mid Pliocene warm period, 3: Onset of the Quaternary glacial cycles, 4: Middle Pleistocene transition. The island ages are only shown for exclusively volcanic archipelagos.

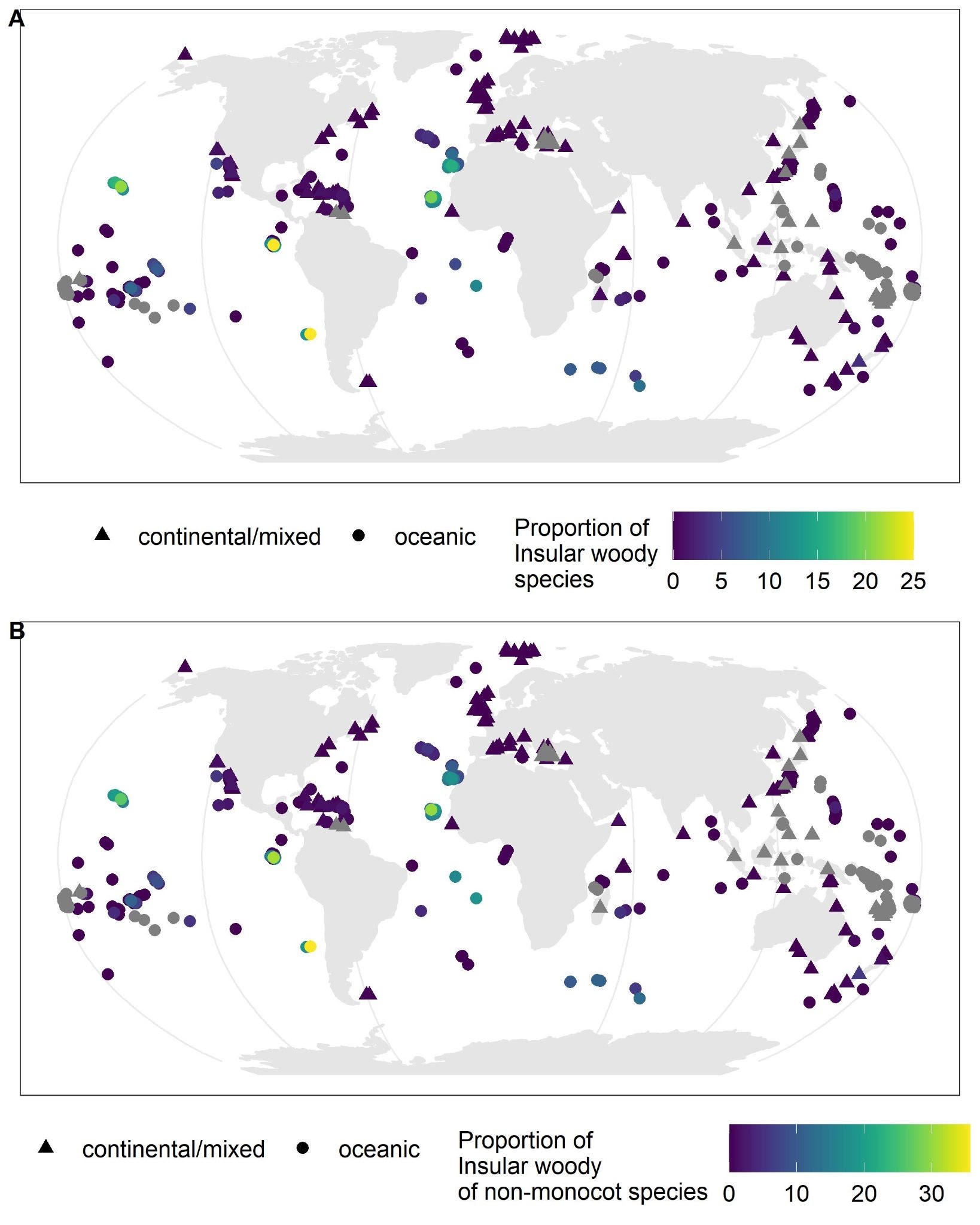
**Extended Data Figure 2.** The proportion of insular woody species on islands worldwide. **A)** The proportion of insular woody species on the total native angiosperm flora per island. **B)** The proportion of the native insular angiosperm flora excluding monocots (since monocots never produce wood and can therefore never evolve into insular woody lineages). Grey triangles indicate islands for which no estimates of total native angiosperm species richness were available.

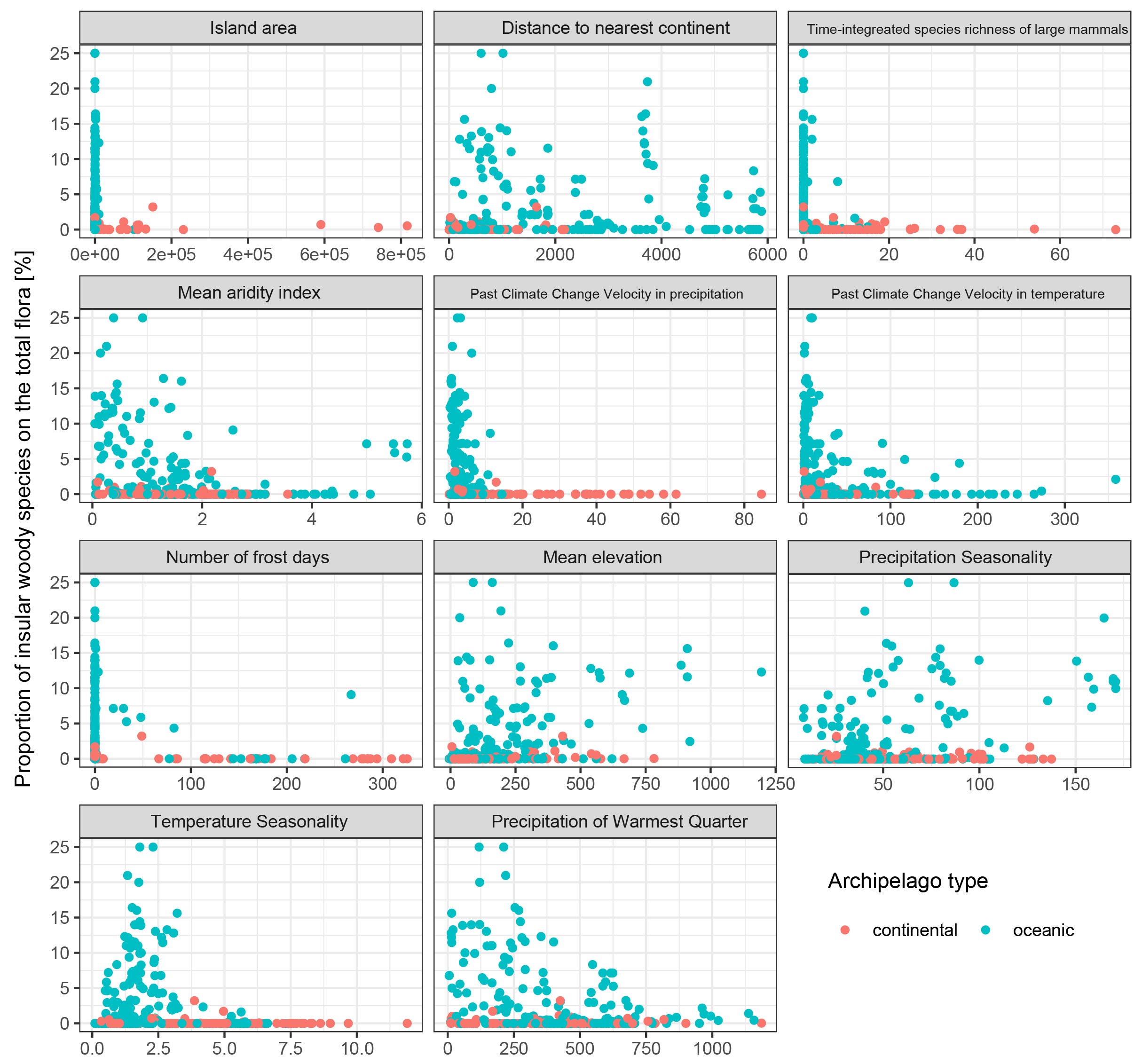

**Extended Data Figure 3.** The proportion of insular woody species on 322 islands worldwide plotted against the environmental variables included in the structural equation models that relate the number of insular woody species with island characteristics. See Figure S2 for model structure and Table S4 for effect size estimates in the combined model.

| **Hypothesis** | **Predictor** | **Expectation** | **Full model (n = 323)** | | **Full - Haw&Can (n = 308)** | | **Oceanic only (n = 203)** | | **Oceanic only  -Haw&Can (n = 188)** | |
| --- | --- | --- | --- | --- | --- | --- | --- | --- | --- | --- |
|  |  |  | **Estimate** | **Std.err** | **Estimate** | **Std.err** | **Estimate** | **Std.err** | **Estimate** | **Std.err** |
| **Number of insular woody species** | | | R2 = 0.552 | | R2 = 0.502 | | R2 =0.614 | | R2 = 0.499 | |
| neutral | Mean elevation | + | 0.399 | 0.05 | 0.249 | 0.045 | 0.324 | 0.064 | 0.214 | 0.055 |
| neutral | Total richness | + | 0.359 | 0.074 | 0.221 | 0.054 | 0.359 | 0.078 | 0.253 | 0.064 |
| herbivory | Mammal diversity | - | -0.319 | 0.054 | -0.286 | 0.045 | **ns** | **ns** | **-0.309** | **0.097** |
| drought | Aridity index | - | -0.196 | 0.068 | -0.178 | 0.044 | ns | ns | ns | ns |
| comeptition | Distance | + | 0.187 | 0.046 | 0.105 | 0.037 | 0.22 | 0.047 | 0.116 | 0.042 |
| drought | Precipitation seasonality | + | 0.163 | 0.057 | 0.145 | 0.046 | 0.184 | 0.055 | 0.193 | 0.044 |
| neutral | Area | + | 0.163 | 0.057 | 0.213 | 0.045 | ns | ns | ns | ns |
| drought | Precipitation warmest quarter | - | **-0.127** | **0.058** | **ns** | **ns** | -0.32 | 0.052 | -0.192 | 0.045 |
| climate | Temperature seasonality | - | -0.119 | 0.054 | -0.104 | 0.042 | ns | ns | ns | ns |
| climate | Number frost days | - | ns | ns | ns | ns | -0.19 | 0.052 | -0.122 | 0.043 |
| climate | Temperature change velocity | - | ns | ns | ns | ns | ns | ns | ns | ns |
| climate | Precipitation change velocity | - | ns | ns | ns | ns | -0.483 | 0.096 | -0.279 | 0.082 |
| **Total species richness** | | | R2 = 0.724 | | R2 = 0.733 | | R2 = 0.563 | | R2 = 0.569 | |
|  | Area | + | 0.498 | 0.026 | 0.481 | 0.026 | 0.524 | 0.047 | 0.478 | 0.051 |
|  | Distance | - | -0.281 | 0.026 | -0.304 | 0.027 | -0.264 | 0.032 | -0.289 | 0.034 |
|  | Precipitation warmest quarter | + | 0.263 | 0.037 | 0.288 | 0.038 | 0.287 | 0.053 | 0.34 | 0.055 |
|  | Number frost days | - | -0.259 | 0.032 | -0.245 | 0.033 | -0.194 | 0.055 | -0.15 | 0.057 |
|  | Precipitation seasonality | - | -0.19 | 0.036 | -0.204 | 0.037 | -0.198 | 0.048 | -0.213 | 0.049 |
|  | Temperature seasonality | - | 0.171 | 0.041 | 0.164 | 0.042 | 0.173 | 0.075 | 0.157 | 0.071 |
|  | Aridity index | + | -0.136 | 0.049 | -0.155 | 0.05 | -0.201 | 0.068 | -0.249 | 0.071 |
|  | Mean elevation | + | ns | ns | ns | ns | ns | ns | ns | ns |
| **Mammal diversity** | | | R2 = 0.462 | | R2 = 0.459 | | R2 = 0.410 | | R2 = 0.412 | |
|  | Temperature seasonality | - | 0.385 | 0.06 | 0.384 | 0.061 | 0.331 | 0.046 | 0.336 | 0.046 |
|  | Area | + | 0.305 | 0.043 | 0.298 | 0.044 | ns | ns | ns | ns |
|  | Distance | - | -0.222 | 0.034 | -0.225 | 0.036 | -0.1 | 0.023 | -0.094 | 0.024 |
|  | Number of frost days | - | -0.176 | 0.039 | -0.18 | 0.04 | ns | ns | ns | ns |
|  | Elevation | + | -0.14 | 0.048 | -0.116 | 0.053 | -0.057 | 0.027 | -0.067 | 0.031 |
|  | Aridity index | + | ns | ns | ns | ns | -0.1 | 0.023 | -0.09 | 0.034 |
|  | Precipitation seasonality | - | ns | ns | ns | ns | -0.082 | 0.034 | -0.076 | 0.035 |
|  | Precipitation warmest quarter | + | ns | ns | ns | ns | ns | ns | ns | ns |

**Extended Data Figure 4.** Effect sizes estimated in the structural equation models. Bold values differ between full models and sensitivity analyses without Hawaii and the Canaries.

| **Model** | **Component** | **Predictor** | **Coefficient estimate** | **Standard error** | **z** | **Pr(>\|z\|)** | **Significance level** |
| --- | --- | --- | --- | --- | --- | --- | --- |
| I | count | (Intercept) | 0.031 | 1.775 | 0.017 | 0.986 |  |
| I | count | Area | 0.319 | 0.11 | 2.913 | 0.004 | ** |
| I | count | Distance | -0.105 | 0.206 | -0.511 | 0.609 |  |
| I | count | Absolute latitude | -0.002 | 0.015 | -0.106 | 0.916 |  |
| I | count | Age | -0.240 | 0.264 | -0.91 | 0.363 |  |
| I | zero hurdle | (Intercept) | 1.5 | 2.106 | 0.712 | 0.476 |  |
| I | zero hurdle | Area | 0.040 | 0.137 | 0.295 | 0.768 |  |
| I | zero hurdle | Distance | -0.317 | 0.241 | -1.312 | 0.189 |  |
| I | zero hurdle | Absolute latitude | -0.008 | 0.017 | -0.455 | 0.649 |  |
| I | zero hurdle | Age | -0.149 | 0.325 | -0.46 | 0.646 |  |
| II | count | (Intercept) | -1.519 | 1.0 | -1.525 | 0.127 |  |
| II | count | Area | 0.573 | 0.080 | 7.167 | 7.68E-13 | *** |
| II | count | Distance | -0.261 | 0.083 | -3.137 | 0.002 | ** |
| II | count | Absolute latitude | 0.02 | 0.011 | 1.834 | 0.067 |  |
| II | count | Age | -0.118 | 0.222 | -0.529 | 0.597 |  |
| II | zero hurdle | (Intercept) | 1.086 | 2.037 | 0.533 | 0.594 |  |
| II | zero hurdle | Area | 0.106 | 0.130 | 0.818 | 0.414 |  |
| II | zero hurdle | Distance | -0.311 | 0.231 | -1.343 | 0.179 |  |
| II | zero hurdle | Absolute latitude | -0.0070 | 0.017 | -0.417 | 0.676 |  |
| II | zero hurdle | Age | -0.14 | 0.320 | -0.436 | 0.663 |  |
| III | count | (Intercept) | -0.893 | 1.477 | -0.605 | 0.545 |  |
| III | count | Area | 0.302 | 0.108 | 2.772 | 0.006 | ** |
| III | count | Distance | -0.024 | 0.173 | -0.137 | 0.891 |  |
| III | count | Absolute latitude | -0.003 | 0.014 | -0.249 | 0.803 |  |
| III | zero hurdle | (Intercept) | -0.304 | 1.590 | -0.191 | 0.848 |  |
| III | zero hurdle | Area | 0.059 | 0.111 | 0.53 | 0.596 |  |
| III | zero hurdle | Distance | -0.182 | 0.179 | -1.017 | 0.309 |  |
| III | zero hurdle | Absolute latitude | -0.003 | 0.016 | -0.161 | 0.872 |  |
| IV | count | (Intercept) | -1.7 | 0.925 | -1.839 | 0.066 |  |
| IV | count | Area | 0.555 | 0.071 | 7.774 | 7.63E-15 | *** |
| IV | count | Distance | -0.251 | 0.080 | -3.126 | 0.002 | ** |
| IV | count | Absolute latitude | 0.018 | 0.010 | 1.754 | 0.080 |  |
| IV | zero hurdle | (Intercept) | -0.66 | 1.546 | -0.427 | 0.670 |  |
| IV | zero hurdle | Area | 0.12 | 0.105 | 1.145 | 0.252 |  |
| IV | zero hurdle | Distance | -0.175 | 0.172 | -1.012 | 0.312 |  |
| IV | zero hurdle | Absolute latitude | -0.002 | 0.016 | -0.129 | 0.897 |  |

**Extended Data Figure 5.** Results of modelling the minimum number of evolutionary transitions from herbaceous to insular woodiness on archipelagos worldwide using a hurdle model.

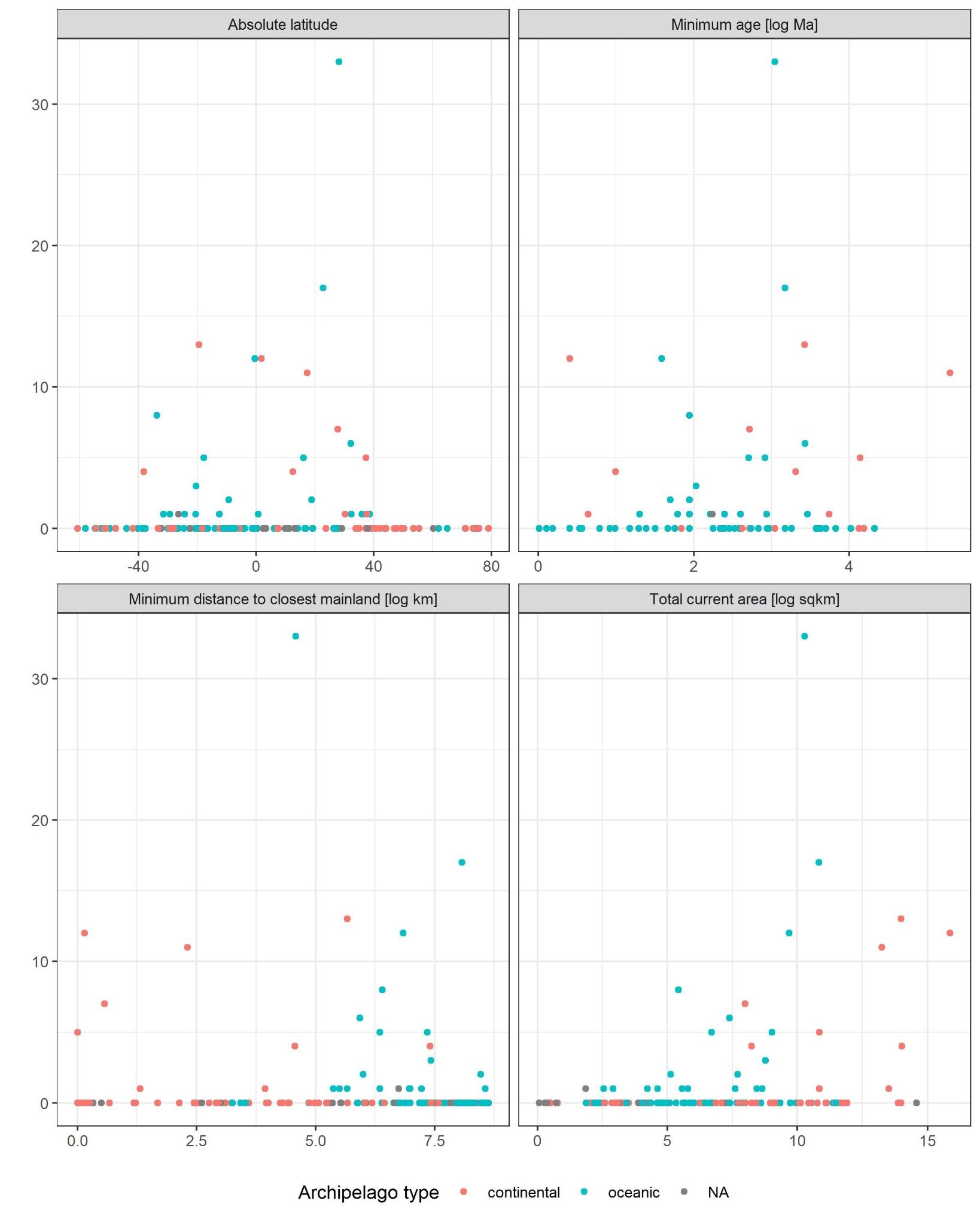

**Extended Data Figure 6.** Data used to model the minimum number of evolutionary transitions towards insular woodiness dependent on archipelago characteristics. For visualization the number of shifts is plotted against the predictor variables separately, each dot represents one archipelago (n = 45). See Supplementary Information Section 5 for information on model structure and Extended Data Figure 5 for effect size estimates.

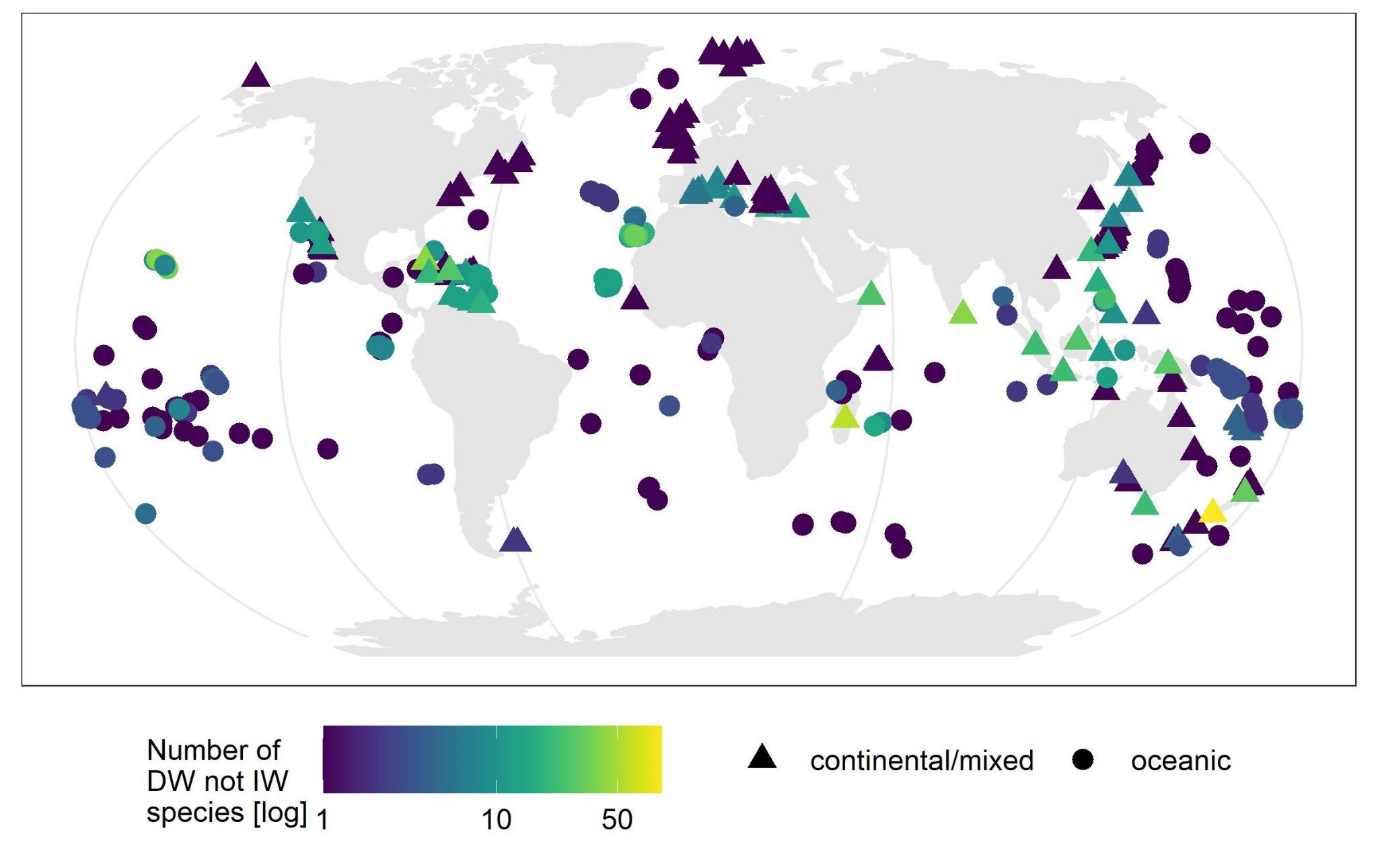

**Extended Data Figure 7.** The number of derived woody species that are not insular woody (i.e. dispersed to the islands after evolving woodiness on a nearby continent) on islands worldwide. Note that the number of derived wood species on New Zealand is much higher than the number of insular woody species due to the spectacular radiation of woody endemic veronicas that evolved their woodiness in Australia.
