## SUpplementary Information Section 1 for "Plant longevity, drought and island isolation favoured rampant evolutionary transitions towards insular woodiness"

Supplementary Figures and Tables

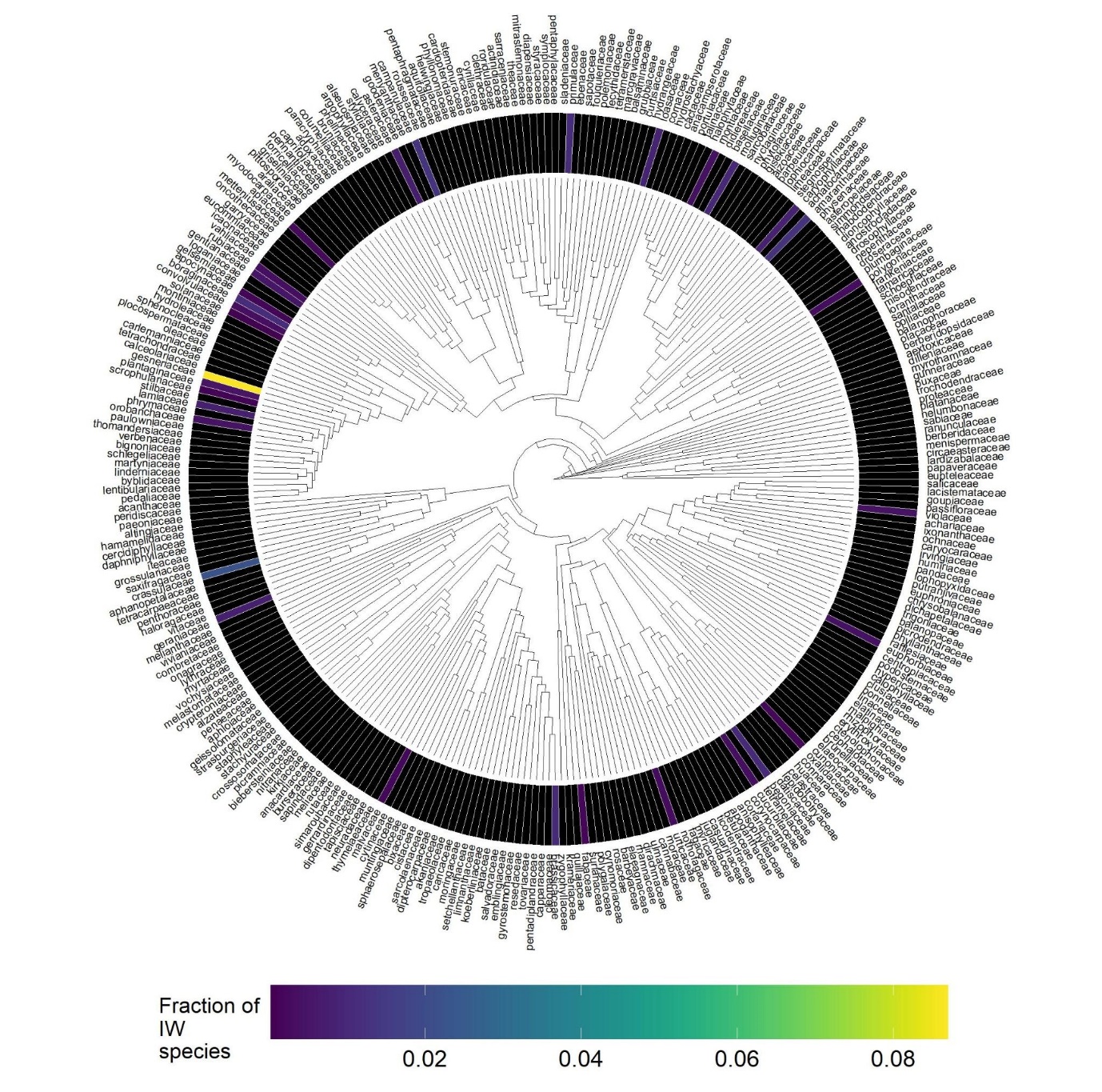

**Supplementary Figure 1.** The proportion of insular woody species per family across the eudicot Tree of Life. Black rectangles indicate families with no insular woody species.

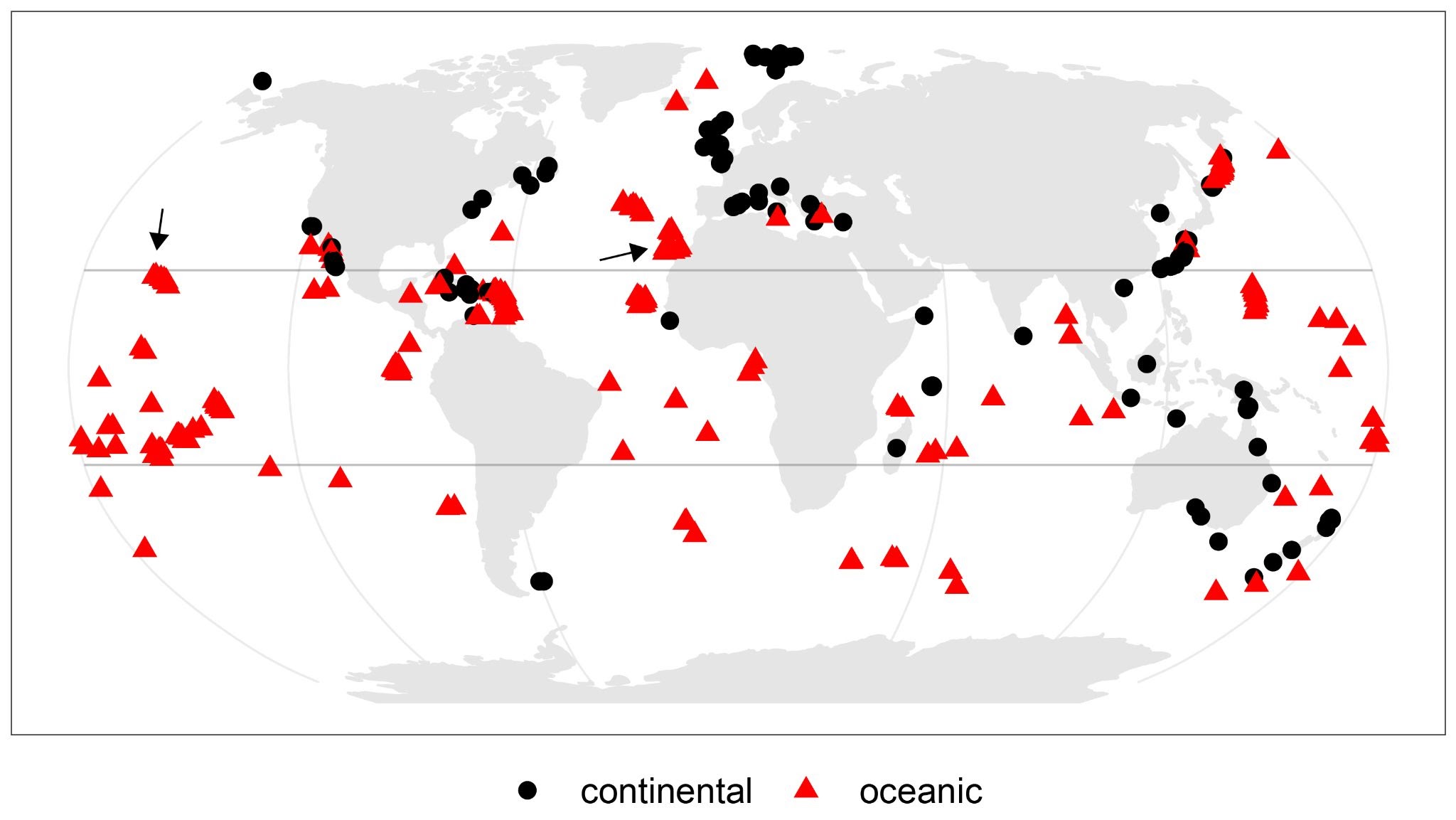

**Supplementary Figure 2.** The islands included in structural equation modelling (SEM) of the number of insular woody species and their geological origin. The arrows indicated the Hawaiian and Canary archipelagos, which we excluded in sensitivity analyses for the SEM (See Extended Data Figure 4 for results). The grey lines indicate the tropics of Cancer and Capricorn.

**Supplementary Table 1.** Environmental explanatory variables included in the structural equation models to predict the number of insular woody species on islands worldwide and the rationale for their inclusion in the analysis. The +/- signs in the parenthesis indicated if the expected effect is positive or negative.

| **Explanatory variable** | **Description** | **Explanation for inclusion (expected effect direction)** | **Related hypothesis** |
| --- | --- | --- | --- |
| IW species number | The number of insular woody species on the islands | This is the response variable. | - |
| Species number | The total number of angiosperm species on the islands as derived from GIFT flora checklists. | Under a neutral model, islands with more angiosperm species harbour more IWS (+). | control |
| Island area | The size of the island area | Under a neutral model, larger islands have more angiosperm species and hence more IWS (+). | control |
| Mean elevation | Mean elevation. | More IWS on islands with higher elevation are predicted due to higher habitat heterogeneity (+). | control |
| Distance to nearest continent | The distance to the nearest continental landmass estimating the extent of island isolation. | IW is the result of increased competition among herbaceous lineages for light in the absence of trees and shrubs (which are superior for competition for light). Assumption is that herbs are better colonizers, and hence reach isolated islands before trees and shrubs (+). | competition |
| Mean aridity index (AI) | Aridity is expressed as a generalized function of mean annual precipitation (MAP) and mean annual evapotranspiration (MAP), with lower AI values pointing to drier conditions. AI = MAP / MAE. | If IW is an adaptation to drought, more IWS are expected under drier conditions, i.e. lower AI values (-). | drought |
| Precipitation of Warmest Quarter | Precipitation of Warmest Quarter. | More IWS are predicted on islands with less precipitation in the warmest quarter (of the growing season) (-). | drought |
| Precipitation seasonality | Precipitation Seasonality, Coefficient of Variation. | More IWS are expected on islands with more pronounced differences in seasonal precipitation (+). | drought |
| Number of frost days | The number of days per year with a minimum temperature below 0°C | If IW is a result of a more stable favourable climate leading to continuous growth, a higher number of frost days is predicted negatively impact the occurrence of IWS (-). | favourable aseasonal climate |
| Climate change velocity in temperature | Climate Change Velocity in temperature since the last glacial maximum 21,000 y BP as the ratio between temporal change and contemporary spatial change in temperature. | If IW is a result of a favourable stable climate, more IWS are expected on islands with low temperature change velocity (-). | favourable aseasonal climate |
| Climate change velocity in precipitation | Climate Change Velocity in precipitation since the last glacial maximum 21,000 y BP as the ratio between temporal change and contemporary spatial change in precipitation. | If IW is the result of a favourable climate, less IWS are expected on islands with higher precipitation change velocity (-). | favourable aseasonal climate |
| Temperature Seasonality | Temperature Seasonality, standard deviation * 100. | More IWS are expected on islands with less temperature fluctuation between seasons (-). | favourable aseasonal climate |
| Time-integrated species richness of large mammal herbivores | The number of known fossil mammal species plus the number of potentially naturally distributed mammals. Only mammals with body mass > 1 kg and with > 20% plant material in the diet are taken into account. | If IW is the result of reduced herbivore pressure, more IWS are expected on islands with fewer native herbivore species (-). | reduced herbivory |

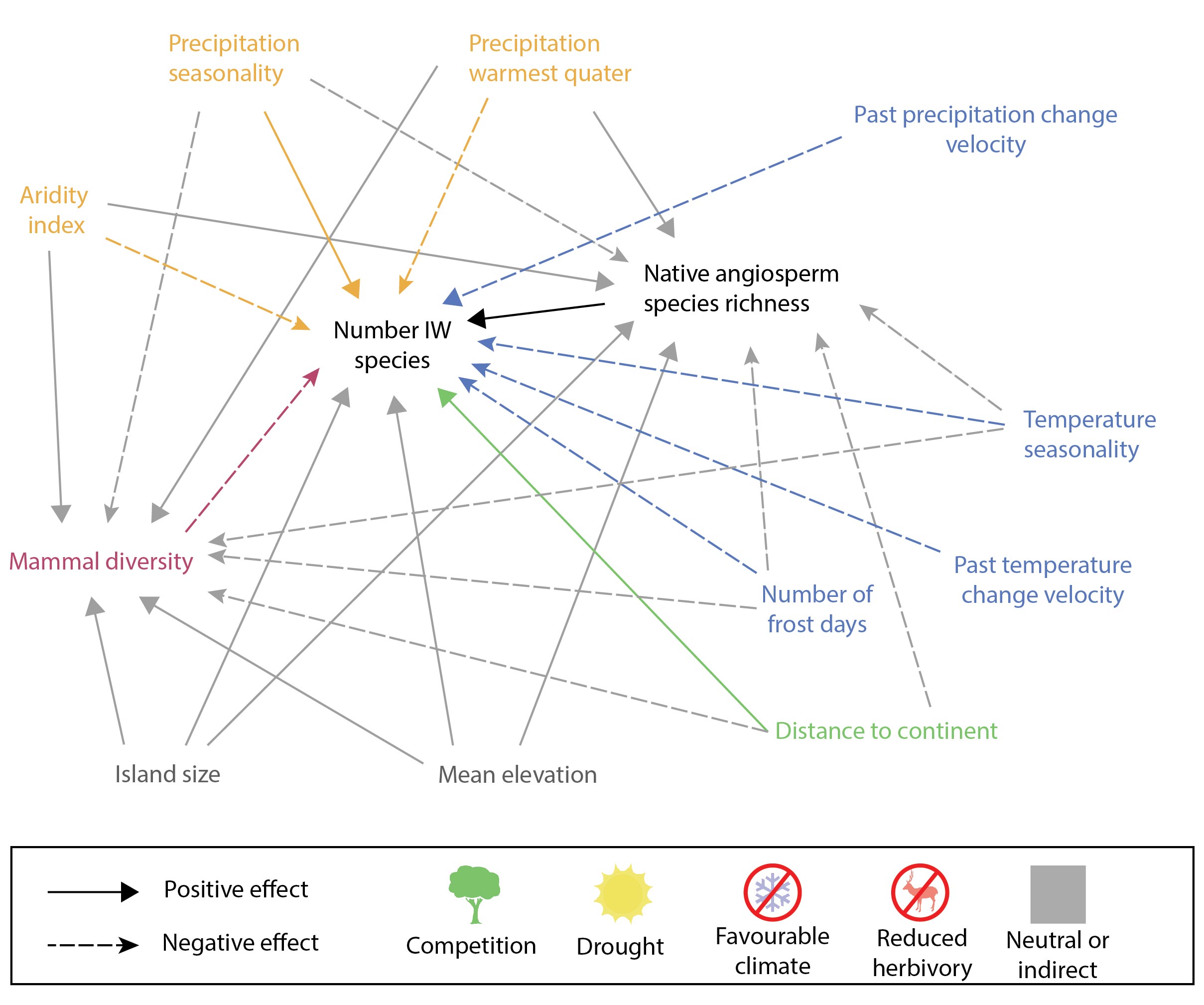

**Supplementary Figure 3** The structural equation models used to relate the number of insular woody species per island to island environment.

**Supplementary Table 2.** Predictors of the number of evolutionary transitions towards insular woodiness across archipelagos globally and the rationale for their inclusion in the analysis.

| **Explanatory variable** | **Description** | **Explanation for inclusion (expected effect direction)** | **Related hypothesis** |
| --- | --- | --- | --- |
| Archipelago area | The total land area of islands in this archipelago. | Under a neutral model, larger islands within a single archipelago harbour more lineages and thus a higher probability for lineages to shift to IW (+). | control |
| Minimum archipelago age | The oldest known age of an island in this archipelago. | Under a neutral model, lineages on older archipelagos had more evolutionary time to evolve into IW compared to younger archipelagos (+). | control |
| Distance to nearest continent | The minimum distance to the next continent's landmass. | Because more isolated archipelagos are less likely to be colonized by trees, herbaceous lineages that compete for light will lead to more IW shifts. (+). | competition |
| Absolute latitude | The absolute latitude of the archipelago centroid. | Islands closer to the equator experience more stable temperatures, also during evolutionary time. If IW is the result of favourable stable temperatures (especially lack of frost), more shifts towards IW are expected at lower absolute latitudes (-). | favourable aseasonal climate |
