## SUpplementary Information Section 4 for "Plant longevity, drought and island isolation favoured rampant evolutionary transitions towards insular woodiness"

Supplementary literature list

### Insular woody species identity

Adamo I. 2017. Investigating convergent evolutionary shifts towards derived woodiness within the Brassicaceae tribes Anastaticeae and Erysimeae: a molecular phylogenetic approach. MSc thesis, Leiden University, The Netherlands.

Adams, C. D. 1972. Flowering Plants of Jamaica. University of the West Indies, Mona, Jamaica.

Adamson RS 1955 The South African species of Aizoaceae. I *Adenogramma & Polpoda*. J S Afr Bot 21: 83-95.

Aedo C 2012 Revision of *Geranium* (Geraniaceae) in the New World. Systematic Botany Monographs 95: 1-550.

Aedo C, Aldaraso JJ, Saez L, Navarro C. 2003. Taxanomic revision of Geranium sect. Gracilia (Geraniaceae). Brittonia 55: 93-126.

Aellen P 1938 *Halimione* Aellen, eine rehabilitierte Chenopodiaceen-Gattung. *Verhandlungen der Naturforschenden Gesellschaft in Basel* 49: 118-130.

Aellen P 1940 *Exomis* und *Manochlamys* in Südafrika. *Bot Jahrb Syst* 70: 373-381.

Ahlenslager KE. 1984. Systematic studies of *Salvia* subgenus *Calosphace* section *Erythrostachys*. PhD thesis, University of Montana, USA.

Akhani H, G Edwards, EH Roalson 2007 Diversification of the Old World Salsoneae s.l. (Chenopodiaceae): molecular phylogenetic analysis of nuclear and chloroplast data sets and a revised classification. Int J Plant Sci 168: 931-956.

Albach DC, HM Meudt, B Oxelman 2005 Piecing together the “new” Plantaginaceae. Am J Bot 92: 297-315.

Allan GJ, J Fransisco-Ortega, A Santos-Guerra, E Boerner, EA Zimmer 2004 Molecular phylogenetic evidence for the geographic origin and classification of Canary Island *Lotus* (Fabaceae: Loteae). Mol Phyl Evol 32: 123-128.

Allan HH 1961 Flora of New Zealand, vol 1. pp1-1085, RE Owen, Government Printer, New Zealand.

Allan GJ, JM Porter 2000 Tribal delimitation and phylogenetic relationships of Loteae and Coronilleae (Faboideae: Fabaceae) with special reference to *Lotus*: evidence from nuclear ribosomal ITS sequences. Am J Bot 87: 1871-1881.

Al-Shehbaz IA 2010 A synopsis of the South American *Lepidium* (Brassicaceae). Darwiniana 48: 141-167.

Anderberg AA, BG Baldwin, RG Bayer, J Breitwieser, C Jeffrey, MO Dillon, P Eldenäs, V Funk, N Garcia-Jacas, DJN Hind, PO Karis, HW Lack, G Nesom, B Nordenstam, C. Oberprieler, J.L. Panero, C Puttock, H Robinson, FT Stuessy, A Sussana, E Urtubey, R Vogt, J Ward, LE Watson 2007 Compositae. Pages 61-588 *in* K Kubitzki, ed. The families and genera of vascular plants, vol VIII Asterales. Springer, Berlin.

Antonelli A 2009 Have giant lobelias evolved several times independently? Life form shifts and historical biogeography of the cosmopolitan and highly diverse subfamily Lobelioideae (Campanulaceae). BMC Biology 7: 82.

Applequist WL, WL Wagner, EA Zimmer, M Nepokroeff 2006 Molecular evidence resolving the systematic position of *Hectorella* (Portulacaceae). Syst Bot 31: 310-319.

Arias T, Pires JC. 2012. A fully resolved chloroplast phylogeny of the brassica crops and wild relatives (Brassicaceae: Brassiceae): novel clades and potential taxonomic implications. Taxon 61: 980-988.

Atkins H, Cronk QCB 2001. The genus *Cyrtandra* (Gesneriaceae) in Palawan, Philippines. Edinb. J. Bot. 58: 443-458.

Badré F, Cadet T., Cusset G., Hideux M. 1975. Position systématique, etude morphologique et palynologique du genre *Berenice*. Adansonia, ser. 2, 15: 139-146.

Badré F., Cadet T, Malplanche M 1972. Etude systématique et palynologique du genre Heterochaenia (Campanulaceae) endémique des Mascareignes. Adansonia ser 2, 12: 267-278.

Bakker M, Freek T, Culham A, Hettiarachi P, Touloumenidou T, Gibby M. 2004. Phylogeny of Pelargonium (Geraniaceae) based on DNA sequences from three genomes. TAXON, 53(1), 17–31. https://doi.org/10.2307/4135669

Baldwin BG. 1999. *Constancea*, a new genus for *Eriophyllum nevinii* (Compositae-Heliantheae s. lat.). Madrono 46: 159-160.

Baldwin BG. 2009. Chapter 41 Heliantheae alliance. Pages 689-711 *in* VA Funk, A Sussana, TF Stuessy, RJ Bayer, eds. Systematics, evolution, and biogeography of Compositae. International Association for Plant Taxonomy, Vienna.

Baldwin, BG and GD Carr. 2005. Dubautia kalalauensis, a new species of the Hawaiian silversword alliance (Compositae, Madiinae) from northwestern Kaua`i, U.S.A. Novon 15: 25-263.

Baldwin BG, DW Kyhos, J Dvorak, GD Carr. 1991. Chloroplast DNA evidence for a North American origin of the Hawaiian silversword alliance (Asteraceae). PNAS 88: 1840-1843.

Baldwin BG, Wessa BL, Panero JL. 2002. Nuclear DNA evidence for major lineages of helnioid Heliantheae (Compositae). Syst. Bot. 27: 161-198.

Baldwin K, BG Wood. 2016. Origin of the Rapa endemic genus Apostates: Revisiting major disjunctions and evolutionary conservatism in the Bahia alliance (Compositae: Bahieae). Taxon, 65(5), 1064–1080. https://doi.org/10.12705/655.8

Baldwin BG, MJ Sanderson 1998 Age and rate of diversification of the Hawaiian silversword alliance (Compositae). PNAS 95: 9402-9406.

Ballard HE Jr, KJ Sytsma 2000 Evolution and biogeography of the woody Hawaiian violets (*Viola*, Violaceae): arctic origins, herbaceous ancestry and bird dispersal. Evolution 54: 1521-1532.

Banasiak L, Wojewodzka A, Baczynski J, Reduron J-P, Piwczynski M, Kurzyna-Mlynik R, Gutaker R, Czarnocka-Cieciura A, Kosmala-Grzechnik S, Spalik K. 2016. Phylogeny of Apiaceae subtribe Daucinae and the taxonomic delineation of its genera. Taxon 65: 563-585.

Barber JC, J Francisco-Ortega, A Santos-Guerra, KG Turner, RK Jansen 2002 Origin of Macaronesian *Sideritis* L. (Lamioideae: Lamiaceae) inferred from nuclear and chloroplast sequence datasets. Mol Phyl Evol 23: 293-306.

Barres L, Sanmartin I, Anderson CL,. Susanna A, Buerki S. Galbany-Casals M, Vilatersana R. 2013. Reconstructing the evolution and biogeographic history of tribe Cardueae (Compositae). American Journal of Botany, 100(5), 867–882. https://doi.org/10.3732/ajb.1200058

Bartish IV, Aïnouche A, Jia D, Bergstrom D, Chown SL, Winkworth RC, Hennion F 2012 Phylogeny and colonization history of Pringlea antiscorbutica (Brassicaceae), an emblematic endemic from the South Indian Ocean Province. Molecular Phylogenetics and Evolution 65: 748–756

Bayer C, K Kubitzki 2003 Malvaceae. Pages 225-311 *in* K Kubitzki, ed. The families and genera of vascular plants, vol V Malvales, Capparales and non-betalain Caryophyllales. Springer, Berlin.

Beaman JH, Anderson C, Beaman RS. 2001. The plants of Mount Kinabalu 4: dicotyledon families Acanthaceae to Lythraceae. Kota Kinabalu: Natural History Publications.

Beentje H. 2002. Compositae, vol. 2. Flora of Tropical East Africa. Rotterdam: A.A. Balkema. 514-516.

Beilstein MA, IA Al-Shehbaz, S Mathews, EA Kellogg 2008 Brassicaceae phylogeny inferred from phytochrome A and ndhF sequence data: tribes and trichomes revisited. Am J Bot 95: 1307-1327.

Bernardello G, Anderson GJ, Stuessy TF, Crawford DJ. 2006. The angiosperm flora of the Archipelago Juan Fernandez: origin and dispersal. Can J Bot 84: 1266-1281.

Bittrich V 1993 Caryophyllaceae. Pages 206-236 *in* K Kubitzki, ed. The families and genera of vascular plants, vol II Magnoliid, Hamamelid and Caryophyllid families. Springer, Berlin.

Blake SF. 1913. A revision of *Encelia* and some related genera. Contributions from the Gray Herbarium of Harvard University 41: 346-396.

Bonati G 1926 Nouvelles Scrophulariacées Malgaches. Bulletin de la Sociéte Botanique de Genève 18: 1-40.

Borgen L. 1980. A new species of Argyranthemum (Compositae) from the Canary Islands. Norwegian Journal of Botany 27: 163-165.

Borgen L 1987 *Lobularia* (Cruciferae). A biosystematic study with special reference to the Macaronesian region. Op Bot 91: 1-96.

Boulos L. 1967. Nomenclatural changes and new taxa in Sonchus from the Canary Islands. Nytt Mag. Bot. 14: 7-18.

Boulos L, Friis I, Gilbert MG (1991) Notes on the Chenopodiaceae of Ethiopia, Somalia and Southern Arabia. Nordic Journal of Botany 11(3): 309-316

Brako L, Zarucchi JL. 1993. Catalogue of the flowering plants and gymnosperms of Peru. Monographs in Systematic Botany from the Missouri Botanical Garden 45: 1-1286.

Bramwell D. 1977. A revision of *Descurainia* Webb & Berth. section *Sisymbriodendron* (Christ) O.E.Schulz in the Canary Islands. Botanica Macaronesia 4: 31-53.

Bramwell D, ZI Bramwell 1974 Wild flowers of the Canary Islands. Stanley Thornes (Publishers) Ltd., London and Burford.

Brandbyge J 1993 Polygonaceae. Pages 531-544 *in* K Kubitzki, ed. The families and genera of vascular plants, vol II Magnoliid, Hamamelid and Caryophyllid families. Springer, Berlin.

Bräuchler C; Meimberg H, Heubl G. 2004. Molecular phylogeny of the genera Digitalis L. and Isoplexis (Lindley) Loudon (Veronicaceae) based on ITS-and trnL-F sequences. Plant Systematics and Evolution, 248, 111–128.

Bremer B, Eriksson T 2009 Time tree of Rubiaceae: phylogeny and dating the family, subfamilies, and tribes. Int J Plant Sci 170: 766-793.

Brochmann C, Rustan ØH, Lobin W Kilian N 1997 The endemic vascular plants of the Cape Verde Islands. Sommerfeltia 24: 1-356.

Brouillet L, TK Lowrey, L Urbatsch, V Karaman-Castro, G Sancho, S Wagstaff, JC Semple 2009 Chapter 37 Astereae. Pages 589-629 *in* VA Funk, A Sussana, TF Stuessy, RJ Bayer, eds. Systematics, evolution, and biogeography of Compositae. International Association for Plant Taxonomy, Vienna

Brown FBH 1935. Flora of southeastern Polynesia. III Dicotyledons. Bernice P. Bishop Museum Bulletin 130: 1-386.

Burke JM, A Sanchez, K Kron, M Luckow 2010 Placing the woody tropical genera of Polygonaceae: a hypothesis of character evolution and phylogeny. Am J Bot 97: 1377-1390.

Bush NA 1970 Flora of the U.S.S.R., vol VIII, part Cruciferae. Translated from Russian, Pp 13-453, Keter Press, Jerusalem.

Böhle UR, HH Hilger, WF Martin 1996 Island colonization and evolution of the insular woody habit in *Echium* L. (Boraginaceae). PNAS 93: 11740-11745.

Carine MA, SJ Russell, A Santos-Guerra, J Francisco-Ortega 2004 Relationships of the Macaronesian and Mediterranean floras: molecular evidence for multiple colonizations into Macaronesia and back-colonization of the continent in *Convolvulus* (Convolvulaceae). Am J Bot 91: 1070-1085.

Carlquist S. 1957. The genus *Fitchia* (Compositae). University of California Publications in Botany 29: 1-144.

Carvalho JA, Pontes T, Batista-Marques MI, Jardim R. 2010. A new species of *Echium* (Boraginaceae) from the island of Porto Santo (Madeira Archipelago). Anales del Jardín Botánico de Madrid: 67: 87-96.

Cavaco A. 1954. Flore de Madagascar et de Comores (ed. Humbert H), Family 67 Amaranthaceae, 2-56. Firmin-Didot and Company, Paris.

Caviño CI, SG Martínez, SR Downie 2010 Unraveling the taxonomic complexity of *Eryngium* L. (Apiaceae, Saniculoideae): phylogenetic analysis of 11 non-cding cpDNA loci corroborates rapid radiations. Plant Div Evol 128: 137-149.

Cheeseman, T.F., 1925. Urticaceae. In: Oliver, W.R.B. (Ed.), Manual of the New Zealand Flora, vol. 2. W.A.G. Skinner, Wellington, pp. 380–382.

Chen L-Y, Wang Q-F, Renner SS 2016 East Asian Lobelioideae and ancient divergence of a giant rosette Lobelia in Himalayan Bhutan. Taxon 65: 293-304.

Clark JR, WL Wagner, EH Roalson 2009 Patterns of diversification and ancestral range reconstruction in the southeast Asian-Pacific angiosperm lineage *Cyrtandra* (Gesneriaceae). Mol Phyl Evol 53: 982-994.

Crawford DJ. 1971. Morphology, chromosome number, and flavonoid chemistry of *Bidens cordylocarpa*. Madrono 21: 41-47.

Crawford DJ, Mesfin Tadesse, ME Mort, RT Kimball, CP Randle 2009 Chapter 42: Coreopsideae. Page 713-730 *in* VA Funk, A Susanna, T Stuessy, RJ Bayer, eds. Systematics, Evolution, and Biogeography of Compositae. International Association for Plant Taxonomy, Vienna.

Cronk QCB, M Kiehn, WL Wagner, JF Smith 2005 Evolution of *Cyrtandra* (Gesneriaceae) in the Pacific Ocean: the origin of a supertramp clade. Am J Bot 92: 1017-1024.

Cronk QCB 2000 The endemic flora of St Helena. Anthony Nelson Ltd, Oswestry.

Cupido CN, Prebble JM, Eddie WMM. 2013. Phylogeny of southern African and Australasian wahlenbergioids (Campanulaceae) based on ITS and trnL-F sequence data: implications for a reclassification. Systeatic Botany 38: 523-535.

Dalgaard V 1979 Biosystematics of the Macaronesia species of Scrophularia. Oper Botanica 51: 1-64.

Davies FG 1980 The genus *Gynura* (Compositae) in India, Sri Lanka and the Seychelles. Kew Bull 35: 365-367.

Davies FG 1981 The genus *Gynura* (Compositae) in Malesia and Australia. Kew Bull 35: 711-734.

Davis CJS 2010 Malva aethiopica, a new name for Lavatera abyssinica (Malvaceae): an endemic species of the Ethiopian Highlands. *Phytotaxa* 13: 15-18.

De Candolle A.P 1839. Order 55 Campanulaceae. In Prodromus Systematis Naturalis regne Vegetabilis, vol. 7, pp. 414-496. Treuttel and Würtz, Paris, France.

Descoings B 2003 *Kalanchoe*. Pages 143-181 *in* U Eggli, ed., Illustrated handbook of succulent plants: Crassulaceae. Springer, Berlin.

Dessein S 2003 Systematic studies in the Spermacoceae (Rubiaceae). PhD thesis, K.U.Leuven, Belgium.

Devesa JA 2000 Ononis. In: Talavera S. & al. (eds.), Flora Iberica, vol. 7(II). Real Jardín Botánico, Madrid: 590-646

Dillon MO, T Tu, A Soejima, T Yi, Z Nie, A Tye, J Wen 2007 Phylogeny of *Nolana* (Nolaneae, Solanoideae, Solanceae) as inferred from granule-bound starch synthase I (GBSSI) sequences. Taxon 56: 1000-1011.

Dostal J. 1976. *Centaura* L. *In* T. G. Tutin, V. H. Heywood, N. A. Burges, D. M. Moore, D. H. Valentine, S. M. Walters, and D. A. Webb [eds.], Flora Europaea, vol. 4, 254–301. Cambridge University Press, New York, New York, USA.

Dunbar-Co S, AM Wieczorek, CW Morden 2008 Molecular phylogeny and adaptive radiation of the endemic Hawaiian *Plantago* species (Plantaginaceae). Am J Bot 95: 1177-1188.

Eggens F, M Popp, M Nepokroeff, WL Wagner, B Oxelman 2007 The origin and number of introductions of the Hawaiian endemic *Silene* species (Caryophyllaceae) Am J Bot 94: 210-218.

Elliot GF. 1891. New and little-known Madagascar plants. The Journal of the Linnean Society 29: 1-66.

Epling C 1938 The Californian salvias. A review of *Salvia*, section *Audibertia*. Ann Missouri Bot Gard 25: 1-94.

Febles R. 2008. Re-estructutacion del genero Gonospermum Less. (Asteraceae: Anthemideae) en las Islas Canarias. Bot. Macaronesica 27: 101-105.

Ferguson IK. 1972. Verbascum. – In: Tutin, T. G. et al. (eds), Flora Europaea. Vol. 3. Cambridge Univ. Press, pp. 205–216.

Fernandez R 1968 Lavatera. In: Tutin TG, Heywood VH, Burges NA, Moore DM, Valentine DH, Walters SM, Webb DA (eds.), Flora Europaea, vol 2, pp 251-253. Cambridge Univesity Press, Cambridge.

Fischer E 2004 Scophulariaceae. Pages 333-432 *in* K Kubitzki, ed. The families and genera of vascular plants, vol VII Lamiales (except Acanthaceae including Avicenniaceae). Springer, Berlin.

Fishbein M, Livshultz T, Straub SCK, Simoes AO, Boutte J, McDonnell, A, Foote A. 2018. Evolution on the backbone; Apocynaceae phylogenomics and new perspectives on growth forms, flowers, and fruits. American Journal of Botany 105: 495-513.

Fiz O, Vargas P, M Alarcón, C Aedo, LJ García, JJ Aldasoro 2008 Phylogeny and historical biogeography of Geraniaceae in relation to climate changes and pollination ecology. Syst Bot 33: 326-342.

Florence J, Lorence DH 2000 Sertum polynesicum VI. Rubiaceae nouvelles des îles Marquises (Polynésie française). 2. Le genre Hedyotis. Adansonia, sér 3, 22: 223-230.

Florence J 1985 Sertum polynesicum I. Plakothira Florence (Loacaceae), genre nouveau des îles Marquises. Bull. Mus. Natn. Hist. Nat. Paris 4^th^ series, sect B, Adansonia nr 3: 329-245.

Fosberg, F.R. & Sachet, M.H. 1991. Studies in Indo-Pacific Rubiaceae. *Allertonia* 6: 191–278.

Francisco-Ortega, R., J; Santos-Guerra, A; Hines, A; Jansen. (1997). Molecular evidence for a Mediterranean origin of the Macaronesian endemic genus Argyranthemum (Asteraceae). Am J Bot, 84.

Franzke A, D German, IA Al-Shebahz, K Mummenhoff 2009 *Arabidopsis* family ties: molecular phylogeny and age estimates in Brassicaceae. Taxon 58: 425-437.

Fryxell PA 1988 Malvaceae of Mexico. Systematic Botany Monographs 25: 1-522.

Fuentes-Bazan, T., S; Mansion, G; Borsch. (2012). Towards a species level tree of the globally diverse genus Chenopodium (Chenopodiaceae). Mol Phylogenetics Evol, 62.

Fuertes-Aguilar J, MF Ray, J Francisco-Ortega, A Santos-Guerra, RK Jansen 2002 Molecular evidence from chloroplast and nuclear markers for multiple colonizations of *Lavatera* (Malvaceae) in the Canary Islands. Syst Bot 27: 74-83.

Ganders FR, M Berbee, M Pirseyedi. 2000. ITS base sequence phylogeny in *Bidens* (Asteraceae): Evidence for the continental relatives of Hawaiian and Marquesan Bidens. Syst Bot 25: 122-133.

Garcia-Jacas N, T Uysal, K Romashchenko, VN Suárez-Santiago, K Ertuğrul, A Susanna 2006 *Centaurea* revisited: a molecular survey of the *Jacea* group. Ann Bot 98: 741-753.

García-Maroto F, A Mañas-Fernández, JA Garrido-Cárdenas, D López Alonso, JL Guil-Guerrero, B Guzmán, P Vargas 2009 Δ^6^-Desaturase sequence evidence for explosive Pliocene radiations within the adaptive radiation of Macaronesian *Echium* (Boraginaceae). Mol Phyl Evol 52: 563-574.

Germishuizen G, Meyer NL. 2003. Plants of southern Africa: an annotated checklist. Strelitzia 14: 1-1231.

Ghahremaninejad F, Riahi M, Babaei M, Attar F, Behcet L, Sonboli A. 2014. Monophyly of Verbascum (Scrophularieae : Scrophulariaceae): evidence from nuclear and plastid phylogenetic analyses. Australian Journal of Botany 62: 638–646.

Givnish TK, KC Millam, AR Mast, TB Paterson, TJ Theim, AL Hipp, JM Henss, JF Smith, KR Wood, KJ Systsma 2009 Origin, adaptive radiation and diversification of the Hawaiian lobeliads (Asterales: Campanulaceae). Proc R Soc B 276: 407-416.

Goodall-Copestake WP, S Pérez-Espona, DJ Harris, PM Hollingsworth 2010 The early evolution of the mega-diverse genus *Begonia* (Begoniaceae) inferred from organelle DNA phylogenies. Biol J Linnean Soc 101: 243–250.

Goodson BA, A Santos-Guerra, RK Jansen 2006 Molecular systematics of *Descurainia* (Brassicaceae) in the Canary Islands: biogeographic and taxonomic implications. Taxon 55: 671-682.

Grau, J. 1981: Scrophularia in Rechinger, K. H. (ed.), Flora Iranica 147: 213-284. -Akademische Druck. Verlagsanstalt Graz, Graz.

Greenberg AK, MJ Donoghue 2011 Molecular systematics and character evolution in Caryophyllaceae. Taxon 60: 1637-1652.

Grisebach AHR 1864 Flora of the British West Indian Islands pp 459-466. London, Lovell Reeve & Co

Groeninckx I, Dessein S, Ochoterena H, Persson C, Motley TJ, Karehed J, Bremer B, Huysmans S, Smets E 2009 Phylogeny of the herbaceous tribe Spermacoceae (Rubiaceae) based on plastid DNA data. Ann. Missouri Bot. Garden 96: 109–132.

Groeninckx I, de Block P, Rakotonasolo F, Smets, E, Dessein S 2009 Rediscovery of Malagasy *Lathraeocarpa* allows determination of its taxonomic position within Rubiaceae. Taxon 58: 209–226.

Groeninckx I, P De Block, E. Robbrecht, EF Smets, S Dessein 2010 *Amphistemon* and *Thamnoldenlandia*, two new genera of Rubiaceae (Spermacoceae) endemic to Madagascar. Bot J Linn Soc 163: 447-472.

Groeninckx I, de Block P, Rakotonasolo F, Smets, E, Dessein S 2009 Rediscovery of Malagasy *Lathraeocarpa* allows determination of its taxonomic position within Rubiaceae. Taxon 58: 209–226.

Grosse-Veldmann B, Nürk NM, Smissen R, Breitwieser I, Quandt D, Weigend M. 2016. Pulling the sting out of nettle systematics – A comprehensive phylogeny of the genus Urtica L. (Urticaceae). Molecular Phylogenetics and Evolution 102: 9–19.

Gruenstaeudl M, Santos-Guerra, A, Jansen RK. 2013. Phylogenetic analyses of Tolpis Adans. (Asteraceae) reveal patterns of adaptive radiation, multiple colonization and interspecific hybridization. Cladistics 29: 416-434.

Guo X, Wang RJ, Simmons MP, But PPH. & Yu, J. 2013. Phylogeny of the Asian *Hedyotis-Oldenlandia* complex (Spermacoceae, Rubiaceae): Evidence for high levels of polyphyly and the parallel evolution of diplophragmous capsules. *Molec. Phylogen. Evol.* 67: 110–122.

Hao G, Y-M Yuan, C-M Hu, X-J Ge, N-X Zhao 2004 Molecular phylogeny of *Lysimachia* (Myrsinaceae) based on chloroplast *trn*L-F and nuclear ribosomal ITS sequences. Mol Phyl Evol 31: 323-339.

Harbaugh DT, M Nepokroeff M, RK Rabeler, J McNeill, EA Zimmer, WL Wagner. 2010. A new lineage-based tribal classification of the family Caryophyllaceae. Int J Plant Sci 171: 185-198.

Harwood R, Dessein S. 2005. Australian Spermacoce (Rubiaceae: Spermacoceae). I. Northern Territory. Australian Systematic Botany, 18(4), 297. https://doi.org/10.1071/sb03024

Havran C, KJ Sytsma, HE Ballard Jr 2009 Evolutionary relationships, interisland biogeography, and molecular evolution in the Hawaiian violets (*Viola*: Violaceae). Am J Bot 96: 2087-2099.

Hedge IC, Miller AG 1977. New and interesting taxa from NE tropical Africa. Notes RBG Edinb. 35: 179-193.

Hedge C 1974. A revision of *Salvia* in Africa including Madagascar and the Canary Islands. Notes RBG Edinb 33: 1-121.

Heenan PB, Goeke DF, Houliston GJ, Lysak MA. 2012. Phylogenetic analyses of ITS and *rbcL* DNA sequences for sixteen genera of Australian and New Zealand Brassicaceae result in the expansion of the tribe Microlepidieae. Taxon 61: 970-979.

Heenan PB, Mitchell AD, Koch M. 2002. Molecular systematics of the New Zealand *Pachycladon* (Brassicaceae) complex: generic circumscription and relationships to *Arabidopsis* sens. lat. and *Arabis* sens. lat. New Zealand Journal of Botany 40: 543-562.

Heenan PB. 1995. A taxonomic revision of Carmichaelia (Fabaceae — Galegeae) in New Zealand (part I). New Zealand Journal of Botany 33: 455-475.

Heenan PB. 1996. A taxonomic revision of Carmichaelia (Fabaceae — Galegeae) in New Zealand (part II). New Zealand Journal of Botany 34: 157-177.

Heenan PB. 1998. An emended circumscription of *Carmichaelia*, with new combinations, a key, and notes on hybrids, New Zealand Journal of Botany 36: 53-63.

Heenan PB. 2000. *Clianthus* (Fabaceae) in New Zealand: a reappraisal of Colenso’s taxonomy. New Zealand Journal of Botany 38 361-371.

Heenan PB. 2001. Relationships of *Streblorrhiza* (Fabaceae), an extinct monotypic genus from Phillip Island, South Pacific Ocean, New Zealand Journal of Botany 39: 9-15.

Hellwig FH 2004 Centaureinae (Asteraceae) in the Mediterranean – history of ecogeographical radiation. Plant Syst Evol 246: 137-162.

Hewson HJ 1981 The genus *Lepidium* L. (Brassicaceae) in Australia. *Brunonia* 4: 217-308.

Heywood VH and Akeroyd JR. 1993. Brassica in Flora Europea, vol 1, 2^nd^ edition, Tutin TG, Burges NA, Chater AO, Edmondson JR, Heywood VH, Moore DM, Valentine DH, Walters SM, Webb DA. (eds.), pp. 405-409, Cambridge University Press, Cambridge UK.

Hilliard OM, BL Burtt 1971 *Streptocarpus*. An African plant study. Pietermaritzburg, University of Natal Press.

Hilliard OM, BL Burtt 2002 The genus *Agalmyla* (Gesneriaceae-Cyrtandroideae). Edinb J Bot 59: 1-210.

Horn JW, BW van Ee, JJ Morawetz, R Riina, VW Steinmann, PE Berry, KJ Wurdack 2012 Phylogenetics and the evolution of major structural changes in the giant genus *Euphorbia* L. (Euphorbiaceae). Mol Phyl Evol 63: 305-326.

Howell JT. 1942. A list of vascular plants from Guadeloupe Island, lower California. Leaflets of Western Botany 3: 145-155.

Huang X; Denga T, Moore MJ, Wang H, Li Z, Lin N, Yusupov Z, Tojibaev KS, Wang Y, Sun H. 2019. Tropical Asian Origin, boreotropical migration and long-distance dispersal in Nettles (Urticeae, Urticaceae). Molecular Phylogenetics and Evolution, 137, 190–199.

Hufford L, MM McMahon, AM Sherwood, G Reeves, MW Chase. 2003. The major clades of Loasaceae: phylogenetic analysis using the plastid *matK* and *trnL-trnF* regions. Am J Bot 90: 1215-1228.

Humbert H 1960 Flore de Madagascar et des Comores. 189^e^ Famille Composées. Tome 1. Firmin-Didot et C^ie^, Paris.

Humbert H 1971. Pages 47-160 *in* Humbert H, ed. Flore de Madagascar et des Comores. Natural History Museum, Paris.

Humphries CJ 1976 A revision of the Macaronesian genus Argyranthemum Webb ex Schultz Bip. (Compositae-Anthemideae). Bulletin of the British Museum (Natural History) Botany 5: 147-240

Jeffrey C. 1988. The tribe Senecioneae (Compositae) in the Mascarene Islands with an annotated world checklist of the genera of the tribe. Notes on Compositae: VI. Kew Bulletin 43: 49-109.

Jeppesen S 1981 Campanulaceae. Pages 1-184 *in* Harling G, B Sparre, eds. Flora of Ecuador, nr 14. University of Göteborg, Göteborg.

Johnson MA, Pillon Y, Sakishima T, Price DK, Stacy EA. 2019. Multiple colonizations, hybridization and uneven diversification in Cyrtandra (Gesneriaceae) lineages on Hawai’i Island. Journal of Biogeography, 46, 1178–1196.

Johnson MA, Clark JR, Wagner WL, McDade LA. 2017. A molecular phylogeny of the Pacific clade of *Cyrtandra* (Gesneriaceae) reveals a Fijian origin, recent diversification, and the importance of founder effects. Molecular Phylogenetics and Evolution 116: 30-48.

Johnston IM. 1931. The flora of the Revillagigedo Islands. Proceedings of the California Academy of Sciences 20: 9-104.

Jones KE, Reyes-Betancort JA, Hiscock SJ, Carine MA. 2014. Allopatric diversification, multiple habitat shifts, and hybridization in the evolution of Pericallis (Asteraceae), a Macaronesian endemic genus. American Journal of Botany 101: 637-651.

Jørgensen, P. M., M. H. Nee & S. G. Beck. (eds.) 2014. Cat. Pl. Vasc. Bolivia, Monogr. Syst. Bot. Missouri Bot. Gard. 127(1–2): i–viii, 1–1744. Missouri Botanical Garden Press, St. Louis.

Kadereit G, EV Mavrodiev, EH Zacharias, AP Sukhorukov 2010 Molecular phylogeny of Atripliceae (Chenopodieae, Chenopodiaceae): implications for systematics, biogeography, flower and fruit evolution, and the origin of C4 photosynthesis. Am J Bot 97: 1664-1687.

Kadereit G, H Freitag 2011 Molecular phylogeny of Camphorosmeae (Camphorosmoideae, Chenopodiaceae): implications for biogeography, evolution of C4-photosynthesis and taxonomy. Taxon 60: 51-78.

Kato M, Nagamasu H. 1995. Dioecy in the Endemic Genus *Dendrocacalia* (Compositae) on the Bonin (Ogasawara) Islands. *J. Plant Res.* 108: 443-450.

Keeley SC, H Robinson 2009 Chapter 28: Vernonieae. Pages 439-469 *in* VA Funk, A Sussana, TF Stuessy, RJ Bayer, eds. Systematics, evolution, and biogeography of Compositae. International Association for Plant Taxonomy, Vienna.

Kidner C, Groover A, Thomas D, Emelianova K, Soliz-Gamboa C, Lens F. 2016. First steps in studying the origins of secondary woodiness in *Begonia* (Begoniaceae): combining anatomy, phylogenetics, and stem transcriptomics. *Biological Journal of the Linnean Society* 117: 121-138

Kilian N, B Gemeinholzer, HW Lack. 2009. Chapter 24: Cichorieae. Pages 343-383 *in* VA Funk, A Sussana, TF Stuessy, RJ Bayer, eds. Systematics, evolution, and biogeography of Compositae. International Association for Plant Taxonomy, Vienna.

Kilian N, Galbany-Casals M, Sommerer R, Oberprieler C, Smissen R, Miller A, Rabe K. 2017. Systematics of *Libinhania*, a new endemic genus of Gnaphalieae (Asteraceae) from the Socotra archipelago (Yemen), inferred from plastid, low-copy nuclear and nuclear ribosomal DNA loci. 2017. Botanical Journal of the Linnean Society 183: 373-412.

Kim SC, MR McGowen, P Lubinsky, JC Barber, ME Mort, A Santos-Guerra 2008 Timing and tempo of early and successive adaptive radiations in Macaronesia. PLos ONE 3: e2139.

Kim S-C, L Chunghee, JA Mejias 2007 Phylogenetic analysis of chloroplast DNA matK gene and ITS of nrDNA sequences reveals polyphyly of the genus *Sonchus* and new relationships among the subtribe Sonchinae (Asteraceae: Cichorieae). Mol Phyl Evol 44: 578-597.

Kissling J, Y-M Yuan, P Küpfer, G Mansion 2009 The polyphyletic genus *Sebaea* (Gentianaceae): a step forward in understanding the morphological and karyological evolution of the Exaceae. Mol Phyl Evol 53: 734-748.

Klackenberg J 1985 The genus *Exacum* (Gentianaceae). Opera Bot 84: 1-144.

Klackenberg J 1987 Revision of the genus *Tachiadenus*. Bull Mus Natn Hist Nat, Paris, 4^th^ ser, 9, section B, Adansonia 1: 43-80.

Klackenberg J 1990 Famille 168. Gentianacées. Famillie 168nis. Menyanthacées. Pages 1-185 in Morat P (ed) Flore de Madagascar et de Comores. Association de Botanique Tropical, Paris, France.

Klackenberg J 2002 Tribe Exaceae. Pages 66-108 *in* L Struwe, V Albert, eds. Gentianaceae: systematics and natural history. Cambridge University Press, Cambridge.

Knope ML; Morden CW, Funk VA, Fukami T 2012. Area and the rapid radiation of Hawaiian Bidens (Asteraceae). Journal of Biogeography, 39, 1206–1216.

Knox EB, Li C 2017. The East Asian origin of the giant lobelias. American Journal of Botany, 104, 924–938. https://doi.org/10.3732/ajb.1700025

Knox EB, Luke, Q, Thulin M 2004. A new giant Lobelia from the Eastern Arc Mts, Tanzania. Kew Bulletin, 59, 189–194.

Knox T, EB, Pocs T. 1992. Lobelia morogoroensis: Another Tanzanian giant. Kew Bulletin, 47, 503–508.

Kondraskov P, Schutz N, Schussler C, Menezes de Sequeira M, Guerra AS, Caujape-Castells J, Jaen-Molina R, Marrero-Rodríguez A, Koch MA, Linder P, Kovar-Eder J, Thiv M 2015. Biogeography of Mediterranean hotspot biodiversity: Re-evaluating the ‘Tertiary Relict’ hypothesis of Macaronesian laurel forests. PLoS One, 10, e0132091.

Kool A 2012 Desert plants and deserted islands: systematics and ethnobotany in Caryophyllaceae. Digital Comprehensive Summaries of Uppsala Dissertations from the Faculty of Science and Technology 972.

Kool A, Thulin M. 2017. A giant spurrey on a tiny island: on the phylogenetic position of Sanctambrosia manicata (Caryophyllaceae) and the generic circumsciptions of *Spergula* and *Rhodalsine*. Taxon 66: 615-622.

Koster JT 1970 The Compositae of New Guinea II. Blumea 18: 137-145.

Kramina TE, Degtjareva GV, Samigullin TH, Valiejo-Roman CM, Kirkbride JH, Volis S, Deng T, Sokoloff DD 2016. Phylogeny of Lotus (Leguminosae: Loteae): Partial incongruence between nrITS, nrETS and plastid markers and biogeographic implications. Taxon, 65(5), 997–1018. https://doi.org/10.12705/655.4

Kühn U 1993 Chenopodiaceae. Pages 253-281 *in* K Kubitzki, ed. The families and genera of vascular plants, vol II Magnoliid, Hamamelid and Caryophyllid families. Springer, Berlin.

Labat J-N, Beentje H. 2003. A new species of *Psiadia* (Compositae) from Mayotte. Kew Bulletin 58: 971-975.

Lammers TG. 1995. Transfer of the southern African species of *Lightfootia* nom. illeg., to *Wahlenbergia*. *Taxon* 44: 333–339.

Lammers, TG. 2000. Revision of Lobelia sect. *Tupa* (Campanulaceae: Lobelioideae). Sida 19: 87–110.

Lander NS. 1989. Apostates (Asteraceae: Astereae), a new genus from the South-eastern Polynesian Island of Rapa. Australian Journal of Botany 2: 129-133.

Landis MJ, Freyman WA, Baldwin BG 2018. Retracing the Hawaiian silversword radiation despite phylogenetic, biogeographic, and paleogeographic uncertainty. Evolution, 72, 2343-2359.

Lee J, BG Baldwin, LD Gottlieb 2002 Phylogeny of *Stephanomeria* and related genera (Compositae-Lactuceae) based on analysis of 18S-26S nuclear rDNA ITS and ETS sequences. Am J Bot 89: 160-168.

Lens F, P Caris, E Smets, L Serlet, S Jansen. 2005. Comparative wood anatomy of the primuloid clade (Ericales *s.l.*). Syst Bot 30: 162-182.

Lens F, I Groeninckx, E Smets, S Dessein 2009 Woodiness within the Spermacoceae-Knoxieae alliance (Rubiaceae): retention of the basal woody condition in Rubiaceae or recent innovation?

Lewis G, Schrire B, Mackinder B, Lock M. 2005. Legumes of the world. Royal Botanic Gardens, Kew, UK.

Lindqvist C, VA Albert. 2002. Origin of the Hawaiian endemic mints within North American *Stachys* (Lamiaceae). Am J Bot 89: 1709-1724.

Lindqvist C, Motley TJ, Jeffrey JJ, Albert VA. 2003. Cladogenesis and reticulation in the Hawaiian endemic mints (Lamiaceae). Cladistics 19: 480-495.

Liogier, H.A. 1997. Descriptive flora of Puerto Rico and adjacent Islands. Vol. 5. Editorial de la Universidad de Puerto Rico. Pp. 121-125.

Lorence, D. H. and S. Perlman. 2007. A new species of Cyrtandra (Gesneriaceae) from Hawai`i, Hawaiian Islands. Novon 17: 357-361.

Lorence DH, Wagner WL. 2011. Revision of *Kadua* (Rubiaceae) in the Marquesas Islands, French Polynesia, with description of the new species K. lichtlei. In: Lorence DH, Wagner WL (Eds) Botany of the Marquesas Islands: new taxa, combinations, and revisions. PhytoKeys 4: 125–138.

Lourteig A 1994 *Oxalis* L. subgénero *Thamnoxys* (Endl.) Reiche emend. Lourt. Bradea 7: 1-199.

Lourteig A 2000 *Oxalis* L. subgéneros *Monoxalis* (Small) Lourt, *Oxalis* y *Trifidus* Lourt. Bradea 7: 201-629.

Mabberley DJ 1974 The pachycaul lobelias of Africa and St Helena. Kew Bulletin 29: 535-584.

Maire R. 1967. Flore de l’Afrique du Nord. Vol 13. Rhoedales: Cruciferae, pp 6-365. Editions Paul Lechevalier, Paris.

Mandakova T, Pouch M, Harmanova K, Zhan SH, Mayrose I, Lysak MA 2017. Multispeed genome diploidization and diversification aftern an ancient allopolyploidization. Molecular Ecology, 26, 6445–6462.

Mansion G, L Struwe 2004 Generic delimitation and phylogenetic relationships within the subtribe Chironiinae (Chironieae: Gentianaceae), with special reference to *Centaurium*: evidence from nrDNA and cpDNA sequences. Mol Phyl Evol 32: 951-977.

Marais W 1970 Cruciferae. Pages 1-221 *in* LE Codd, B De Winter, DJB Killick, eds. Flora of Southern Africa, vol 13. Department of Agricultural Technical Services, South Africa.

Marais W 1984 Madagascan Nesogenes (Nesogenaceae). Kew Bull 38: 37-39.

Marcussen T, Meseguer AS. 2017. Species-level phylogeny, fruit evolution and diversification history of *Geranium* (Geraniaceae). Molecular phylogenetics and Evolution 110: 134-149.

McMullen CK 1999 Flowering plants of the Galapagos. Cornell University Press, Ithaca, New York, USA.

McVaugh R. 1943. Campanulaceae. North American Flora 32A: 1-110.

Mears JA 1982 A summary of *Blutaparon* Rafinesque including spieces earlier known as *Philoxerus* R.Brown (Amaranthaceae). Taxon 31: 111-117.

Meimberg H, T Abele, C Bräuchler C, JK McKay, PL Pérez de Paz, G Heubl 2006 Molecular evidence for adaptive radiation of *Micromeria* Benth. (Maliaceae) on the Canary Islands a inferred from chloroplast and nuclear DNA sequences and ISSR fingerprint data. Mol Phyl Evol 41: 566-578.

Mejias JA, Kim S-C 2012 Taxonomic treatment of Cichorieae (Asteraceae) endemic to the Juan Fernandez and Desventuradas Islands (SE Pacific). Ann Bot Fennici 49: 171-178.

Menezes de Sequeira M, Jardim R, Silva M, Carvalho L. 2007. Musschia isambertoi M. Seq., R. Jardim, M. Silva & L. Carvalho (Campanulaceae), a new species from the Madeira Archipelago (Portugal). Anales del Jardín Botánico de Madrid 64: 135-146.

Mesa A 1981 Nolanaceae. Flora Neotropica nr 26. New York Botanical Garden, New York.

Mesfin Tadesse 1993 An account for *Bidens* (Compositae: Heliantheae) for Africa. Kew Bull 48: 437-516.

Meusel H, Kästner A. 1975. *Carlina* L. Pages 597-602 *in* PH Davies, ed. Flora of Turkey and the East Aegian Islands, vol 5. Edinburgh University Press, Edinburgh.

Meve U, Laurente O, Alejandro GJ, Livshultz T. 2009: Systematics of *Clemensiella (Apocynaceae, Asclepiadoideae).* Edinburgh J. Bot. **66:** 447–457.

Michener DC 1983 Systematic and ecological wood anatomy of Californian Scrophulariaceae. I. *Antirrhinum*, *Castilleja*, *Galvezia*, and *Mimulus* sect. *Diplacus*. Aliso 10: 471-487.

Miller AL, Duncan RP. 2004. The impact of exotic weed competition on a rare New Zealand outcrop herb, *Pachycladon cheesemanii* (Brassicaceae). New Zealand Journal of Ecology 28: 113-124.

Moazzeni H, Zarre S, Pfeil BE, Bertrand YJK, German DA, Al-Shehbaz IA, Mummenhoff K, Oxelman B. 2014. Phylogenetic perspectives on diversification and character evolution in the species-rich genus *Erysimum* (Erysimeae; Brassicaceae) based on a densely sampled ITS approach. Botanical Journal of the Linnean Society 175: 497-522.

Moghaddam M, Kazempour Osaloo S, Hosseiny, Azimi F. 2017. Phylogeny and divergence times of the Coluteoid clade with special reference to *Colutea* (Fabaceae) inferred from nrDNA ITS and two cpDNAs, matK and rpl32-trnL(UAG) sequences data. Plant Biosystems 151: 1082-1093.

Moore JW 1933. New and critical plants from Raiatea. Bernice P. Bishop Museum Bulletin 102: 1-53.

Moore MJ, J Francisco-Ortega, A Santos-Guerra, RK Jansen 2002 Chloroplast DNA evidence for the roles of island colonization and extinction in *Tolpis* (Asteraceae: Lactuceae). Am J Bot 89: 518-526.

Moran, R. 1996. *The flora of Guadalupe Island, Mexico.* Memoirs of the California Academy of Sciences, Number 19. San Francisco: California Academy of Sciences.

Morawetz JJ, Randle CP, Wolfe AD. 2010. Phylogenetic relationships within the tropical clade of Orobanchaceae. Taxon 59: 416-426.

Mort ME, TR O'Leary, P Carillo-Reyes, T Nowell, JK Archibald, CP Randle. 2010. Phylogeny and evolution of Crassulaceae: Past, present and future. Schumannia 6: 69-86.

Mummenhoff K, Polster A, Mülhausen A, Theissen G. 2009. *Lepidium* as a model system for studying the evolution of fruit development in Brassicaceae. Journal of Experimental Botany 60: 1503-1513.

Mummenhoff K, H Brüggemann, JJ Bowman 2001 Chloroplast DNA phylogeny and biogeography of *Lepidium* (Brassicaceae). Am J Bot 88: 2051-2063.

Munz PA 1959 Californian flora. University of California Press, Berkeley and Los Angeles.

Möller M, Forrest A, Wei YG, Weber A 2011. A molecular phylogenetic assessment of the advanced Asiatic and Malesian didymocarpoid Gesneriaceae with focus on non-monophyletic and monotypic genera. Plant Syst Evol, 292.

Müller K, T Borsch 2005 Phylogenetics of Amaranthaceae based on matK/trnK sequence data – evidence from parsimony, likelihood, and Bayesian analyses. Ann Missouri Bot Gard 92: 66-102.

Navajas-Perez R, de la Herran R, Lopez Gonzalez G, Jamilena M, Lozano R, Ruiz Rejon C, Ruiz Rejon M, Garrido-Ramos A. 2005. The evolution of reproductive systems and sex-determining mechanisms within *Rumex* (Polygonaceae) inferred from nuclear and chloroplastidial sequence data. Molecular Biology and Evolution 22:1929-1939. 2005

Negron-Ortiz V, Watson LE. 2003. Hypotheses for the colonization of the Caribbean basin by two genera of the Rubiaceae: *Erithalis* and *Ernodea*. Syst. Bot. 28: 442-451.

Negron-Ortiz V, Hickey RJ 1996 The genus Ernodea (Rubiaceae) in the Caribbean Basin. II. Morphological analyses and systematics. Systematic Botany 21: 445-458.

Nesom GL. 1989. Infrageneric taxonomy of New World *Erigeron* (Compositae: Asteraceae). Phytologia 67: 67-93.

Neupane S, Lewis PO, Dessein S, Shanks H, Paudyal S, Lens F. 2017. Evolution of woody life form on tropical mountains in the tribe Spermacoceae (Rubiaceae). *American Journal of Botany* 104: 419-438.

Neupane S, Dessein S, Wikström N, Lewis PO, Long C, Bremer B, Motley TJ. 2015. The *Hedyotis-Oldenlandia* complex (Rubiaceae: Spermacoceae) in Asia and the Pacific: Phylogeny revisited with new generic delimitations. *Taxon* 64: 299–322.

Nishii K, Hughes M, Briggs M, Haston E, Christie F, DeVilliers MJ, Hanekom T, Roos WG, Bellstedt DU, Möller M. 2015. *Streptocarpus* redefined to include all Afro-Malagasy Gesneriaceae: Molecular phylogenies prove congruent with geographical distribution and basic chromosome numbers and uncover remarkable morphological homoplasies. Taxon 64: 1243-1274.

Nordenstam B. 1978. Taxonomic studies in the tribe Senecioneae. Opera Botanica 44: 3-83.

Nordenstam B. 2006. New genera and combinations in the Senecioneae of the Greater Antilles. Compositae Newsletter 44: 50-73.

Nürk N, Atchinson GW, Hughes CE 2019. Island woodiness underpins accelerated disparification in plant radiations. New Phytologist, 244, 518–531.

Nyffeler R 2003 *Aeonium*. Pages 15-23 *in* U Eggli, ed., Illustrated handbook of succulent plants: Crassulaceae. Springer, Berlin.

Oberprieler C, S Himmelreich, M Källersjö, J Vallès, LE Watson, R Vogt 2009 Chapter 38 Anthemideae. Pages 631-666 *in* VA Funk, A Sussana, TF Stuessy, RJ Bayer, eds. Systematics, evolution, and biogeography of Compositae. International Association for Plant Taxonomy, Vienna.

Ogundipe OT, Chase M. 2009. Phylogenetic analyses of Amaranthaceae based on matK DNA sequence data with emphasis on west African species. Turkish Journal of Botany 33: 153–161

Ojeda I, Santos-Guerra A, Jaén-Molina R, Oliva-Tejera F, Caujapé-Castells J, Cronk. (2012). The origin of bird pollination in Macaronesian Lotus (Loteae, Leguminosae). Molecular Phylogenetics and Evolution, 62(1), 306–318. https://doi.org/10.1016/j.ympev.2011.10.001

Oxelman B, P Kornhall, RG Olmstead, B Bremer 2005 Further disintegration of Scrophulariaceae. Taxon 54: 411-425.

Panero JL, RK Jansen, JA Clevinger 1999 Phylogenetic relationships of subtribe Ecliptinae (Asteraceae: Heliantheae) based on chloroplast DNA restriction site data. Am J Bot 66: 413-427.

Pax DL, RA Price, HJ Michaels 1997 Phylogentic position of the Hawaiian Geraniums based on *rbcL* sequences. Am J Bot 84: 72-78.

Pelser PB, B Nordenstam, JW Kadereit, LE Watson 2007 An ITS phylogeny of tribe Senecioneae (Asteraceae) and a new delimitation of *Senecio* L. Taxon 56: 1077-1104.

Perez de Paz PL 1978 Revision del genero Micromeria Bentham (Lamiaceae-Stachyoideae) en la region Macaronesica. Instituto de Estudios Canarios, Monografías 16: 1–306.

Plana V, Sands MJS, Beentje HJ. 2006. Begoniaceae. In: Beentje HJ, Ghanzafar SA, eds. Flora of tropical East Africa. Kew: Royal Botanic Gardens, 1–53.

Plana V 2003 Phylogenetic relationships of the Afro-Malagasy members of the large genus *Begonia* inferred from *trnL* intron sequences. Syst Bot 28: 693–704.

Polatschek A. 1976. Die Gattung *Erysimum* auf den Kapverden, Kanaren und Madeira. Am Naturhist Mus Wien 80: 93-103.

Polatschek A. 2011. Revision der Gattung *Erysimum* (Cruciferae), Teil 2: Georgien, Armenien, Azerbaidzan, Türkei, Syrien, Libanon, Israel, Jordanien, Irak, Iran, Afghanistan. Am Naturhist Mus Wien, B 112: 369-497.

Powell AM. 1974. Taxonomy of *Perityle* section *Perityle* (Compositae – Peritylinae). Rhodora 76: 229-306.

Press JR, MJ Short 1994 Flora of Madeira. The Natural History Museum, London.

Pruski J. F. 2015. Studies of Neotropical Compositae–X. Revision of the West Indian genus *Narvalina* (Coreopsideae). Phytoneuron 2015–31: 1–15.

Puglisi C, Middleton DJ, Triboun P, Möller M. 2011. New insights into the relationships between *Paraboea*, *Trisepalum*, and *Phylloboea* (Gesneriaceae) and their taxonomic consequences

Puppo P, Meimberg H. 2015. New species and new combinations in *Micromeria* (Lamiaceae) from the Canary Islands and Madeira. Phytotaxa 230: 1-21.

Rauscher JT 2002 Molecular phylogenetics of the *Espeletia* complex (Asteraceae): evidence from the nrDNA ITS sequences of the closest relatives of an Andean adaptive radiation. Am J Bot 89: 1074-1084.

Raven PH 1963 A flora of San Clemente Island, California. Aliso 5: 289-347.

Rechinger, K. H. 1982. Phlomis. – In: Rechinger, K. H. (ed.), Flora Iranica, no. 150. Akad. Druck-u. Verlagsanstalt, pp. 292–317.

Reifenberger U, Reifenberger A. 1992. Sonchus wildpretii (Compositae), ein neuer Endemit der Insel La Gomera (Kanarische Inseln). Willdenowia 22: 49-53.

Reitsma JM. 1984. Begonia section Bacca begonia Reitsma, sect. nov. Agricultural University Wageningen Papers 84: 95–111.

Richardson IBK. 1972. Scrophularia. – In: Tutin, T. G. et al. (eds), Flora Europaea. Vol. 3. Cambridge Univ. Press, pp. 216–220.

Richardson IBK 1979 A distinctive new species of *Wahlenbergia* (Campanulaceae from Mauritius. Kew Bull 33: 547-550.

Ridley HN 1905 The Gesneriaceae of the Malay Peninsula. Journal of the Straits Branch Royal Asiatic Society 44: 1-92.

Ridley HN 1923 The flora of the Malay Peninsula, vol 2, order 103 Gesneraceae, pp 495-547, L Reeve and Co, Ashford, Kent, UK.

Roalson EH, Boggan JK, Skog LE, Zimmer EA. 2005. Untangling Gloxinieae (Gesneriaceae). I. Phylogenetic patterns and generic boundaries inferred from nuclear, chloroplast and morphological cladistics datasets. Taxon 389-410.

Rodda M, Ercole E. 2014. *Hoya papaschonii* (Apocynaceae: Asclepiadoideae), a new species from southern Thailand with a peculiar corona. *Phytotaxa* 175: 97–106.

Rodda M, Simonsson Juhonewe N, Ercole E. 2013. Hoya corymbosa (Apocynaceae, Asclepiadoideae), a new unusual species from Sabah, Borneo, and its systematic position based on phylogenetic analysis. Systematic Botany 38: 1125-1131.

Rodda M, Simonsson N. 2011. *Hoya medinillifolia* (Apocynaceae Asclepiadoideae), a new species from lowland forests of Sarawak, Borneo. *Webbia* 66: 149–154.

Rodriguez RL 1957 Systematic anatomical studies on *Myrrhidendron* and other woody Umbellales. Univ Calif Publ Bot 29: 145-318.

Rollins RC 1970 Notes on *Streptanthus* and *Erysimum* (Cruciferae). Contrib Gray Herb 200: 190-195.

Rønsted N, MW Chase, DC Albach, MA Bello 2002 Phylogenetic relationships within *Plantago* (Plantaginaceae): evidence from nuclear ribosomal ITS and plastid *trnL-F* sequence data. Bot J Linnean Soc 139: 323-338.

Sanchez A, KA Kron 2008 Phylogenetics of Polygonaceae with an emphasis on the Evolution of Eriogonoideae. Syst Bot 33: 87-96.

Sanchez A, TM Schuster, KA Kron 2009 A large scale phylogeny of Polygonaceae based on molecular data. *Int. J. Plant Sci.* 170: 1044-1055.

Schaefer H 2002. Flora of the Azores. A filed guide. Margraf Verlag, Weikersheim, Germany, 264pp.

Schaefer H, C Heibl, SS Renner 2009 Gourds afloat: a dated phylogeny reveals an Asian origin of the gourd family (Cucurbitaceae) and numerous oversea dispersal events. Proc R Soc B 276: 843-851.

Schaefer H, SS Renner 2011. Cucurbitaceae. Pages 112-174 *in* K Kubitzki, ed. The families and genera of vascular plants. Springer, Berlin.

Scheunert A, Heubl G. 2014. Diversification of *Scrophularia* (Scrophulariaceae) in the western Mediterranean and Macaronesia – phylogenetic relationships, reticulate evolution and biogeographical patterns. Molecular Phylogenetics and Evolution 70: 296-313.

Schlechter R 1923 Gesneriaceae papuanae. Bot Jahrb Syst 58: 255-379.

Schuster TM, JL Reveal, MJ Bayly, KA Kron. 2015. An updated molecular phylogeny of Polygonoideae (Polygonaceae): Relationships of Oxygonum, Pteroxygonum, and Rumex, and a new circumscription of Koenigia. Taxon 64: 1188-1208.

Schuster TM, JL Reveal, KA Kron 2011 Phylogeny of Polygoneae (Polygonaceae: Polygonoideae). Taxon 60: 1653-1666.

Scott AJ. 1987. A second species of *Cylindrocline* (Compositae-Inuleae). Kew Bulletin 42: 476.

Scott AJ. 1991. Notes on Compositae-Astereae for the ‘Flore des Mascareignes’. Kew Bulletin 46: 339-353.

Sebastian P, Schaefer H, Lira R, Telford I, Renner SS. 2012. Radiation following long-distance dispersal: the contributions of time, opportunity and diaspore morphology in *Sicyos* (Cucurbitaceae). Journal of Biogeography 39: 1427–1438

Shannon RK, Wagner WL. 1997. *Oparanthus* (Asteraceae, subtribe Coreopsidinae) revisited. Allertonia 7: 273-295.

Sherff EE 1945 Revision of the genus *Schiedea* Cham. & Schlecht. Brittonia 5: 308-335.

Shu L 2003. Polygonum. In: Wu ZY, Raven PH, Hong DY, eds. Flora of China, vol. 5. Beijing: Science Press, 278–315.

Skog LE 1976 A study of the tribe Gesnerieae, with a revision of *Gesneria* (Gesneriaceae: Gesnerioideae) Smith Contr Bot 29: 1-182.

Skottsberg C 1921 The phanerogams of the Juan Fernandez Islands. Uppsala, Almquist and Wiksells Boktryckeri.

Skottsberg C 1957 Une seconde espèce de Centaurodendron Johow. Bull Jard Bot Etat Brux 27: 585-589.

Sloan DB, Oxelman B, Rautenberg A, Taylor DR 2010. Phylogenetic analysis of mitochondrial substitution rate variation in the angiosperm tribe Sileneae. BMC Evolutionary Biology, 10, 12.

Smith A. C. 1991. Flora Vitiensis Nova. A new flora of Fiji (spermatophytes only), Vol. 5. Angiospermae: Dicotyledones, Families 170–186; Monocotyledones, Family 32. Lawai, Kauai, Hawaii: National Tropical Botanical Garden.

Smith JF, SB Draper, LC Hileman, DA Baum. 2004. A phylogenetic analysis within tribes Gloxinieae and Gesnerieae (Gesnerioideae: Gesneriaceae). Syst Bot 29: 947-958.

Smith JF, JC Wolfram, KD Brown, CL Carroll, DS Denton. 1997. Tribal relationships in the Gesneriaceae: Evidence from DNA sequences of the chloroplast gene *ndh*F. Ann. Missouri Bot. Gard. 84: 50-66.

Smith, J. F. 2001. The phylogenetic relationships of *Lembocarpus* and *Goyazia* (Gesneriaceae) based on *ndh*F sequences. *Ann. Missouri Bot. Gard.* 88: 135-143.

Snogerup S, Gustafsson M, von Bothmer R 1990 Brassica sect Brassica (Brassicaceae) – I. Taxonomy and variation. Wildenowia 19: 271-365.

Snogerup S. 1967. Studies in the Aegean flora. VIII. Erysimum sect. Cheiranthus. A Taxonomy. Opera Botanica 13: 1-70.

Solbrig OT. 1962. The South American species of Erigeron. Contributions from the Gray Herbarium of Harvard University 191: 3-79.

Soza VL and Olmstead RG 2010 Molecular systematics of tribe Rubieae (Rubiaceae): Evolution of major clades, development of leaf-like whorls, and biogeography. Taxon 59: 755–771

Stefanović S, L Krueger, RG Olmstead 2002 Monophyly of the Convolvulaceae and circumscription of their major lineages based on DNA sequences of multiple loci. Am J Bot 89: 1510-1522.

Stern WL, GK Brizicky, RH Eyde 1969 Comparative wood anatomy and relationships of Columelliaceae. J Arnold Arb 50: 36-75.

Stiefelhagen H 1910 Systematische and pflanzengeograpfische Studies zur Kenntnis der Gattung Scrophularia. Bot. Jahrb. Syst. 44: 406-496.

Strijk JS, Noyes RD, Strasberg D, Cruaud C, Gavory F, Chase MW, Abbott RJ, Thébaud C. 2012. In and out of Madagascar: Dispersal to Peripheral Islands, Insular Speciation and Diversification of Indian Ocean Daisy Trees (Psiadia, Asteraceae). PLoS One 7: e42932.

Strother JL. 1977. Taxonomy of *Chrysactinia*, *Harnackia*, and *Lescaillea* (Compositae: Tageteae). Madrono 24: 129-139.

Struwe L, JW Kadereit, J Klackenberg, S Nilsson, M Thiv, KB von Hagen, VA Albert 2002 Systematics, character evolution, and biogeography of Gentianaceae, including a new tribal and subtribal classification. Pages 21-309 *in* L Struwe, VA Albert, eds. Gentianaceae. Systematics and natural history. Cambridge University Press, Cambridge.

Sun Y, Li YS, Vargas-Mendoza CF, Wang FG, Xing F 2016. Colonization and diversification of the Euphorbia species (Sect. Aphyllis subsect. Macaronesicae) on the Canary Islands. Scientific Reports, 6, 34454.

Tank DC, RG Olmstead 2008 From annuals to perennials: phylogeny of subtribe Castillejinae (Orobanchaceae). Am J Bot 95: 608-625.

Tao, C, and Ehrendorfer F. 2011. Rubia. *In* Z. Y. Wu, P. H. Raven, D. Y. Hong [eds.], Flora of China, vol. 19, pp. 305-319. Missouri Botanical Garden Press, St. Louis, Missouri, USA.

Tavakkoli S, Kazempour Osaloo SH, Mozaffarian V, Maassoumi A. 2015. Molecular phylogeny of Atraphaxis and the woody Polygonum species (Polygonaceae): taxonomic implications based on molecular and morphological evidence. Plant Systematics and Evolution 301:1157–1170

Taylor FH 1972 The secondary xylem of the Violaceae: a comparative study. Bot Gaz 133: 230-242.

Terrell, EE, Robinson, H, Wagner, WL & Lorence, DH. 2005. Resurrection of genus *Kadua* for Hawaiian Hedyotidinae (Rubiaceae), with emphasis on seed and fruit characters and notes on South Pacific species. *Syst. Bot.* 30: 818–833.

Thiv M, Thulin M, Kilian N, Linder HP. 2006 Eritreo*-*Arabian affinities of the Socotran flora as revealed from the molecular phylogeny of Aerva. (Amaranthaceae). Syst. Bot. 31: 560–570.

Thiv M, JW Kadereit 2002 Tribe Chironieae. Pages 108-137 *in* L Struwe, V Albert, eds. Gentianaceae: systematics and natural history. Cambridge University Press, Cambridge.

Thiv M, Struwe L, Kadereit JW 1999. The phylogenetic relationships and evolution of the Canarian laurel forest endemic Ixanthus viscosus (Aiton) Griseb. (Gentianaceae): Evidence from matK and ITS sequences, and floral morphology and anatomy. Plant Systematics and Evolution, 218(3–4), 299–317. https://doi.org/10.1007/BF01089233

Thomas DC, Weigend M, Hilger HH. 2008. Phylogeny and systematics of Lithodora (Boraginaceae-Lithospermeae) and its affinities ro the monotypic genera Mairetis, Halacsya and Paramoltkia based on ITS1 and trnL sequence data and morphology. Taxon 57: 79-97.

Thomas DC, M Hughes, T Phutthai, WH Ardi, S Rajbhandary, R Rubite, AD Twyford, JE Richardson 2011 West to east dispersal and subsequent rapid diversification of the mega-diverse genus *Begonia* (Begoniaceae) in the Malesian archipelago. J Biogeogr 39: 98-113.

Thompson IR 2015. Senecio, in Flora of Australia vol 37, Asteraceae I, (Wilson A, ed.), CSIRO Publishing, Melbourne, Australia, pp. 209-307.

Thulin M, Moore AJ, El-Seedi H, Larsson A, Christin P-A, Edwards EJ. 2016. Phylogeny and generic delimitation in Molluginaceae, new pigment data in Caryophyllales, and the new family Corbichoniaceae. Taxon 65: 775-793.

Thulin M 1980 *Nesocodon*, a new genus in Campanulaceae. Kew Bull 34: 813-814.

Thulin, M. 1984. Lobeliaceae. In: Polhill, R.M. (ed.) Flora of tropical East Africa. Rotterdam: Balkema, pp1-60.

Townsend CC 1993 Amaranthaceae. Pages 70-91 *in* K Kubitzki, ed. The families and genera of vascular plants, vol II Magnoliid, Hamamelid and Caryophyllid families. Springer, Berlin.

Tran TB, Kim JH, Kim DK, Lee J, Ha BT, Simonsson Juhonewe N, Rodda M. 2011. *Hoya ignorata* (Apocynaceae, Asclepiadoideae) an overlooked species widely distributed across southeast Asia. Novon 21: 508–514.

Turini FG, C Bräuchler, G Heubl 2010 Phylogenetic relationships and evolution of morphological characters in *Ononis* L. (Fabaceae). Taxon 59: 1077-1090.

Tye A. 2010. The Galapagos endemic Darwiniothamnus alternifolius (Asteraceae, Astereae) transferred to Erigeron. Novon 20: 112

Upson T, S Andrews 2004 The genus *Lavandula*. The Royal Botanic Gardens, Kew.

Urban I 1910 Zwei neue Loasaceen von Sto. Domingo. Berichte der Deutsche Botanische Gesellschaft 28: 515-523.

van der Walt, JJA 1977. Pelargoniums of Southern Africa, vol. 1. Purnell and Sons SA Ltd, Cape Town.

van der Walt, JJA & Vorster, PJ. 1981. Pelargoniums of Southern Africa, vol. 2. Juta, Cape Town.

van der Walt, JJA & Vorster, PJ. 1988. Pelargoniums of Southern Africa, vol. 3. National Botanic Gardens, Kirstenbosch.

Verdcourt B 1989 Rubiaceae. In Flora Zambesiaca vol 5.1. (ed. E. Launert), Spermacoce, pp. 165-188. Royal Botanical Gardens Kew, UK.

Villa I, Fernaandez de Castro AG, Fuertes-Aguilar J, Nieto G 2018. Out of North Africa by different routes. Phylogeography and species distribution model of the western Mediterranean Lavatera maritima (Malvaceae). Botanical Journal of the Linnean Society, 187, 441–455.

Villaverde T, Pokorny L, Olsson S, Rincón-Barrado M, Johnson MG, Gardner EM, Wickett NJ, Molero J, Riina R, Sanmartín I 2018. Bridging the micro- and macroevolutionary levels in phylogenomics: Hyb-Seq solves relationships from populations to species and above. New Phytologist, 220, 636–650.

Wagenitz G 1975 *Centaurea* L. Pages 465-584 *in* PH Davies, ed. Flora of Turkey and the East Aegian Islands, vol 5. Edinburgh University Press, Edinburgh.

Wagner, WL., Weller, SG, Sakai, AK. 1994. Description of a rare new cliff-dwelling species from Kaua`i, Schiedea attenuata (Caryophyllaceae). Novon 4: 187-190.

Wagner WL, Clark JR, Lorence DH. 2014. Revision of endemic Marquesas Islands *Bidens* (Asteraceae, Coreopsideae). PhytoKeys 38: 37–67.

Wagner WL, Lorence DH. 2011. Two new Marquesan species of the southeastern Polynesian genus *Oparanthus* (Asteraceae, Coreopsidinae). In: Lorence DH, Wagner WL (Eds) Botany of the Marquesas Islands: new taxa, combinations, and revisions. PhytoKeys 4: 139–148.

Wagner WL, Wagner AJ, Lorence DH 2013. Revision of *Cyrtandra* (Gesneriaceae) in the Marquesas Islands. Phytokeys 30: 33-64.

Wagner WL, DR Herbst, SH Sohmer. 1990. Manual of the flowering plants of Hawaii, vols. 1 and 2. University of Hawaii Press and Bishop Museum Press, Honolulu, Hawaii, USA.

Wagner WL, Herbst DR, Khan N, Flynn T. 2012. Hawaiian vascular plant updates: a supplement to the manual of the flowering plants of Hawai’I and Hawai’I’s ferns and fern allies. Version 1.3, downloaded online

Wagner WL. 1999. A New Species of Hawaiian Phyllostegia (Lamiaceae) from Kaua'i and recognition of a Wai'anae Mountain, O'ahu, endangered variety of *Phyllostegia parviflora*. Novon 9: 280-283

Wagstaff, S.J. & Hennion, F. 2007. Evolution and biogeography of *Lyallia* and *Hectorella* (Portulacaceae), geographically isolated sisters from the Southern Hemisphere. *Antarc. Sci.* 19: 417–426.

Wahrmund U, H Heklau, M Röser, A Kärstner, E Vitek, F Ehrendorfer, KB von Hagen 2010 A molecular phylogeny reveals frequent changes of growth form in *Carlina* (Asteraceae) Taxon 59: 367-378.

Walker JB, Sytsma KJ, J Treutlein, M Wink 2004 *Salvia* (Lamiaceae) is not monophyletic: implications for the systematics, radiation, and ecological specilializations of *Salvia* and tribe Mentheae. Am J Bot 91: 1115-1125.

Wang W, P Kaiyu, L Zhanyu, AL Weitzman, L.E. Skog 1998 Gesneriaceae. Pages 244-401 *in* W Zheng-yi, PH Raven, eds., Flora of China, vol 18. Science Press, Beijng.

Wanntorp L, Meve U. 2011. New combinations in *Hoya* for the species of *Clemensiella* (Marsdenieae, Apocynaceae). Willdenowia 41: 97-99.

Warwick SI, CA Sauder 2005 Phylogeny of tribe Brassiceae (Brassicaceae) based on chloroplast restriction site polymorphisms and nuclear ribosomal internal transcribed spacer and chloroplast *trn*L intron sequences. Can J Bot 83: 467-483.

Warwick SI, K Mummenhoff, CA Saunder, MA Koch, IA Al-Shehbaz 2010 Closing the gaps: phylogenetic relationships in the Brassicaceae based on DNA sequence data of nuclear ribosomal ITS region. Plant Syst Evol 285: 209-232.

Warwick SI, CA Sauder, IA Shehbaz 2008 Phylogenetic relationships in the tribe Alysseae (Brassicaceae) based on nuclear ribosomal ITS DNA sequences. Botany 86: 315-336**.**

Weber A 2004 Gesneriaceae. Pages 63-162 *in* K Kubitzki, ed. The families and genera of vascular plants, vol VII Lamiales (except Acanthaceae including Avicenniaceae). Springer, Berlin.

Weigend M 2006 validating subfamily, genus and species names in Loasaceae (Cornales). Taxon 55: 463-468.

Weigend M, Gottschling M, Hoot S, Ackermann M. 2004. A preliminary phylogeny of Loasaceae subfam. Loasoideae (Angiospermae: Cornales) based on trnL(UAA) sequence data, with consequences for systematics and historical biogeography. Organisms, Diversity & Evolution 4: 73-90

Weller SG, WL Wagner, AK Sakai 1995 A phylogenetic analysis if *Schiedea* and *Alsinidendron* (Caryophyllaceae: Alsinoideae): implications for the evoluation of breeding systems. Syst Bot 20: 315-337.

Wheeler JR 1987 Malvaceae pp 141-147. In Marchant NG, Wheeler JR, Rye BL, Bennett EM, Lander NS, Macfarlane TD (eds), Flora of the Perth Region, part 1, Western Australian Herbarium, Western Australia.

Wiggins, D., IL; Porter. (1971). Flora of the Galapagos Islands. In Flora of the Galapagos Islands. Standford: Standford University Press.

Wikström, N., Neupane, S., Kårehed, J., Motley, T.J. & Bremer, B. 2013. Phylogeny of *Hedyotis* L. (Rubiaceae: Spermacoceae): Redefining a complex Asian-Pacific assemblage. *Taxon* 62: 357–374.

Wilbur, R. L. 1991. Synopsis of the Mexican and Central American representatives of Lobelia section *Tylomium* (Campanulaceae: Lobelioideae). Sida 14: 555–567.

Wilson PG 1983 A taxonomic revision of the tribe Chenopodieae (Chenopodiaceae) in Australia. *Nuytsia* 4: 135-262.

Wimmer FE. 1948. Vorarbeiten zur Monographie der Campanulaceae-Lobelioideae: II Trib. Lobelieae. Ann. Naturhist. Mus. Wien 56: 317-374.

Wu Z-Y, Liu J, Provan J, Wang H, Chen C.-J, Cadotte MW, Luo YH, Amorim BS, Li D.-Z, Milne RI 2018. Testing Darwin’s transoceanic dispersal hypothesis for the inland nettle family (Urticaceae). Ecology Letters, 21, 1515–1529.

Wörz A 2011 Revision of *Eryngium* L. (Apiaceae-Sanuculoideae): general part and palaearctic species. Bibl Bot 159: 1-498.

Xu Z, Burtt BL, Skog LE, Middleton DJ 2008 A revision of *Paraboea* (Gesneriaceae). Edinb J Bot 65: 161-347.

Xu L, Choi B-H. 2010. Fabaceae. Flora of China, vol 10, pp 512-514 (eds. Zhengyi W, Raven PH, Deyuan H), Science Press (Beijing), Missouri Bot Gard Press (St Louis).

Yang Y, PE Berry 2011 Phylogenetics of the *Chamaesyce* clade (*Euphorbia*, Euphorbiaceae): reticulate evolution and long-distance dispersal in a prominent C4 lineage. Am J Bot 98: 1486-1503.

Yang Y, Morden CW, Sporck-Koehler MJ, Sack L, Wagner WL, Berry PE 2018. Repeated range expansion and niche shift in a volcanic hotspot archipelago: Radiation of C4 Hawaiian Euphorbia subgenus Chamaesyce (Euphorbiaceae). Ecology and Evolution, 8, 8523–8536.

Yurtseva OV, Kuznetsova OI, Mavrodieva ME, Mavrodiev EV. (2016) What is *Atraphaxis* L. (Polygonaceae, Polygoneae): cryptic taxa and resolved taxonomic complexity instead of the formal lumping and the lack of morphological synapomorphies. *PeerJ* 4:e1977

Zacharias EH, BG Baldwin 2010 A molecular phylogeny of North American Atripliceae (Chenopodiaceae), with implications for floral and photosynthethic pathway evolution. Syst Bot 35: 839-857.

Zaini NH. 2015. Stem anatomy and growth habit evolution of Asian *Begonia* (Begoniaceae). MSc-thesis, Naturalis Biodiversity Center, the Netherlands, and Singapore Botanical Gardens, Singapore.

Zohary M 1972 Ononis pp 113-122 in Flora Palestina part 2, The Israel Academy of Sciences and Humanities, Jerusalem.

Zuloaga FO, Morrone O, Begrano MJ. 2008. Catalago de las plantas vasculares del Cono Sur. Vol 2. Monographs in Systematic Botany from the Missouri Botanical Garden 107: 985-2286.

### Age of insular woody clades

Affenzeller, M., Kadereit, J. W., & Comes, H. P. (2018). Parallel bursts of recent and rapid radiation in the Mediterranean and Eritreo-Arabian biodiversity hotspots as revealed by Globularia and Campylanthus (Plantaginaceae). Journal of Biogeography, 45(3), 552–566. https://doi.org/10.1111/jbi.13155

Ahlstrand, N. I., Verstraete, B., Hassemer, G., Dunbar-Co, S., Hoggard, R., Meudt, H. M., & Rønsted, N. (2019). Ancestral range reconstruction of remote oceanic island species of Plantago (Plantaginaceae) reveals differing scales and modes of dispersal. Journal of Biogeography, 46(4), 706–722. https://doi.org/10.1111/jbi.13525

Antonelli, A. (2009). Have giant lobelias evolved several times independently? Life form shifts and historical biogeography of the cosmopolitan and highly diverse subfamily Lobelioideae (Campanulaceae). BMC Biology, 7(1), 82. https://doi.org/10.1186/1741-7007-7-82

Baldwin, B. G., & Wood, K. R. (2016). Origin of the Rapa endemic genus Apostates: Revisiting major disjunctions and evolutionary conservatism in the Bahia alliance (Compositae: Bahieae). Taxon, 65(5), 1064–1080. https://doi.org/10.12705/655.8

Bartish, I. V., Aïnouche, A., Jia, D., Bergstrom, D., Chown, S. L., Winkworth, R. C., & Hennion, F. (2012). Phylogeny and colonization history of Pringlea antiscorbutica (Brassicaceae), an emblematic endemic from the South Indian Ocean Province. Molecular Phylogenetics and Evolution, 65(2), 748–756. https://doi.org/10.1016/j.ympev.2012.07.023

Bruyns, P. V., Klak, C., & Hanáček, P. (2011). Age and diversity in Old World succulent species of Euphorbia (Euphorbiaceae). TAXON, 60(6), 1717–1733. https://doi.org/10.1002/tax.606016

Carine, M. A., Russell, S. J., Santos-Guerra, A., & Francisco-Ortega, J. (2004). Relationships of the Macaronesian and Mediterranean floras: Molecular evidence for multiple colonizations into Macaronesia and back-colonization of the continent in Convolvulus (Convolvulaceae). American Journal of Botany, 91(7), 1070–1085. https://doi.org/10.3732/ajb.91.7.1070

Chen, L. Y., Wang, Q. F., & Renner, S. S. (2016). East Asian lobelioideae and ancient divergence of a giant rosette Lobelia in Himalayan Bhutan. Taxon, 65(2), 293–304. https://doi.org/10.12705/652.6

Francisco-Ortega, J., Barber, J. C., Santos-Guerra, A., Febles-Hernández, R., & Jansen, R. K. (2001). Origin and Evolution of the Endemic Genera of Gonosperminae (Asteraceae: Anthemideae) from the Canary Islands: Evidence from Nucleotide Sequences of the Internal Transcribed Spacers of the Nuclear Ribosomal DNA. American Journal of Botany, 88(1), 161–169. https://doi.org/10.2307/2657136

Francisco-Ortega, J., Santos-Guerra, A., Mosa-Coello, R., González-Feria, E., & Crawford, D. J. (1996). Genetic resource conservation of the endemic genus Argyranthemum Sch. Bip. (Asteraceae: Anthemideae) in the Macaronesian Islands. Genetic Resources and Crop Evolution, 43(1), 33–39. https://doi.org/10.1007/BF00126938

Franzke, A., Sharif Samani, B.-R., Neuffer, B., Mummenhoff, K., & Hurka, H. (2017). Molecular evidence in Diplotaxis (Brassicaceae) suggests a Quaternary origin of the Cape Verdean flora. Plant Systematics and Evolution, 303(4), 467–479. https://doi.org/10.1007/s00606-016-1384-5

García-Verdugo, C., Caujapé-Castells, J., Mairal, M., & Monroy, P. (n.d.). How repeatable is microevolution on islands? Patterns of dispersal and colonization-related plant traits in a phylogeographical context. 12.

Givnish, T. J., Millam, K. C., Mast, A. R., Paterson, T. B., Theim, T. J., Hipp, A. L., Henss, J. M., Smith, J. F., Wood, K. R., & Sytsma, K. J. (2009). Origin, adaptive radiation and diversification of the Hawaiian lobeliads (Asterales: Campanulaceae). Proceedings of the Royal Society B: Biological Sciences, 276(1656), 407–416. https://doi.org/10.1098/rspb.2008.1204

Guzmán, B., Heleno, R., Nogales, M., Simbaña, W., Traveset, A., & Vargas, P. (2017). Evolutionary history of the endangered shrub snapdragon (Galvezia leucantha) of the Galápagos Islands. Diversity and Distributions, 23(3), 247–260. https://doi.org/10.1111/ddi.12521

Havran, J. C., Sytsma, K. J., & Ballard, H. E. (2009). Evolutionary relationships, interisland biogeography, and molecular evolution in the Hawaiian violets (Viola: Violaceae). American Journal of Botany, 96(11), 2087–2099. https://doi.org/10.3732/ajb.0900021

Huang, X., Deng, T., Moore, M. J., Wang, H., Li, Z., Lin, N., Yusupov, Z., Tojibaev, K. Sh., Wang, Y., & Sun, H. (2019). Tropical Asian Origin, boreotropical migration and long-distance dispersal in Nettles (Urticeae, Urticaceae). Molecular Phylogenetics and Evolution, 137, 190–199. https://doi.org/10.1016/j.ympev.2019.05.007

Huysduynen, A. H. van, Janssens, S., Merckx, V., Vos, R., Valente, L., Zizka, A., Larter, M., Karabayir, B., Maaskant, D., Witmer, Y., Fernández-Palacios, J. M., Nascimento, L. de, Jaén, R. M., Castells, J. C., Marrero-Rodríguez, Á., Arco, M. del, & Lens, F. (2020). Multiple origins of insular woodiness on the Canary Islands are consistent with palaeoclimatic aridification. BioRxiv, 2020.05.09.084582. https://doi.org/10.1101/2020.05.09.084582

Johnson, M. A., Pillon, Y., Sakishima, T., Price, D. K., & Stacy, E. A. (2019). Multiple colonizations, hybridization and uneven diversification in Cyrtandra (Gesneriaceae) lineages on Hawai’i Island. Journal of Biogeography, 46(6), 1178–1196. https://doi.org/10.1111/jbi.13567

Jones, K. E., Reyes-Betancort, J. A., Hiscock, S. J., & Carine, M. A. (2014). Allopatric diversification, multiple habitat shifts, and hybridization in the evolution of Pericallis (Asteraceae), a Macaronesian endemic genus. American Journal of Botany, 101(4), 637–651. https://doi.org/10.3732/ajb.1300390

Kim, S.-C., McGowen, M. R., Lubinsky, P., Barber, J. C., Mort, M. E., & Santos-Guerra, A. (2008). Timing and Tempo of Early and Successive Adaptive Radiations in Macaronesia. PLOS ONE, 3(5), e2139. https://doi.org/10.1371/journal.pone.0002139

Knope, M. L., Morden, C. W., Funk, V. A., & Fukami, T. (2012). Area and the rapid radiation of Hawaiian Bidens (Asteraceae). Journal of Biogeography, 39(7), 1206–1216. https://doi.org/10.1111/j.1365-2699.2012.02687.x

Knox, E. B., & Li, C. (2017). The East Asian origin of the giant lobelias. American Journal of Botany, 104(6), 924–938. https://doi.org/10.3732/ajb.1700025

Kondraskov, P., Schütz, N., Schüßler, C., Sequeira, M. M. de, Guerra, A. S., Caujapé-Castells, J., Jaén-Molina, R., Marrero-Rodríguez, Á., Koch, M. A., Linder, P., Kovar-Eder, J., & Thiv, M. (2015). Biogeography of Mediterranean Hotspot Biodiversity: Re-Evaluating the ‘Tertiary Relict’ Hypothesis of Macaronesian Laurel Forests. PLOS ONE, 10(7), e0132091. https://doi.org/10.1371/journal.pone.0132091

Landis, M. J., Freyman, W. A., & Baldwin, B. G. (2018). Retracing the Hawaiian silversword radiation despite phylogenetic, biogeographic, and paleogeographic uncertainty. Evolution, 72(11), 2343–2359. https://doi.org/10.1111/evo.13594

Mandáková, T., Pouch, M., Harmanová, K., Zhan, S. H., Mayrose, I., & Lysak, M. A. (2017). Multispeed genome diploidization and diversification after an ancient allopolyploidization. Molecular Ecology, 26(22), 6445–6462. https://doi.org/10.1111/mec.14379

Mansion, G., & Struwe, L. (2004). Generic delimitation and phylogenetic relationships within the subtribe Chironiinae (Chironieae: Gentianaceae), with special reference to Centaurium: Evidence from nrDNA and cpDNA sequences. Molecular Phylogenetics and Evolution, 32(3), 951–977. https://doi.org/10.1016/j.ympev.2004.03.016

Marcussen, T., & Meseguer, A. S. (2017). Species-level phylogeny, fruit evolution and diversification history of Geranium (Geraniaceae). Molecular Phylogenetics and Evolution, 110, 134–149. https://doi.org/10.1016/j.ympev.2017.03.012

Moazzeni, H., Zarre, S., Pfeil, B. E., Bertrand, Y. J. K., German, D. A., Al-Shehbaz, I. A., Mummenhoff, K., & Oxelman, B. (2014). Phylogenetic perspectives on diversification and character evolution in the species-rich genus Erysimum (Erysimeae; Brassicaceae) based on a densely sampled ITS approach. Botanical Journal of the Linnean Society, 175(4), 497–522. https://doi.org/10.1111/boj.12184

Navarro-Pérez, M. L., Vargas, P., Fernández-Mazuecos, M., López, J., Valtueña, F. J., & Ortega-Olivencia, A. (2015). Multiple windows of colonization to Macaronesia by the dispersal-unspecialized Scrophularia since the Late Miocene. Perspectives in Plant Ecology, Evolution and Systematics, 17(4), 263–273. https://doi.org/10.1016/j.ppees.2015.05.002

Neupane, S., Lewis, P. O., Dessein, S., Shanks, H., Paudyal, S., & Lens, F. (2017). Evolution of woody life form on tropical mountains in the tribe Spermacoceae (Rubiaceae). American Journal of Botany, 104(3), 419–438. https://doi.org/10.3732/ajb.1600248

Nürk, N. M., Atchison, G. W., & Hughes, C. E. (2019). Island woodiness underpins accelerated disparification in plant radiations. New Phytologist, 224(1), 518–531. https://doi.org/10.1111/nph.15797

Nylinder, S., Razafimandimbison, S. G., & Anderberg, A. A. (2016). From the Namib around the world: Biogeography of the Inuleae–Plucheinae (Asteraceae). Journal of Biogeography, 43(9), 1705–1716. https://doi.org/10.1111/jbi.12764

Ojeda, I., Santos-Guerra, A., Jaén-Molina, R., Oliva-Tejera, F., Caujapé-Castells, J., & Cronk, Q. C. B. (2012). The origin of bird pollination in Macaronesian Lotus (Loteae, Leguminosae). Molecular Phylogenetics and Evolution, 62(1), 306–318. https://doi.org/10.1016/j.ympev.2011.10.001

Pax, D. L., Price, R. A., & Michaels, H. J. (1997). Phylogenetic Position of the Hawaiian Geraniums Based on rbcL Sequences. American Journal of Botany, 84(1), 72–78. https://doi.org/10.2307/2445884

Pelser, P. B., Kennedy, A. H., Tepe, E. J., Shidler, J. B., Nordenstam, B., Kadereit, J. W., & Watson, L. E. (2010). Patterns and causes of incongruence between plastid and nuclear Senecioneae (Asteraceae) phylogenies. American Journal of Botany, 97(5), 856–873. https://doi.org/10.3732/ajb.0900287

Puppo, P., Curto, M., Velo-Antón, G., Paz, P. L. P. de, & Meimberg, H. (2014). The influence of geological history on diversification in insular species: Genetic and morphological patterns of Micromeria Benth. (Lamiaceae) in Tenerife (Canary archipelago). Journal of Biogeography, 41(10), 1871–1882. https://doi.org/10.1111/jbi.12354

Roalson, E. H., & Roberts, W. R. (2016). Distinct Processes Drive Diversification in Different Clades of Gesneriaceae. Systematic Biology, 65(4), 662–684. https://doi.org/10.1093/sysbio/syw012

Rønsted, N., Chase, M. W., Albach, D. C., & Bello, M. A. (2002). Phylogenetic relationships within Plantago (Plantaginaceae): Evidence from nuclear ribosomal ITS and plastid trnL-F sequence data. Botanical Journal of the Linnean Society, 139(4), 323–338. https://doi.org/10.1046/j.1095-8339.2002.00070.x

Schaefer, H., Heibl, C., & Renner, S. S. (2009). Gourds afloat: A dated phylogeny reveals an Asian origin of the gourd family (Cucurbitaceae) and numerous oversea dispersal events. Proceedings of the Royal Society B: Biological Sciences, 276(1658), 843–851. https://doi.org/10.1098/rspb.2008.1447

Schneider, A. C., & Moore, A. J. (2017). Parallel Pleistocene amphitropical disjunctions of a parasitic plant and its host. American Journal of Botany, 104(11), 1745–1755. https://doi.org/10.3732/ajb.1700181

Schüßler, C., Bräuchler, C., Reyes-Betancort, J. A., Koch, M. A., & Thiv, M. (2019). Island biogeography of the Macaronesian Gesnouinia and Mediterranean Soleirolia (Parietarieae, Urticaceae) with implications for the evolution of insular woodiness. TAXON, 68(3), 537–556. https://doi.org/10.1002/tax.12061

Schüssler, C., Freitag, H., Koteyeva, N., Schmidt, D., Edwards, G., Voznesenskaya, E., & Kadereit, G. (2017). Molecular phylogeny and forms of photosynthesis in tribe Salsoleae (Chenopodiaceae). Journal of Experimental Botany, 68(2), 207–223. https://doi.org/10.1093/jxb/erw432

Sebastian, P., Schaefer, H., Lira, R., Telford, I. R. H., & Renner, S. S. (2012). Radiation following long-distance dispersal: The contributions of time, opportunity and diaspore morphology in Sicyos (Cucurbitaceae). Journal of Biogeography, 39(8), 1427–1438. https://doi.org/10.1111/j.1365-2699.2012.02695.x

Sloan, D. B., Oxelman, B., Rautenberg, A., & Taylor, D. R. (2009). Phylogenetic analysis of mitochondrial substitution rate variation in the angiosperm tribe Sileneae. BMC Evolutionary Biology, 9(1), 260. https://doi.org/10.1186/1471-2148-9-260

Strijk, J. S., Noyes, R. D., Strasberg, D., Cruaud, C., Gavory, F., Chase, M. W., Abbott, R. J., & Thébaud, C. (2012). In and out of Madagascar: Dispersal to Peripheral Islands, Insular Speciation and Diversification of Indian Ocean Daisy Trees (Psiadia, Asteraceae). PLOS ONE, 7(8), e42932. https://doi.org/10.1371/journal.pone.0042932

Sun, Y., Li, Y., Vargas-Mendoza, C. F., Wang, F., & Xing, F. (2016). Colonization and diversification of the Euphorbia species (sect. Aphyllis subsect. Macaronesicae ) on the Canary Islands. Scientific Reports, 6(1), 34454. https://doi.org/10.1038/srep34454

Villa-Machío, I., Fernández de Castro, A. G., Fuertes-Aguilar, J., & Nieto Feliner, G. (2018). Out of North Africa by different routes: Phylogeography and species distribution model of the western Mediterranean Lavatera maritima (Malvaceae). Botanical Journal of the Linnean Society, 187(3), 441–455. https://doi.org/10.1093/botlinnean/boy025

Villaverde, T., Pokorny, L., Olsson, S., Rincón-Barrado, M., Johnson, M. G., Gardner, E. M., Wickett, N. J., Molero, J., Riina, R., & Sanmartín, I. (2018). Bridging the micro- and macroevolutionary levels in phylogenomics: Hyb-Seq solves relationships from populations to species and above. New Phytologist, 220(2), 636–650. https://doi.org/10.1111/nph.15312

Vitales, D., García-Fernández, A., Garnatje, T., Vallès, J., Cowan, R. S., Fay, M. F., & Pellicer, J. (2015). Conservation genetics of the rare Iberian endemic Cheirolophus uliginosus (Asteraceae). Botanical Journal of the Linnean Society, 179(1), 157–171. https://doi.org/10.1111/boj.12302

Willyard, A., Wallace, L. E., Wagner, W. L., Weller, S. G., Sakai, A. K., & Nepokroeff, M. (2011). Estimating the species tree for Hawaiian Schiedea (Caryophyllaceae) from multiple loci in the presence of reticulate evolution. Molecular Phylogenetics and Evolution, 60(1), 29–48. https://doi.org/10.1016/j.ympev.2011.04.001

Wu, Z.-Y., Liu, J., Provan, J., Wang, H., Chen, C.-J., Cadotte, M. W., Luo, Y.-H., Amorim, B. S., Li, D.-Z., & Milne, R. I. (2018). Testing Darwin’s transoceanic dispersal hypothesis for the inland nettle family (Urticaceae). Ecology Letters, 21(10), 1515–1529. https://doi.org/10.1111/ele.13132

Yan, H.-F., Zhang, C.-Y., Anderberg, A. A., Hao, G., Ge, X.-J., & Wiens, J. J. (2018). What explains high plant richness in East Asia? Time and diversification in the tribe Lysimachieae (Primulaceae). New Phytologist, 219(1), 436–448. https://doi.org/10.1111/nph.15144

Yang, Y., Morden, C. W., Sporck-Koehler, M. J., Sack, L., Wagner, W. L., & Berry, P. E. (2018). Repeated range expansion and niche shift in a volcanic hotspot archipelago: Radiation of C4 Hawaiian Euphorbia subgenus Chamaesyce (Euphorbiaceae). Ecology and Evolution, 8(16), 8523–8536. https://doi.org/10.1002/ece3.4354

### Occurrence of large fossil herbivores

Abbazzi, L., Delfino, M., Gallai, G., Trebini, L., & Rook, L. (2008). New data on the vertebrate assemblage of Fiume Santo (North‐West Sardinia, Italy), and overview on the Late Miocene Tusco‐Sardinian palaeobioprovince. Palaeontology, 51(2), 425–451.

Adams, S. J., McDowell, M. C., & Prideaux, G. J. (2016). Understanding accumulation bias in the ecological interpretation of archaeological and paleontological sites on Kangaroo Island, South Australia. Journal of Archaeological Science: Reports, 7, 715–729.

Angelone, C., Čermák, S., Moncunill-Solé, B., Quintana, J., Tuveri, C., Arca, M., & Kotsakis, T. (2018). Systematics and paleobiogeography of Sardolagus obscurus. Journal of Paleontology, 92(a), 506–522.

Athanassiou, A., Herridge, V., Reese, D. S., Iliopoulos, G., Roussiakis, S., Mitsopoulou, V., Tsiolakis, E., & Theodorou, G. (2015). Cranial evidence for the presence of a second endemic elephant species on Cyprus. Quaternary International, 379, 47–57.

Athanassiou, A., Van der Geer, A. A. E., & Lyras, G. A. (2019). Pleistocene insular Proboscidea of the Eastern Mediterranean: A review and update. Quaternary Science Reviews, 218, 306–321.

; Bates, M., Pope, M., Shaw, A., Scott, B. and Schwenninger, J.L. (2013). Late Neanderthal occupation in North-West Europe: rediscovery, investigation and dating of a last glacial sediment sequence at the site of La Cotte de Saint Brelade, Jersey. Journal of Quaternary Science, 28(7), pp.647-652.

Bergh, G. D. (1999). The Late Neogene elephantoid-bearing faunas of Indonesia and their palaeozoogeographic implications. Scripta Geologica, 117, 1–419.

Bergh, G. D., Vos, J., & Sondaar, P. Y. (2001). The Late Quaternary palaeogeography of mammal evolution in the Indonesian Archipelago. Palaeogeography, Palaeoclimatology, Palaeoecology, 171(3–4), 385–408.

Black, K. H., Archer, M., Hand, S. J., & Godthelp, H. (2012). The rise of Australian marsupials: A synopsis of biostratigraphic, phylogenetic, palaeoecologic and palaeobiogeographic understanding. In Earth and life (pp. 983–1078). Springer.

Boulbes, N., & Van Asperen, E. N. (2019). Biostratigraphy and palaeoecology of European Equus. Frontiers in Ecology and Evolution, 7, 301.

Bover, P., Quintana, J., & Alcover, J. A. (2008). Three islands, three worlds: Paleogeography and evolution of the vertebrate fauna from the Balearic Islands. Quaternary International, 182(1), 135–144.

Bover, P., Quintana, J., & Alcover, J. A. (2010). A new species of Myotragus Bate, 1909 (Artiodactyla, Caprinae) from the Early Pliocene of Mallorca (Balearic Islands, western Mediterranean. Geological Magazine, 147(6), 871–885.

Bover, P., Rofes, J., Bailon, S., Agusti, J., Cuenca-Bescós, G., Torres, E., & Alcover, J. A. (2014). Late Miocene/Early Pliocene vertebrate fauna from Mallorca (Balearic Islands, Western Mediterranean): An update. Integrative Zoology, 9(2), 183–196.

Breda, M., Collinge, S. E., Parfitt, S. A., & Lister, A. M. (2010). Metric analysis of ungulate mammals in the early Middle Pleistocene of Britain, in relation to taxonomy and biostratigraphy: I: Rhinocerotidae and Bovidae. Quaternary International, 228(1–2), 136–156.

Cranbrook, E. O. (2010). Late quaternary turnover of mammals in Borneo: The zooarchaeological record. Biodiversity & Conservation, 19(2), 373.

Crowley, B. E., Godfrey, L. R., Bankoff, R. J., Perry, G. H., Culleton, B. J., Kennett, D. J., Sutherland, M. R., Samonds, K. E., & Burney, D. A. (2017). Island‐wide aridity did not trigger recent megafaunal extinctions in Madagascar. Ecography, 40(8), 901–912.

Currant, A., & Jacobi, R. (2001). A formal mammalian biostratigraphy for the Late Pleistocene of Britain. Quaternary Science Reviews, 20(16–17), 1707–1716.

De Groote, I., Lewis, M., & Stringer, C. (2018). Prehistory of the British Isles: A tale of coming and going. Bulletins et Mémoires de La Société d’Anthropologie de Paris, 30(1–2), 1–13.

Denham, T., & Mountain, M. J. (2016). Resolving some chronological problems at Nombe rock shelter in the highlands of Papua New Guinea. Archaeology in Oceania, 51(S1), 73–83.

Doukas, C. S., & Athanassiou, A. (2003). Review of the Pliocene and Pleistocene proboscidea (Mammalia) from Greece. Deinsea, 9(1), 97–110.

Faure, M., Guérin, C., Genty, D., Gommery, D., & Ramanivosoa, B. (2010). Le plus ancien hippopotame fossile (Hippopotamus laloumena) de Madagascar (Belobaka, Province de Mahajanga. Comptes Rendus Palevol, 9(4), 155–162.

Flannery, T. F. (1999). The Pleistocene mammal fauna of Kelangurr Cave, central montane Irian Jaya, Indonesia. Records of the Western Australia Museum, 57, 341–350.

Gillespie, R., Camens, A. B., Worthy, T. H., Rawlence, N. J., Reid, C., Bertuch, F., Levchenko, V., & Cooper, A. (2012). Man and megafauna in Tasmania: Closing the gap. Quaternary Science Reviews, 37, 38–47.

Gommery, D., Ramanivosoa, B., Mein, P., Sénégas, F., Faure, M., & Guérin, C. (2018). Les recherches franco-malgaches à Belobaka de 2003 à 2012 (Province de Mahajanga, nord-ouest de Madagascar). Revue de Paléobiologie. https://doi.org/10.5281/zenodo.2545113

Gonzalez, S., Kitchener, A. C., & Lister, A. M. (2000). Survival of the Irish elk into the Holocene. Nature, 405(6788), 753–754.

Gruwier, B., Vos, J., & Kovarovic, K. (2015). Exploration of the taxonomy of some Pleistocene Cervini (Mammalia, Artiodactyla, Cervidae) from Java and Sumatra (Indonesia): A geometric-and linear morphometric approach. Quaternary Science Reviews, 119, 35–53.

Handa, N. (2019). Reassessment of a Pleistocene rhinocerotid (Mammalia, Perissodactyla) from Aira, Kagoshima, southwestern Japan. Paleontological Research, 23(1), 55–64.

Handa, N., & Kato, T. (2020). A Pliocene rhinocerotid (Mammalia, Perissodactyla) from Ajimu, Oita Prefecture, southwestern Japan, with comments on the Japanese Pliocene rhinocerotid fossil records. PalZ, 94(4), 759–768.

Handa, N., & Kawabe, S. (2016). Femur of Schizotheriinae (Perissodactyla, Chalicotheriidae) from the Lower Miocene Hiramaki Formation of the Mizunami Group in Gifu Prefecture, Central Japan. Journal of Vertebrate Paleontology, 36(4), 1131163.

Handa, N., & Pandolfi, L. (2016). Reassessment of the Middle Pleistocene Japanese rhinoceroses (Mammalia, Rhinocerotidae) and paleobiogeographic implications. Paleontological Research, 20(3), 247–260.

Hansford, J., Nuñez-Miño, J.M., Young, R.P., Brace, S., Brocca, J.L. and Turvey, S.T. (2012). Taxonomy-testing and the ‘Goldilocks Hypothesis’: morphometric analysis of species diversity in living and extinct Hispaniolan hutias. Systematics and Biodiversity, 10(4), pp.491-507.

Hooijer, D. A. (1945). A fossil gazelle (Gazelle schreuderae nov. Spec.) from the Netherlands. Zoologische Mededelingen, 25(9), 55–64.

Hooijer, D. A. (1964). New records of mammals from the Middle Pleistocene of Sangiran, Central Java. Zoologische Mededelingen, 40(10), 73–88.

Hooijer, D. A., & Kurth, G. (1962). The middle pleistocene fauna of Java. Gustav Fischer Verlag.

Hope, J. H. (1974). The biogeography of the mammals of the islands of Bass Strait. In Biogeography and ecology in Tasmania (pp. 397–415). Springer.

Ingicco, T., Bergh, G., Vos, J., Castro, A., Amano, N., & Bautista, A. (2016). A new species of Celebochoerus (Suidae, Mammalia) from the Philippines and the paleobiogeography of the genus Celebochoerus Hooijer, 1948. Geobios, 49(4), 285–291.

Iwase, A., Hashizume, J., Izuho, M., Takahashi, K., & Sato, H. (2012). Timing of megafaunal extinction in the late Late Pleistocene on the Japanese Archipelago. Quaternary International, 255, 114–124.

Kawamura, A., Chang, C. H., & Kawamura, Y. (2016). Middle Pleistocene to Holocene mammal faunas of the Ryukyu Islands and Taiwan: An updated review incorporating results of recent research. Quaternary International, 397, 117–135.

Kouvari, M., & Geer, A. A. E. (2018). Biogeography of extinction: The demise of insular mammals from the Late Pleistocene till today. Palaeogeography, Palaeoclimatology, Palaeoecology, 505, 295–304.

Lister, A. M. (1989). Rapid dwarfing of red deer on Jersey in the last interglacial. Nature, 342(6249), 539–542.

Lister, A. M., Parfitt, S. A., Owen, F. J., Collinge, S. E., & Breda, M. (2010). Metric analysis of ungulate mammals in the early Middle Pleistocene of Britain, in relation to taxonomy and biostratigraphy: II: Cervidae, Equidae and Suidae. Quaternary International, 228(1–2), 157–179.

Lomolino, M. V., Geer, A. A., Lyras, G. A., Palombo, M. R., Sax, D. F., & Rozzi, R. (2013). Of mice and mammoths: Generality and antiquity of the island rule. Journal of Biogeography, 40(8), 1427–1439.

Louys, J., Kealy, S., O’Connor, S., Price, G. J., Hawkins, S., Aplin, K., Rizal, Y., Zaim, J., Tanudirjo, D. A., Santoso, W. D., & Hidayah, A. R. (2017). Differential preservation of vertebrates in Southeast Asian caves. International Journal of Speleology, 46(3), 6.

Louys, J., Price, G. J., & O’Connor, S. (2016). Direct dating of Pleistocene stegodon from Timor Island, East Nusa Tenggara. PeerJ, 4, 1788.

Mackness, B. (2013). On the identity of ‘Kolopsis’ watutense (Anderson, 1937)(Diprotodontidae, Marsupialia) and the New Guinean diprotodontid radiation. Alcheringa: An Australasian Journal of Palaeontology, 37(1), 39–47.

Macphail, M. K. (1996). A habitat for the enigmatic Wynyardia bassiana Spencer, 1901, Australia’s first described Tertiary land mammal? Alcheringa, 20(3), 227–243.

MacPhee, R.D., White, J.L. and Woods, C.A. (2000). New megalonychid sloths (Phyllophaga, Xenarthra) from the Quaternary of Hispaniola. American Museum Novitates, 2000(3303), pp.1-32.

MacPhee, R. D. E. (2009). Insulae infortunatae: Establishing a chronology for Late Quaternary mammal extinctions in the West Indies. In American megafaunal extinctions at the end of the Pleistocene (pp. 169–193). Springer.

Manamendra-Arachchi, K., & Wickremasinghe, H. (2014). Action Plan for Conservation and Sustainable Use of Palaeobiodiversity in Sri Lanka. Biodiversity Secretariat, Ministry of Environment & Renewable Energy.

Marra, A. C. (2013). Evolution of endemic species, ecological interactions and geographical changes in an insular environment: A case study of Quaternary mammals of Sicily (Italy, EU. Geosciences, 3(1), 114–139.

McDowell, M. C., Prideaux, G. J., Walshe, K., Bertuch, F., & Jacobsen, G. E. (2015). Re‐evaluating the Late Quaternary fossil mammal assemblage of Seton Rockshelter, Kangaroo Island, South Australia, including the evidence for late‐surviving megafauna. Journal of Quaternary Science, 30(4), 355–364.

McFarlane, D. A., Lundberg, J., & Maincent, G. (2014). New specimens of Amblyrhiza inundata (Rodentia: Caviomorpha) from the Middle Pleistocene of Saint Barthélemy, French West Indies. Caribbean Journal of Earth Science, 47, 15–19.

McFarlane, D. A., MacPhee, R. D., & Ford, D. C. (1998). Body Size Variability and a Sangamonian Extinction Model forAmblyrhiza, a West Indian Megafaunal Rodent. Quaternary Research, 50(1), 80–89.

Melis, R. T., Palombo, M. R., Ghaleb, B., & Meloni, S. (2016). A key site for inferring the timing of dispersal of giant deer in Sardinia, the Su Fossu de Cannas cave. Sadali, Italy. Quaternary Research, 86(3), 335–347.

Mennecart, B., Zoboli, D., Costeur, L., & Pillola, G. L. (2019). On the systematic position of the oldest insular ruminant Sardomeryx oschiriensis (Mammalia, Ruminantia) and the early evolution of the Giraffomorpha. Journal of Systematic Palaeontology, 17(8), 691–704.

Menzies, J. I., & Ballard, C. (1994). Some new records of Pleistocene megafauna from New Guinea. Science in New Guinea, 20(2/3), 113–139.

Monaghan, N. T. (2017). Irish Quaternary vertebrates. In Advances in Irish Quaternary Studies (pp. 255–291). Atlantis Press.

Morgan, G. S., Macphee, R. D., Woods, R., & Turvey, S. T. (2019). Late Quaternary fossil mammals from the Cayman Islands, West Indies. Bulletin of the American Museum of Natural History, 2019(428), 1–82.

Mörs, T., & Tomida, Y. (2018). Euroxenomys nanus sp. Nov., a minute beaver (Rodentia, Castoridae) from the Early Miocene of Japan. Paleontological Research, 22(2), 145–149.

Moyà-Solà, S., Agustí, J., & Pons, J. (1984). The Mio-Pliocene insular faunas from the west Mediterranean origin and distribution factors. Paléobiologie Continentale, 14(2), 347–357.

Muhs, D. R., Simmons, K. R., Groves, L. T., McGeehin, J. P., Schumann, R. R., & Agenbroad, L. D. (2015). Late Quaternary sea-level history and the antiquity of mammoths (Mammuthus exilis and Mammuthus columbi. Quaternary Research, 83(3), 502–521.

Newsom, L. A., & Wing, E. S. (2004). On land and sea: Native American uses of biological resources in the West Indies. University of Alabama Press.

Nishioka, Y., & Ando, Y. (2016). A cervoid tooth from the lower Miocene Nakamura Formation of the Mizunami Group in Kani City, Gifu Prefecture, central Japan. Bulletin of the Mizunami Fossil Museum, 42, 39–44.

Oliver, J. R. (1995). The archaeology of Lower Camp site, Culebra Island: Understanding variability in peripheral zones. In R. E. Alegría & M. Rodríguez (Eds.), Proceedings of the XV International Congress for Caribbean Archaeology, Centro de Estudios Avanzados de Puerto Rico y el Caribe (pp. 485–500).

Otskuka, H., & Takahashi, A. (2000). Pleistocene vertebrate faunas in the Ryukyu Islands: Their migration and extinction. Tropics, 10(1), 25–40.

Palombo, M. R., Antonioli, F., Di Patti, C., Valeria, L. P., & Scarborough, M. E. (2020). Was the dwarfed Palaeoloxodon from Favignana Island the last endemic Pleistocene elephant from the western Mediterranean islands? Historical Biology, 1–19.

Palombo, M. R., & Rozzi, R. (2014). How correct is any chronological ordering of the Quaternary Sardinian mammalian assemblages? Quaternary International, 328, 136–155.

Palombo, M. R., Rozzi, R., & Bover, P. (2013). The endemic bovids from Sardinia and the Balearic Islands: State of the art. Geobios, 46(1–2), 127–142.

Pereira, E., Salotti, M., & Bonifay, M. F. (2005). A synthesis of knowledge on the large Pleistocene mammalian fauna from Corsica. Proceedings of the International Symposium ‘“Insular Vertebrate Evolution: The Paleontological Approach,”’Monografies de La Societat d’Historia Natural de Les Balears, 12, 287–292.

Pigati, J. S., Muhs, D. R., & McGeehin, J. P. (2017). On the importance of stratigraphic control for vertebrate fossil sites in Channel Islands National Park. Quaternary International, 443, 129–139.

Preece, R. C., Meijer, T., Penkman, K. E., Demarchi, B., Mayhew, D. F., & Parfitt, S. A. (2020). The palaeontology and dating of the ‘Weybourne Crag’, an important marker horizon in the Early Pleistocene of the southern North Sea basin. Quaternary Science Reviews, 236, 106177.

Prideaux, G. (2004). Systematics and evolution of the sthenurine kangaroos (Vol. 146). Univ of California Press.

Pushkina, D. (2007). The Pleistocene easternmost distribution in Eurasia of the species associated with the Eemian Palaeoloxodon antiquus assemblage. Mammal Review, 37(3), 224–245.

Puspaningrum, M. R., Bergh, G. D., Chivas, A. R., Setiabudi, E., & Kurniawan, I. (2020). Isotopic reconstruction of Proboscidean habitats and diets on Java since the Early Pleistocene: Implications for adaptation and extinction. Quaternary Science Reviews, 228, 106007.

Quintana, J., & Agustí, J. (2007). Los mamíferos insulares del Mioceno medio y superior de Menorca (islas Baleares, Mediterráneo occidental. Geobios, 40(5), 677–687.

Quintana, J., Bover, P., Alcover, J. A., Agustí, J., & Bailon, S. (2010). Presence of Hypolagus Dice, 1917 (Mammalia, Lagomorpha) in the Neogene of the Balearic Islands (Western Mediterranean): Description of Hypolagus balearicus nov. Sp. Geobios, 43(5), 555–567.

Quintana, J., Köhler, M., & Moyà-Solà, S. (2011). Nuralagus rex, gen. Et sp. Nov., an endemic insular giant rabbit from the Neogene of Minorca (Balearic Islands, Spain. Journal of Vertebrate Paleontology, 31(2), 231–240.

Quintana, J., & Moncunill-Sole, B. (2014). Hypolagus balearicus Quintana, Bover, Alcover, Agustí & Bailon, 2010 (Mammalia: Leporidae): New data from the Neogene of Eivissa (Balearic Islands, Western Mediterranean. Geodiversitas, 36(2), 283–310.

Reese, D. S. (1995). The Pleistocene vertebrate sites and fauna of Cyprus. Bulletin of the Geological Survey of Cyprus, 9, 1–203.

Rivals, F., & Lister, A. M. (2016). Dietary flexibility and niche partitioning of large herbivores through the Pleistocene of Britain. Quaternary Science Reviews, 146, 116–133.

Rozzi, R. (2017). A new extinct dwarfed buffalo from Sulawesi and the evolution of the subgenus Anoa: An interdisciplinary perspective. Quaternary Science Reviews, 157, 188–205.

Rozzi, R. (2018). Space–time patterns of body size variation in island bovids: The key role of predatory release. Journal of Biogeography, 45(5), 1196–1207.

Rozzi, R., & Palombo, M. R. (2014). Lights and shadows in the evolutionary patterns of insular bovids. Integrative Zoology, 9(2), 213–228.

Rozzi, R., Varela, S., Bover, P., & Martin, J. M. (2020). Causal explanations for the evolution of ‘low gear’locomotion in insular ruminants. Journal of Biogeography, 47(10), 2274–2285.

Rozzi, R., Winkler, D. E., De Vos, J., Schulz, E., & Palombo, M. R. (2013). The enigmatic bovid Duboisia santeng (Dubois, 1891) from the Early–Middle Pleistocene of Java: A multiproxy approach to its paleoecology. Palaeogeography, Palaeoclimatology, Palaeoecology, 377, 73–85.

Saarinen, J., Eronen, J., Fortelius, M., Seppä, H., & Lister, A. M. (2016). Patterns of diet and body mass of large ungulates from the Pleistocene of Western Europe, and their relation to vegetation. Palaeontol. Electron, 19(3), 1–58.

Salotti, M., Bailon, S., Bonifay, M.-F., Courtois, J.-Y., Ferrandini, J., Ferrandini, M., La Milza, J.-C., Mourer-Chauviré, C., Popelard, J.-B., Quinif, Y., Réal-Testud, A.-M., Miniconi, C., Pereira, E., & Persiani, C. (1997). Castiglione 3, un nouveau remplissage fossilifère d’âge Pléistocène moyen dans le karst de la région d’Oletta (Haute -Corse). C.R. Acad. Sci. Paris, II a, 67–74.

Semprebon, G. M., Rivals, F., Fahlke, J. M., Sanders, W. J., Lister, A. M., & Göhlich, U. B. (2016). Dietary reconstruction of pygmy mammoths from Santa Rosa Island of California. Quaternary International, 406, 123–136.

Sherani, S. (2021). Short notes on a second tiger (Panthera tigris) from Late Pleistocene Borneo. Historical Biology, 33(4), 463–467. https://doi.org/10.1080/08912963.2019.1625348

Símonarson, L. A., & Eiríksson, J. (2012). Steingervingar og setlög á Íslandi. Náttúrufræðingurinn, 82(1–4), 13–25.

Stefaniak, K., Stachowicz-Rybka, R., Borówka, R. K., Hrynowiecka, A., Sobczyk, A., Moskal-del Hoyo, M., Kotowski, A., Nowakowski, D., Krajcarz, M. T., Billia, E. M. E., Persico, D., Burkanova, E. M., Leshchinskiy, S. V., van Asperen, E., Ratajczak, U., Shpansky, A. V., Lempart, M., Wach, B., Niska, M., … Kovalchuk, O. (2020). Browsers, grazers or mix-feeders? Study of the diet of extinct Pleistocene Eurasian forest rhinoceros Stephanorhinus kirchbergensis (Jäger, 1839) and woolly rhinoceros Coelodonta antiquitatis (Blumenbach, 1799). Quaternary International, S1040618220305048. https://doi.org/10.1016/j.quaint.2020.08.039

Stewart, J. R. (2008). The progressive effect of the individualistic response of species to Quaternary climate change: An analysis of British mammalian faunas. Quaternary Science Reviews, 27(27–28), 2499–2508.

Storm, P., Aziz, F., Vos, J., Kosasih, D., Baskoro, S., & Hoek Ostende, L. W. (2005). Late Pleistocene Homo sapiens in a tropical rainforest fauna in East Java. Journal of Human Evolution, 49(4), 536–545.

Sutherland, F. L., & Kershaw, R. C. (1971). The Cainozoic geology of Flinders Island, Bass Strait. Papers and Proceedings of the Royal Society of Tasmania, 105, 151–178.

Sutton, A., Mountain, M. J., Aplin, K., Bulmer, S., & Denham, T. (2009). Archaeozoological records for the highlands of New Guinea: A review of current evidence. Australian Archaeology, 69(1), 41–58.

Taçon, P. S., & Webb, S. (2017). Art and megafauna in the Top End of the Northern Territory, Australia: Illusion or reality? The Archaeology of Rock Art in Western Arnhem Land, Australia, 47, 145.

Theodorou, G., Symeonidis, N., & Stathopoulou, E. (n.d.). Elephas tiliensis n. Sp. From Tilos island (Dodecanese, Greece). 42, 14.

Tomida, Y., Nakaya, H., Saegusa, H., Miyata, K., & Fukuchi, A. (2013). Miocene land mammals and stratigraphy of Japan. In X. Wang, L. J. Flynn, & M. Fortelius (Eds.), Neogene Terrestrial Mammalian Biostratigraphy and Chronology of Asia (pp. 314–333). Columbia University Press.

Trias, M., Ottenwalder, J.A. and Jaume, D. (1997). Una campaña en la República Dominicana. Resultados preliminares. Endins: publicació d'espeleologia, pp.63-74.

Turner, A. (2009). The evolution of the guild of large Carnivora of the British Isles during the Middle and Late Pleistocene. Journal of Quaternary Science: Published for the Quaternary Research Association, 24(8), 991–1005.

Turney, C. S., Flannery, T. F., Roberts, R. G., Reid, C., Fifield, L. K., Higham, T. F., Jacobs, Z., Kemp, N., Colhoun, E. A., Kalin, R. M., & Ogle, N. (2008). Late-surviving megafauna in Tasmania, Australia, implicate human involvement in their extinction. Proceedings of the National Academy of Sciences, 105(34), 12150–12153.

Turvey, S. T., Crees, J. J., Hansford, J., Jeffree, T. E., Crumpton, N., Kurniawan, I., Setiyabudi, E., Guillerme, T., Paranggarimu, U., Dosseto, A., & Van Den Bergh, G. D. (2017). Quaternary vertebrate faunas from Sumba, Indonesia: Implications for Wallacean biogeography and evolution. Proceedings of the Royal Society B: Biological Sciences, 284(1861), 20171278.

Turvey, S. T., Walsh, C., Hansford, J. P., Crees, J. J., Bielby, J., Duncan, C., Hu, K., & Hudson, M. A. (2019). Complementarity, completeness and quality of long-term faunal archives in an Asian biodiversity hotspot. Philosophical Transactions of the Royal Society B, 374(1788), 20190217.

Tweet, J. S., Santucci, V. L., Convery, K., Hoffman, J., & Kirn, L. (2020). Channel Islands National Park: Paleontological resource inventory (public version [Natural Resource Report NPS/CHIS/NRR—2020/2171.]. National Park Service, Fort Collins. https://doi.org/10.36967/nrr-2278664

Upham, N. S. (2017). Past and present of insular Caribbean mammals: Understanding Holocene extinctions to inform modern biodiversity conservation. Journal of Mammalogy, 98(4), 913–917.

Van Den Bergh, G. D., Awe, R. D., Morwood, M. J., Sutikna, T., & Saptomo, E. W. (2008). The youngest Stegodon remains in Southeast Asia from the Late Pleistocene archaeological site Liang Bua, Flores, Indonesia. Quaternary International, 182(1), 16–48.

Van der Geer, A. A. E. (2014). Parallel patterns and trends in functional structures in extinct island mammals. Integrative Zoology, 9(2), 167–182.

Van Der Geer, A. A. E. (2018). Uniformity in variety: Antler morphology and evolution in a predator-free environment. Palaeontologia Electronica, 21.

Van der Geer, A. A. E., & Drinia, H. (2016). Bears on islands: The fossil record, a re-appraisal. 21st International Cave Bear Symposium.

Van der Geer, A. A. E., Lyras, G., De Vos, J., & Dermitzakis, M. (2011). Evolution of island mammals: Adaptation and extinction of placental mammals on islands. John Wiley & Sons.

Van Weers, D. J. (2005). A taxonomic revision of the Pleistocene Hystrix (Hystricidae, Rodentia) from Eurasia with notes on the evolution of the family. Contributions to Zoology, 74(3–4), 301–312.

Vartanyan, S. L., Arslanov, K. A., Karhu, J. A., Possnert, G., & Sulerzhitsky, L. D. (2008). Collection of radiocarbon dates on the mammoths (Mammuthus primigenius) and other genera of Wrangel Island, northeast Siberia, Russia. Quaternary Research, 70(1), 51–59.

Webb, S. (2013). Corridors to extinction and the Australian megafauna. Newnes.

Woods, R., Barnes, I., Brace, S., & Turvey, S. T. (2021). Ancient DNA Suggests Single Colonization and Within-Archipelago Diversification of Caribbean Caviomorph Rodents. Molecular Biology and Evolution, 38(1), 84–95.

Yalden, D. W. (1982). When did the mammal fauna of the British Isles arrive? Mammal Review, 12(1), 1–57.

Zhao, L. X., & Zhang, L. Z. (2013). New fossil evidence and diet analysis of Gigantopithecus blacki and its distribution and extinction in South China. Quaternary International, 286, 69–74.

Zijlstra, J. S., McFarlane, D. A., Van Den Hoek Ostende, L. W., & Lundberg, J. (2014). New rodents (Cricetidae) from the N eogene of Curaçao and Bonaire, Dutch Antilles. Palaeontology, 57(5), 895–908.

Zoboli, D., & Pillola, G. L. (2017). Early Miocene insular vertebrates from Laerru (Sardinia, Italy): Preliminary note. Rivista Italiana di Paleontologia e Stratigrafia, 123, 149–158.

(N.d.). https://paleobiodb.org

### First age of appearance of large mammal herbivores on selected archipelagos

Andermann, T., Faurby, S., Turvey, S. T., Antonelli, A., & Silvestro, D. (2020). The past and future human impact on mammalian diversity. Science Advances, 6(36), eabb2313.

Carlson, L. A., & Steadman, D. W. (2009). Examining temporal differences in faunal exploitation at two ceramic age sites in Puerto Rico. The Journal of Island and Coastal Archaeology, 4(2), 207–222.

Cooke, S. B., Dávalos, L. M., Mychajliw, A. M., Turvey, S. T., & Upham, N. S. (2017). Anthropogenic extinction dominates Holocene declines of West Indian mammals. Annual Review of Ecology, Evolution, and Systematics, 48, 301–327.

Firmat, C., Rodrigues, H. G., Renaud, S., HUTTERER, R., GARCIA-TALAVERA, F., & MICHAUX, J. (2010). Mandible morphology, dental microwear, and diet of the extinct giant rats Canariomys (Rodentia: Murinae) of the Canary Islands (Spain). Biological Journal of the Linnean Society, 101(1), 28–40.

Florida Museum Vertebrate Paleontology Database. Https://www.floridamuseum.ufl.edu/vertpaleo-search/. (n.d.).

Herridge, V. L. (2010). Dwarf elephants on Mediterranean islands: A natural experiment in parallel evolution [Phdthesis]. UCL (University College London).

Hooijer, D. A. (1962). Quaternary langurs and macaques from the Malay Archipelago. Zoologische Verhandelingen, 55(1), 1–64.

Indriati, E., & Antón, S. C. (2008). Earliest Indonesian facial and dental remains from Sangiran, Java: A description of Sangiran 27. Anthropological Science, 116(3), 219–229.

MacPhee, R. D. E., & Woods, C. A. (1982). A new fossil cebine from Hispaniola. American Journal of Physical Anthropology, 58(4), 419–436.

MacPHEE, R. D., Iturralde-Vinent, M. A., & Gaffney, E. S. (2003). Domo de Zaza, an Early Miocene Vertebrate Locality in South-Central Cuba, with Notes on the Tectonic Evolution of Puerto Rico and the Mona Passage1. American Museum Novitates, 2003(3394), 1–42.

McAfee, R. K., & Beery, S. M. (2019). Intraspecific variation of Megalonychid sloths from Hispaniola and the taxonomic implications. Historical Biology.

MCFARLANE, D. A., LUNDBERG, J., & MAINCENT, G. (2014). New specimens of Amblyrhiza inundata (Rodentia: Caviomorpha) from the Middle Pleistocene of Saint Barthélemy, French West Indies. Science, 47, 15–19.

Michaux, J., López-Martínez, N., & Hernândez-Pacheco, J. (1996). A 14 C dating of Canariomys bravoi (Mammalia Rodentia), the extinct giant rat from Tenerife (Canary Islands, Spain), and the recent history of the endemic mammals in the archipelago. Vie et Milieu/Life & Environment, 261–266.

Nieves-Rivera, Á. M., Vila, J. P. Z., Hernández, C. E. F., García-Hernández, J. E., & Schizas, N. V. (n.d.). Recent and Historical Explorations of the Underwater Section of Cueva del Agua, Punta Los Ingleses, Mona Island (Puerto Rico), with a New Faunal Record1.

Orihuela, J., Viñola, L. W., Vázquez, O. J., Mychajliw, A. M., de Lara, O. H., Lorenzo, L., & Soto-Centeno, J. A. (2020). Assessing the role of humans in Greater Antillean land vertebrate extinctions: New insights from Cuba. Quaternary Science Reviews, 249, 106597.

Pregill, G. K., Steadman, D. W., & Watters, D. R. (1994). Late Quaternary vertebrate faunas of the Lesser Antilles: Historical components of Caribbean biogeography. Carnegie Museum of Natural History.

Rega, E., McFarlane, D. A., Lundberg, J., & Christenson, K. (2002). A new megalonychid sloth from the Late Wisconsinan of the Dominican Republic.

Rosenberger, A. L., Pickering, R., Green, H., Cooke, S. B., Tallman, M., Morrow, A., & Rímoli, R. (2015). 1.32$\textbackslashpm$0.11 Ma age for underwater remains constrain antiquity and longevity of the Dominican primate Antillothrix bernensis. Journal of Human Evolution, 88, 85–96.

Scarborough, M. E. (2020). Insular adaptations in the appendicular skeleton of Sicilian and Maltese dwarf elephants.

Turvey, S. T., Grady, F. V., & Rye, P. (2006). A new genus and species of ‘giant hutia’(Tainotherium valei) from the Quaternary of Puerto Rico: An extinct arboreal quadruped? Journal of Zoology, 270(4), 585–594.

Van den Bergh, G. D. (1999). The Late Neogene elephantoid-bearing faunas of Indonesia and their palaeozoogeographic implications. Scripta Geologica, 117, 1–419.

van den Bergh, M., G. D. ,. Prasetyo, U. ,. Setiyabudi, E. ,. Puspaningrum, M. ,. Kurniawan, ,. I. ,. Storey. (2019). The Early Pleistocene terrestrial vertebrate faunal sequence of Java, Indonesia. 210.

van der Geer, A. A. E., Lyras, G. A., & Volmer, R. (2018). Insular dwarfism in canids on Java (Indonesia) and its implication for the environment of Homo erectus during the Early and earliest Middle Pleistocene. Palaeogeography, Palaeoclimatology, Palaeoecology, 507, 168–179. https://doi.org/10.1016/j.palaeo.2018.07.009

Westaway, K. E., Morwood, M. J., Roberts, R. G., Rokus, A. D., Zhao, J.-X., Storm, P., Aziz, F., Van den Bergh, G., Hadi, P., & De Vos, J. (2007). Age and biostratigraphic significance of the Punung Rainforest Fauna, East Java, Indonesia, and implications for Pongo and Homo. Journal of Human Evolution, 53(6), 709–717.
