## SUpplementary Information Section 5 for "Plant longevity, drought and island isolation favoured rampant evolutionary transitions towards insular woodiness"

Supplementary methods

### Table of content

### 1. Identification of insular woody species

Due to the large heterogeneity in methods of phylogenetic reconstruction and dating, as well as in data availability in the literature, we identified insular woody species in angiosperms by visually tracing character evolution on published phylogenies following a maximum parsimony approach. We consider this approach conservative, since it allowed us to adapt the interpretation to the detail and quality of the phylogenetic trees, and to resolve issues using expert judgment. Furthermore, underlying comparative sequence data for IWS at larger taxonomic scales were not available for phylogenetic studies rendering a standardized meta-analysis impossible.

To use information on species habitat, we recorded habitat information where available from the literature source and then standardized these free-text habitat descriptions into distinct categories: “forest” (including rainforest, dry forest, mesic forest and deciduous forest), “desert” (including deserts and semi-deserts), “halophytic” (including saline habitats and habitats in the marine splash water zone), “bog” (including bogs, swamps, marshy, flooded and waterlogged soils), “xerophytic” (including undefined dry vegetation), “shrubland” (including shrublands, heath and scrub), “grassland” (all grass dominated systems including grassland, meadows and steppe), “savanna” (including savanna and woodland), “paramo”, “alpine” and “cliffs, rocky outcrops & sandy or gravely soil” (all habitats dominated by bare rock, including cliffs, rocky outcrops, bare rock, gravel and sandy soils). Species could occur in multiple habitats. We then classified these detailed habitat types into forest (forests) vs. open habitats (all others).

### 2. Insular woodiness on the angiosperm Tree of Life

For the calculation of phylogenetic signal using Pagel’s lambda and Blomberg’s K as described in the main manuscript and for the visualization in Figure 1 of the main manuscript, we calculated the proportion of insular woody species per genus and family. To do so, we approximated the total number of plant species in a genus and family, using the Leipzig Catalogue of Vascular Plants^1^.

To identify the age of shifts towards IW, we compiled the stem ages of those insular woody lineages for which phylogenetic studies were available (Supplementary Information Section 4 for full literature list). When multiple age estimates were available for the same clade, we only retained the most recent estimates.

### 4 Potential drivers of insular woodiness

#### 4.1 Environmental correlates of the number of insular woody species per island

We compiled data on island environmental characteristics from various sources including several levels of quality check. See the scripts 01-09 available in a zenodo repository upon publication for the R code used to obtain and prepare the data, scripts 10-13 for figure preparation and scripts 14-18 for data analysis.

Since the number of insular woody species on islands may be a result of the total angiosperm diversity, we compiled estimates of total angiosperm species number. To do so, we downloaded checklists of angiosperms from the Global Inventory of Floras and Traits database (GIFT)^2^. GIFT compiles plant checklists from a variety of sources and in varying levels of completeness. We obtained checklists of flowering plants from islands worldwide at GIFT, restricting our search to references including type_ref 1:11 and ref_included 1,2,3,4, and only complete island floras. We only retained species indicated as “native” for each island. Furthermore, we used the taxonomic backbone of GIFT to identify monocot species based on order name, considering species of the orders Arecales, Acorales, Alismatales, Asparagales, Commeliniales, Dasypogonales, Dioscoreales, Liliales, Pandanales, Petrosaviales, Poales, and Zingiberales as monocots. For few islands for which no total angiosperm richness estimates were available from GIFT but from other sources, we added approximations of total angiosperm species richness, specifically the islands of Borneo^3^, Madagascar (Peter Phillipson, pers communication), New Guinea^4^, and Waya^5^.

#### 4.2 Environmental correlates of the number of evolutionary transitions to insular woodiness per archipelago.

We scored the number of evolutionary shifts towards IW per archipelago based on the literature and the accompanying phylogenetic literature. We consider a shift on an archipelago, if it was possible to assign it unambiguously to this archipelago based on a biogeographic analysis of a sister-clade comparison based on a phylogenetic tree from the literature. We then used the environmental data compiled at the island-level to generate archipelago-level information on the absolute latitude of the archipelago centroid, the minimum distance to the closest mainland (as the minimum distance from the closest islands of the archipelago), the minimum age of the archipelago (as the oldest estimated age of the any individual island in the archipelago), and the total archipelago area (as the sum of the land area of all individual islands). We only included archipelagos with at least one shift to IW or data available on all environmental predictors. Since the data on the minimum archipelago age were missing for five archipelagos with at least one IW shift, we imputed island age for these archipelagos using multiple imputation using random forests implemented in the missForest package.

We tested for predictor collinearity using the variance inflation factor (<2 for all predictors). We then transformed island age, island area and distance to the next continent (log(1+x)) and scaled all predictors to zero mean and standard variance. Due to the large number of islands with zero shifts to IW, we chose a hurdle count model developed for regression analyses of zero-inflated count data, as implemented in the ‘pscl’ R package^6^. The hurdle model consists of two components: a truncated Poisson generalized linear model with log link modelling counts above zero (i.e. the number of shifts towards insular woodiness if any occur) and a hurdle component modelling zero vs larger counts (i.e. on which islands do IW shifts occur at all) using a binomial model and a logit link. We fitted four different models; one main model and three additional models for sensitivity analysis

I. main model: n_shifts_ ~ log(area) + log(distance) + absolute latitude + log(age); excluding Hawaii and the Canary island. We removed the Canary islands and Hawaii as outliers from the main model, since we expect them to overly influence the model as high leverage outliers due to their very high numbers of shifts.
II. same as (I) including the Hawaiian and Canary islands,
III. same as (I) but excluding archipelago age as predictor and thereby adding another 15 archipelagos as data points (all without evolutionary shifts), and
IV. same as (III) and adding Hawaii and the Canary Islands.

### 5.1 Software used

We performed all analyses in R^7^, using the ‘ape’ 5.5^8,9^, ‘car’ 3.0-11^10^, ‘cowplot’ 1.1.1^11^, ‘deeptime’ 0.1.0^12^, ‘dssatr’ 0.1.0.9000^13^, ‘elevatr’ 0.4.1^14^, ‘fasterize’ 1.0.3^15^, ‘forcats’ 0.5.1^16^, ‘ggnewscale’ 0.4.5^17^, ‘ggrtree’ 3.0.2^18,19^, ‘gridExtra’ 2.3^20^, ‘LCVP’ 1.0.4^1,21^, ‘lcvplants’ 2.0^22^, ‘missForest’ 1.4.0^23,24^, ‘olsrr’ 0.5.3^25^, ‘phytools’ 0.7-80^26,27^, ‘picante’ 1.8.2^28^, ‘raster’ 3.4-13^29^, ‘readxl’ 1.3.1^30^, ‘rnaturalearth’ 0.1.0^31^, ‘sf’ 1.0-2^32^, ‘sp’ 1.4-5^33,34^, ‘tidyverse’ 1.3.0^35,36^, ‘viridis’ 0.6.1^37^, ‘wesanderson’ 0.3.6^38^, ‘writexl’ 1.4.0^39^ packages.

### 6. Supplementary references

1.	Freiberg, M. *et al.* LCVP, The Leipzig catalogue of vascular plants, a new taxonomic reference list for all known vascular plants. *Sci. Data* 416 (2020) doi:10.1038/s41597-020-00702-z.

2.	Weigelt, P., König, C. & Kreft, H. GIFT – A Global Inventory of Floras and Traits for macroecology and biogeography. *J. Biogeogr.* **47**, 16–43 (2020).

3.	Biodiversity of Borneo. *Wikipedia* (2021).

4.	Cámara-Leret, R. *et al.* New Guinea has the world’s richest island flora. *Nature* **584**, 579–583 (2020).

5.	Gardner, R. The plants of Waya Island, Fij. *Rec. Auckl. Mus.* **53**, 43–76 (2018).

6.	Zeileis, A., Kleiber, C. & Jackman, S. Regression Models for Count Data in R. *J. Stat. Softw.* **27**, 1–25 (2008).

7.	R Core Team. *R: A Language and environment for statistical computing*. (R Foundation for Statistical Computing, 2021).

8.	Paradis, E. *et al.* *ape: Analyses of Phylogenetics and Evolution*. (2021).

9.	Paradis, E. & Schliep, K. ape 5.0: an environment for modern phylogenetics and evolutionary analyses in R. *Bioinformatics* **35**, 526–528 (2019).

10.	Fox, J., Weisberg, S. & Price, B. *car: Companion to Applied Regression*. (2021).

11.	Wilke, C. O. *cowplot: Streamlined Plot Theme and Plot Annotations for ggplot2*. (2020).

12.	Gearty, W. *deeptime: Plotting Tools for Anyone Working in Deep Time*. (2021).

13.	Bocinsky, K. *dssatr: R Bindings for the Decision Support System for Agrotechnology Transfer (DSSAT) Crop Simulation System*. (2016).

14.	Hollister, J. *elevatr: Access Elevation Data from Various APIs*. (2021).

15.	Ross, N. *fasterize: Fast Polygon to Raster Conversion*. (2020).

16.	Wickham, H. *forcats: Tools for Working with Categorical Variables (Factors)*. (2021).

17.	Campitelli, E. *ggnewscale: Multiple Fill and Colour Scales in ggplot2*. (2021).

18.	Yu, G. Using ggtree to Visualize Data on Tree-Like Structures. *Curr. Protoc. Bioinforma.* **69**, e96 (2020).

19.	Yu, G., Lam, T. T.-Y. & Xu, S. *ggtree: an R package for visualization of tree and annotation data*. (2021).

20.	Auguie, B. & Antonov, A. *gridExtra: Miscellaneous Functions for ‘Grid’ Graphics*. (2017).

21.	Gentile, A., Freiberg, M., Winter, M. & Zizka, A. *LCVP: The Leipzig Catalogue of Vascular Plants (LCVP) - datapackage*. (2021).

22.	Vilela, B., Gentile, A., Freiberg, M. & Winter, M. *lcvplants: The Leipzig Catalogue of Vascular Plants (LCVP) - An Improved Taxonomic Reference List for All Known Vascular Plants*. (2021).

23.	Stekhoven, D. J. & Buehlmann, P. MissForest - non-parametric missing value imputation for mixed-type data. *Bioinformatics* **28**, 112–118 (2012).

24.	Stekhoven, D. J. *missForest: Nonparametric Missing Value Imputation using Random Forest*. (2013).

25.	Hebbali, A. *olsrr: Tools for Building OLS Regression Models*. (2020).

26.	Revell, L. J. phytools: an R package for phylogenetic comparative biology (and other things). *Methods Ecol. Evol.* **3**, 217–223 (2012).

27.	Revell, L. J. *phytools: Phylogenetic Tools for Comparative Biology (and Other Things)*. (2021).

28.	Kembel, S. W. *et al.* Picante: R tools for integrating phylogenies and ecology. *Bioinformatics* **26**, 1463–1464 (2010).

29.	Hijmans, R. J. *raster: Geographic Data Analysis and Modeling*. (2021).

30.	Wickham, H. & Bryan, J. *readxl: Read Excel Files*. (2019).

31.	South, A. *rnaturalearth: World Map Data from Natural Earth*. (2017).

32.	Pebesma, E. *sf: Simple Features for R*. (2021).

33.	Pebesma, E. J. & Bivand, R. S. Classes and methods for spatial Data: the sp Package. *R News* **5**, 21–41 (2005).

34.	Pebesma, E. & Bivand, R. *sp: Classes and Methods for Spatial Data*. (2021).

35.	Wickham, H. *et al.* Welcome to the tidyverse. *J. Open Source Softw.* **4**, 1686 (2019).

36.	Wickham, H. *tidyverse: Easily Install and Load the Tidyverse*. (2019).

37.	Garnier, S. *viridis: Colorblind-Friendly Color Maps for R*. (2021).

38.	Ram, K. & Wickham, H. *wesanderson: A Wes Anderson Palette Generator*. (2018).

39.	Ooms, J. *writexl: Export Data Frames to Excel xlsx Format*. (2021).
